# Oral RNAi of *diap1* in a pest results in rapid reduction of crop damage

**DOI:** 10.1101/737643

**Authors:** Yasuhiko Chikami, Haruka Kawaguchi, Takamasa Suzuki, Hirofumi Yoshioka, Yutaka Sato, Toshinobu Yaginuma, Teruyuki Niimi

## Abstract

Selecting an appropriate target gene is critical to the success of feeding RNA interference (f-RNAi)-based pest control. Gene targets have been chosen based on their ability to induce lethality. However, lethality induction by f-RNAi is slow-acting and crop damage can progress during this time. Here, we show that f-RNAi of *death-associated inhibitor of apoptosis protein 1* (*diap1*), but not two conventional targets *vacuolar ATPase subunit A* and *E*, induces acute feeding cessation in the solanaceous pest, *Henosepilachna vigintioctopunctata* during 24–48 hours. We also found that the feeding cessation by *diap1* f-RNAi has species-specificity and occurs with only 1.6 ng dsRNA. Our results suggest that *diap1* is an appropriate target in the context of rapid reduction of crop damage. We propose that acute feeding disorder should be assessed as a novel criterion for selecting appropriate target genes for RNAi-based pest control in addition to the conventional criterion based on lethality.

## Introduction

Techniques for manipulating endogenous essential genes in pests using RNA interference (RNAi) have become popular as alternative strategies to conventional chemical pesticides for use in pest management in the last decade^1,2,3^.

Innovative and practical applications of the technology are currently being developed, such as the use of transgenic plants that produce large amounts of double-stranded RNA (dsRNA) in the chloroplasts^4^ or environmental RNAi applications, such as spraying dsRNA^5^ on crops. Regardless of application method, the selection of an appropriate target gene is essential to achieve effective pest control^6^. The most target genes reported so far are housekeeping genes such as the *v-ATPase* genes and metabolic genes such as the chitin synthase genes (representative targets listed in Baum et al.^2^).

Conventionally, the criterion for selecting target genes is a combination of induction of lethality or growth-inhibition by gene silencing with RNAi^6^. For example, silencing of *v-ATPase subunit A* (*v-ATPase A*) or *v-ATPase subunit E* (*v-ATPase E*) by oral delivery of the dsRNA causes increased mortality in various pests of the order Coleoptera^7,8,9^, Lepidoptera^7,10^, Diptera^7^, Hemiptera^11,12^ and Orthoptera^13^. Such induction of lethality or growth-inhibition is extremely effective in terms of pest reduction and eradication. However neither lethality nor growth-inhibition are phenotype that are quick to induce, and it takes a certain amount of time before maximal effect of treatment is achieved. During the long-time span between treatment and phenotype, insect pests can continue to damage crops. For example, it takes more than a week for the RNAi effect of *v-ATPase A* or *v-ATPase E* to occur. Silencing of *v-ATPase E* requires 9 days to induce 100% mortality through injected and 25 days to induce 100% mortality through oral RNAi in *Tribolium castaneum*^14^. Therefore, in order to achieve effective RNAi-based control of herbivorous pests, it would be desirable to evaluate and search for target genes from the viewpoint of rapid termination of crop damage in addition to the induction of lethality and growth-inhibition.

In this regard, we found that the gene silencing of *death-associated inhibitor of apoptosis protein 1* (*diap1*) gene by oral-feeding RNAi (f-RNAi) causes acute feeding disorder. *diap1* is an insect homolog of the *iap* genes and is known as a suppressor of apoptosis in the fruit fly, *Drosophila melanogaster*^15^. Diap1 has E3 ubiquitin-ligase activity^16^ and strongly suppresses apoptosis via ubiquitination and degradation of the caspase protein^17^. Silencing of the *diap1* gene promotes apoptosis activity, leading to lethality. In addition, the Diap1 protein has an extremely short half-life (c.a. 40 min) in *Drosophila* S2 cells^18^. In general, short protein half-life is one of the keys for the success of RNAi targeting protein-coding genes^19^. Therefore, RNAi of the *diap1* gene is expected to have a powerful effect. In some insects, it is known that *diap1* mRNA highly expresses in the midgut^20^ (Fig. S1 in *D. melanogaster*). For these reasons, *diap1* is one of the candidate target gene for RNAi pesticides and has been investigated in various pests. So far, RNAi of the *diap1* gene by injecting the dsRNA into the hemocoel of pest has been shown to induce over 70% mortality in *T. castaneum* (Coleoptera)^14^, *Anoplophora glabripennis* (Coleoptera)^20^, *Musca domestica* (Diptera)^21^, *Delia radicum* (Diptera)^21^, *Heliothis virescens* (Lepidoptera)^22^, *Lygus lineolaris* (Hemiptera)^23^ and *Apolygus lucorum* (Hemiptera)^24^. Diap1 silencing by f-RNAi has also been reported to induce 30– 78% of mortality in *Agrilus planipennis*^24^, *T. castaneum*^14^, and *Diaphorina citri*^25^. In these reports, the maximum lethal effect of *diap1* silencing by f-RNAi takes 5–30 days. This lethal effect of *diap1* by f-RNAi is also dose-dependent. For example, in *A. planipennis*, the mortality of *diap1* f-RNAi using 1, 6 and 10 µg/µl dsRNA is 30, 35 and 78%, respectively^24^. In *M. domestica* and *D. radicum*, it was reported that *diap1* gene silencing by injection of the dsRNA induced lethality but the f-RNAi assay did not^21^. The comparative f-RNAi assay of *diap1* and *v-ATPase E* using *T. castaneum* shows that silencing of *diap1* causes less mortality than that of *v-ATPase E*^14^. Although there are differences depending on the species, it seems that lethal effects of *diap1* f-RNA require a large amount of dsRNA and relatively long time before the effect appears. In the present study, we evaluated a novel effect of the *diap1* gene as a target of f-RNAi-based pest control. Our work investigated whether *diap1* rapidly induce cessation of feeding in addition to the conventional criterion based on lethality.

In this dsRNA feeding assay, we used the 28-spotted ladybird, *Henosepilachna vigintioctopunctata*, which is a representative solanaceous pest in Asia (Fig. S2). This species eats solanaceous plants in both the larval and adult stages. The larvae molts about once every 3–5 days and pupates at the fourth molting. In this study, we fed *diap1* dsRNA to 3rd instar larvae, which are easy to bioassay in terms of body size and amount of feeding. We also compared the effects of *diap1* f-RNAi on feeding disorder with *v-ATPase A* and *v-ATPase E*. For the effect of *diap1* f-RNAi, we also evaluated the species-specificity and dose-sensitivity, which are generally challenges to select target genes for RNAi-based pest control.

## Results

### Identification and expression of target genes in *H. vigintioctopunctata*

We isolated the homologous sequences of three target genes, *diap1, v-ATPase A* and *v-ATPase E*, from cDNA of *H. vigintioctopunctata* (Fig. S3). The *diap1* homologous sequence of *H. vigintioctopunctata* was 2012 bp and 391 amino acids. Similar to the report in *D. melanogaster*^17^, the Diap1 homologous sequences had three domains, two baculoviral inhibitor of apoptosis protein repeats (BIR) and one really interesting new gene (RING) domain. The *v-ATPase A* and *v-ATPase E* homologous sequence of *H. vigintioctopunctata* had 2503 bp and 1282 bp, respectively. The V-ATPase A and V-ATPase E homologous sequence had 614 and 226 amino acids, respectively. We cannot detect any in-frame isoforms in these target genes.

To confirm the orthology of these genes, we performed molecular phylogenetic analysis for amino acid sequence of each target gene. All of these genes were located in the robust supported clade of each gene (Fig. S4). Therefore, the isolated homologous sequences were identified as the ortholog of each target gene in *H. vigintioctopunctata*. To observe the expression profile of each target gene, we performed RT-PCR using cDNA from the nerve cord, legs, wing disc, fat body, Malpighian tube, gut and carcass of *H. vigintioctopunctata* larvae. We confirmed that the *diap1, v-ATPase A* and *v-ATPase E* orthologs were transcribed in whole larval body including the gut (Fig. 1a).

**Fig. 1.**
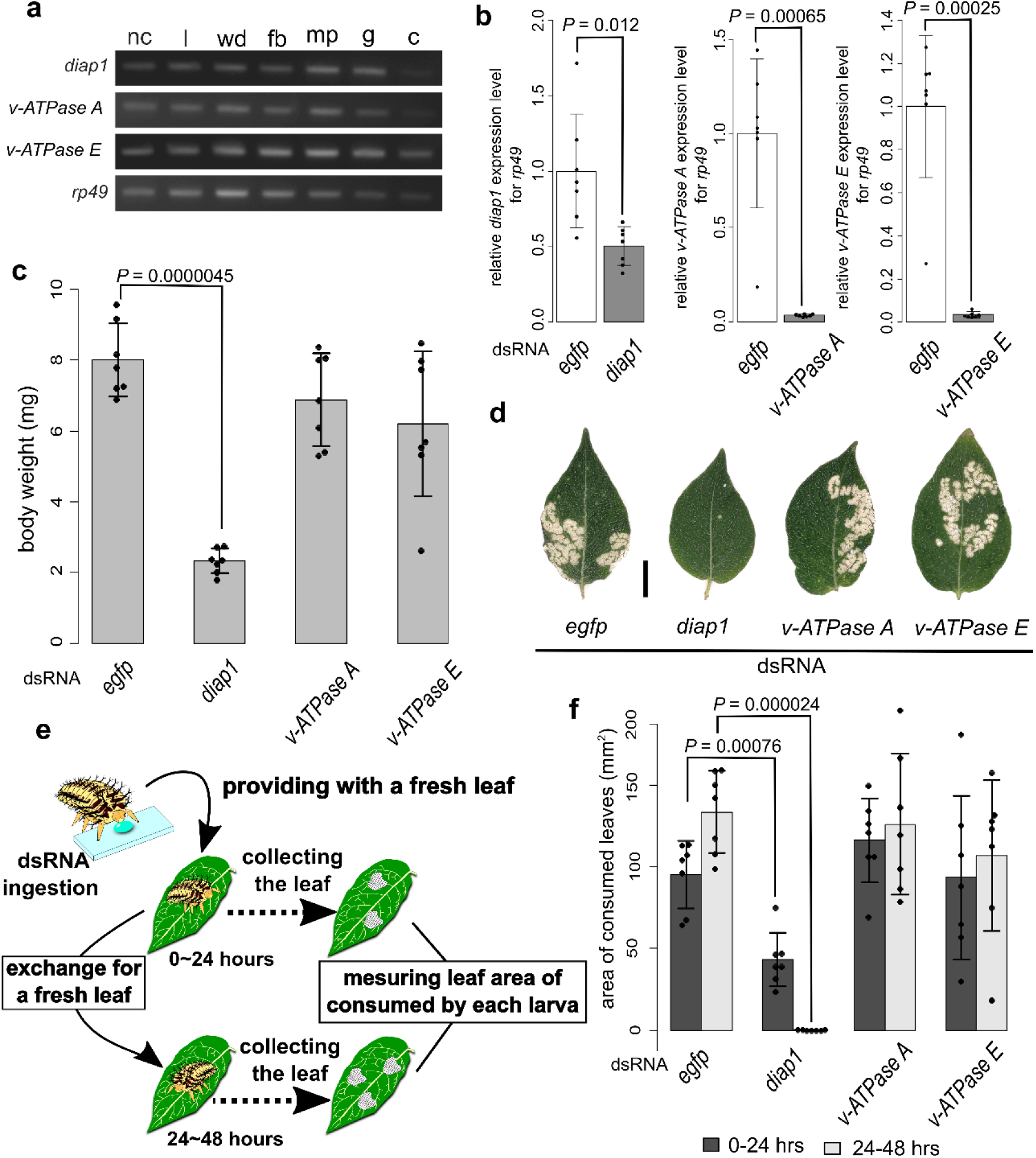
Effects of f-RNAi of *diap1, v-ATPase A* and *v-ATPase E* in *H. vigintioctopunctata*. **a** Tissue expression profile in 4th instar larva. nc, nerve cord; l, legs; wd, wing discs; fb, fatbody; mp, Malpighian tube; g, gut; c, carcass. **b** qRT-PCR in the larvae at two-days after f-RNAi. **c** Body-weight of the larvae at two-days after f-RNAi. **d** The leaves consumed by larvae two-days after f-RNAi. Scale, 10 mm. **e** Scheme of f-RNAi experiment to evaluate feeding disorders. **f** Leaf area consumed by the larvae for 0-24 hours (dark gray) and 24-48 hours (right gray) after f-RNAi. *P*-values were calculated by Welch’s *t*-test and adjusted by Holm’s method for multiple comparisons. Adjusted *P*-values were shown in c and f, and non-significances were not shown. Data of b, c and f are mean ± s.d., *n*=7 individual larvae for each f-RNAi treatment (black dot). Information for c and f are provided in Table S1.

### Knockdown effect of f-RNAi of target genes in *H. vigintioctopunctata*

For the oral feeding RNA interference (f-RNAi) assay, we designed and synthesized the 300–400 bp double-stranded RNA (dsRNA) targeting each gene (yellow lines in Supplementary Sequence). We fed 3rd instar larvae with a droplet containing 50 ng of the dsRNA (Fig. S5, *n* = 7 larvae for each gene). As the negative control, we fed 3rd instar larvae with the same dose of *egfp* dsRNA. To evaluate the knockdown effect of each gene, we performed quantitative (q) RT-PCR using cDNA of the whole-body larvae at two-days after f-RNAi treatment. The qRT-PCR analysis revealed that the expression level of each target gene in the f-RNAi treated larvae significantly decreased as compared to the control larvae (Welch’s *t* test, *P* = 0.01, 0.0007 and 0.0002 in *diap1, v-ATPase A* and *v-ATPase E*, respectively) (Fig. 1b). Compared to the control larvae, the mean expression level of *diap1, v-ATPase A* and *v-ATPase E* in the f-RNAi treated larvae reduced to 50.2, 3.6, 3.4%, respectively. The knockdown effect of *diap1* f-RNAi seems to be lower than that of *v-ATPase* genes, although, the reduced rate of *diap1* mRNA level is nearly equivalent to that in previous *diap1* f-RNAi studies^14,20,22,25^ showing the obvious phenotype. Considering these facts, we determined that our knockdown effect of *diap1* is reasonable. Therefore, our results indicate that the knockdown of each target gene by f-RNAi was effective in *H. vigintioctopunctata*.

### Acute feeding disorder effect of *diap1* silencing in *H. vigintioctopunctata*

To observe the f-RNAi effect in *H. vigintioctopunctata*, we provided the 3rd instar larvae with 50 ng (100 ng/µl) of the *diap1, v-ATPase A* or *v-ATPase E* dsRNA (*n* = 7 in each target gene). Then, we provided the treated larvae with potato leaves as food.

At 2 days after f-RNAi treatment, we found that the body weight significantly decreased in the *diap1* f-RNAi treated larvae compared to the control larvae fed with *egfp* dsRNA (Welch’s *t* test, *P* < 0.001) (Fig. 1c and Table S1). The mean body weight of the *diap1* f-RNAi larvae was 2.32 mg, which was 3.5-fold lower than the control larvae. In contrast to the *diap1* f-RNAi treatment, we cannot detect significant reduction of body weight in *v-ATPase* genes treated larvae compared to the control larvae (Fig. 1c and Table S1). We further found that the consumed area of potato leave was reduced in *diap1* f-RNAi treatment (Fig. 1d). These results suggested that silencing of the *diap1* gene but not the *v-ATPase* genes could cause feeding disorder in the *H. vigintioctopunctata* larvae.

Therefore, in order to evaluate the influence on crop damage after RNAi treatment, we measured the area of potato leaves consumed by the larvae during 0-24 and 24-48 hours after dsRNA ingestion (Fig. 1e). As the result, the *diap1* f-RNAi treated larvae showed a significantly narrower area of consumed leaves than the control larvae within 24 hours after dsRNA ingestion (mean area: 43.24 mm^2^ in *diap1* f-RNAi and 95.54 mm^2^ in *egfp* f-RNAi, Welch’s *t* test, *P* < 0.001) (Fig. 1f and Tables S1, S2). The mean leaf area consumed by the *diap1* f-RNAi treated larvae was 2.2-fold narrower than the *egfp* f-RNAi treated larvae. Surprisingly, we detected almost complete cessation of crop damage by the *diap1* f-RNAi treated larvae during 24–48 hours after dsRNA ingestion (mean area: 0.15 mm^2^ in *diap1* f-RNAi and 133.90 mm^2^ in *egfp* f-RNAi, Welch’s *t* test, *P* < 0.001) (Fig. 1f and Tables S1, S2). In contrast to the *diap1* f-RNAi, we cannot observe any significant effects on the consumed leaf area in the *v-ATPase A* and *v-ATPase E* RNAi treatment during both 0–24 and 24–48 hours (Fig. 1f and Tables S1, S2). In *egfp, v-ATPase A* and *v-ATPase E* f-RNAi treatment, the consumed leaf area increased to 108–140% during 24–48 hours compared to 0–24 hours. In contrast, the leaf area consumed by the *diap1* f-RNAi treated larvae decreased to 0.35% during 24–48 hours compared to 0–24 hours. All f-RNAi larvae did not show lethality within 48 hours in our experiments. These results showed that the f-RNAi of *diap1* but not *v-ATPase* genes induces acute feeding disorder in the larvae within 24 hours.

### Species-specificity of *diap1* f-RNAi effect in *H. vigintioctopunctata*

Species-specificity is one of the advantages of RNAi-based pest control compared to conventional chemical pesticides. However, since evolutionary and widely conserved genes generally have an overall strongly conserved sequence, there is a concern that it is difficult to design a species-specific dsRNA. Therefore, we evaluated species-specificity in the acute feeding disorder effect by *diap1* silencing.

We compared the nucleotide sequences of *diap1* among 10 insect species belonging to 7 orders of Zygentoma, Orthoptera, Hemiptera, Hymenoptera, Coleoptera, Lepidoptera and Diptera. It showed that the *diap1* sequence had 37.3-53.7% identities among the insects (Fig. 2a). The identity of *diap1* was lower than that of the *v-ATPase* genes in all comparative combinations except for the comparisons with *Locusta migratoria* (Figs. 2a, S6). Notably, the sequences between *H. vigintioctopunctata* and *Harmonia axyridis* belonging to the same family (Coccinellidae) showed only 53.65% identity in *diap1*, which was about 1.5-fold lower than those of *v-ATPase A* and *v-ATPase E* (Figs. 2a, S6). This result showed that *diap1* gene has a moderately diverged sequence compared to *v-ATPase* genes, and suggested that dsRNA targeting the homologous region in *diap1* is usable for species-specific RNAi pesticide reagents.

**Fig. 2.**
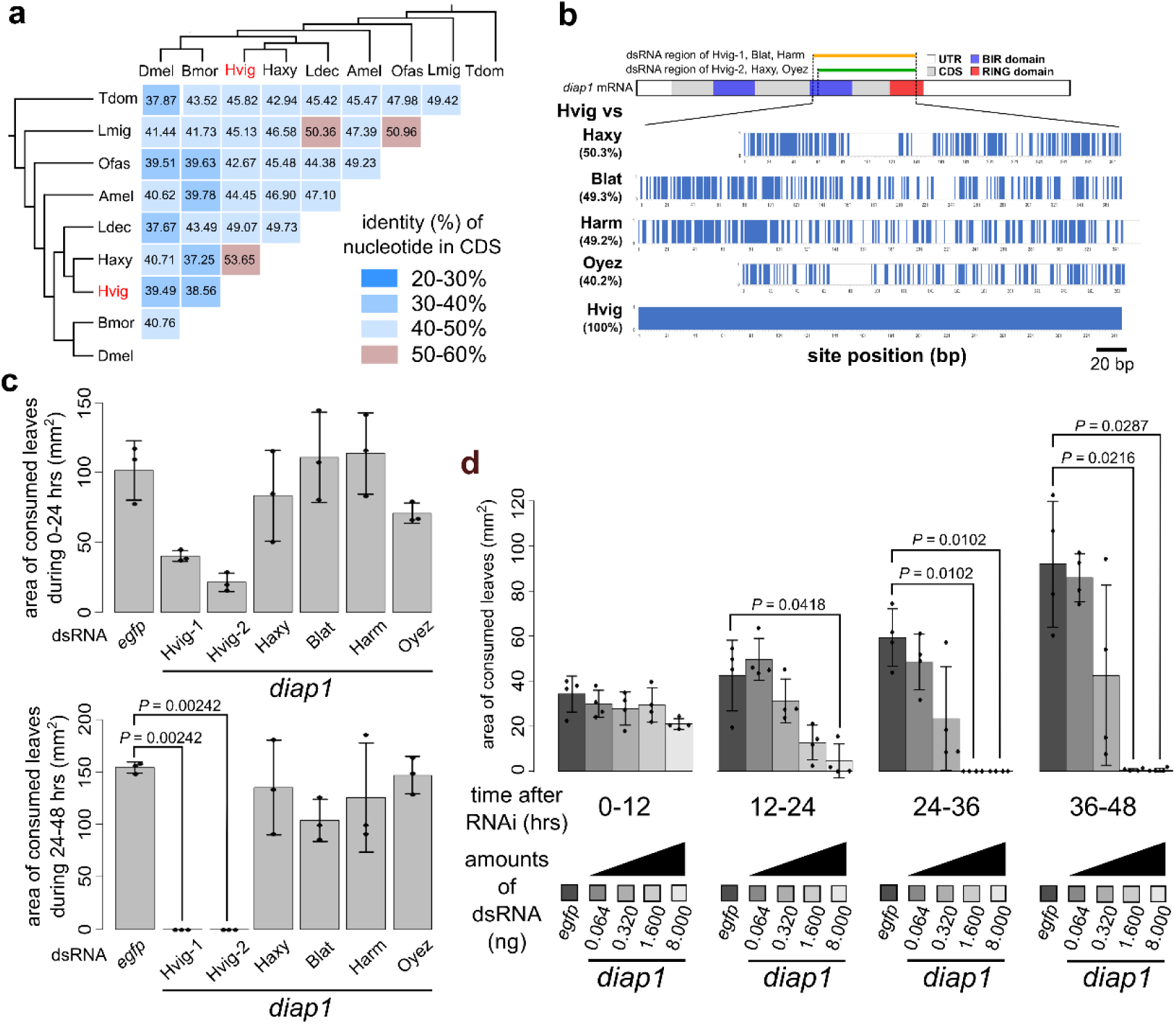
Species-specificity and efficiency of *diap1* dsRNA. **a** Identity of *diap1* nucleotide among insects. Numbers in each column indicates the identity between species. Branches show the phylogenetic relationship^33^. **b** Image of sequence identity of *diap1* dsRNAs among insects. Blue bars indicate the identical nucleotide sites between *H. vigintioctopunctata* and each species. UTR, untranslated region; CDS, coding sequence; BIR, baculovirus inhibitior of apoptosis protein repeat; RING, really interesting new gene. **c** Area consumed by the larvae during 12-24 (upper) and 24-48 (lower) hours after various insect-derived *diap1* dsRNAs ingestion. **d** Area consumed by the *diap1* f-RNAi larvae treated with different amounts of dsRNA. Data of c and d are mean ± s.d., *n*=3 in c and *n*=4 in d (black dot). *P*-values were calculated by Welch’s *t*-test and adjusted by Holm’s method. Adjusted *P*-values were shown in c and d, and non-significances were not shown. Amel, *Apis melifera*; Blat, *Blatta lateraris*; Bmor, *Bombyx mori*; Caqu, *Catajapyx aquilonaris*; Dmel, *Drosophila melanogaster*; Hvig, *Henosepilachna vigintioctopunctata*; Harm, *Helicoverpa armigera*; Haxy, *Harmonia axyridis*; Ldec, *Leptinotarsa decemlineata*; Lmig, *Locusta migratoria*; Ofas, *Oncopeltus fasciatus*; Oyez, *Oxya yezoensis*. Information for a, c and d is provided in Tables S3, 4 and 5, respectively.

Then, we validated species-specificity of *diap1* f-RNAi effect on *H. vigintioctopunctata* larvae using 320 ng dsRNAs targeting *diap1* of 4 other insects, *H. axyridis* (Coleoptera), *Blatta lateraris* (Blattodea), *Helicoverpa armigera* (Lepidoptera), *Oxya yezoensis* (Orthoptera) (*n* = 3 in each treatment) (Fig. 2b). The dsRNA region of *diap1* of *H. vigintioctopunctata* identified 50.3, 49.1, 49.3 and 40.1% with *H. axylidis, B. lateraris, H. armigera* and *O. yezoensis*, respectively. This region did not show consecutive matches over 16 bp between *H. vigintioctopunctata* and each insect. We detected that the *diap1* f-RNAi using dsRNA of *H. axyridis* and *O. yezoensis* caused weak feeding disorder effect in *H. vigintioctopunctata* larvae during 0–24 hours after the treatment (Fig. 2c). The mean leaf area during 0–24 hours after f-RNAi treatment was 2.5, 4.8, 1.2 and 1.4 -folds lower than the *egfp* f-RNAi larvae in *H. vigintioctopunctata* region 1, *H. vigintioctopunctata* region 2, *H. axyridis* and *O. yezoensis* dsRNA treatment, respectively. The *diap1* f-RNAi also showed the slightly lower consumed leaf area in all f-RNAi treatments with *diap1* dsRNA of the other species during 24–48 hours after treatment (Fig. 2c). In contrast, of the f-RNAi treatments, only *H. vigintioctopunctata diap1* dsRNA treatment showed complete feeding cessation during 24-48 hours after the treatment (Fig. 2c). We also detected the significant reduction of the consumed leaf area between 0–24 and 24–48 hours after f-RNAi treatment in only *H. vigintioctopunctata diap1* dsRNA treatment (Welch’s *t* test, *P* = 0.003 and 0.03 in region 1 and 2, respectively). The *B. laterais diap1* dsRNA treatment showed the reduction of the consumed area between 0–24 and 24–48 hours after the treatment, although, the reduction was only 6.3% in rate and did not show significant difference (Welch’s *t* test, *P* = 0.8) (Table S5). These results show that the *diap1* dsRNA used here cause weak cross-species effect in the context of reduction of leaf consuming but not in the context of the feeding cessation and inhibition of increase in appetite of *H. vigintioctopunctata* larvae.

### Dose-sensitivity of *diap1* f-RNAi effect in *H. vigintioctopunctata*

Dosage of dsRNA is one of the key factors of success in RNAi-based pest control, which is involved in cost and efficiency of RNAi. Therefore, we investigated the dose-sensitivity of *diap1* f-RNAi. We provided the *H. vigintioctopunctata* larvae with 0.064, 0.32, 1.6 or 8.0 ng of *diap1* dsRNA and 1000 ng of *egfp* dsRNA as the negative control (*n* = 4 in each treatment). We found that the 8.0 ng (16 ng/µl) dsRNA treatment caused significant decline of the consumed leaf area during 12–24 hours after f-RNAi (Welch’s *t* test, *P* = 0.04) (Fig. 2d, Table S5). The mean of consumed area of 8.0 ng dsRNA treatment was 9.3-fold narrower than that of the control. Notably, during both 24–36 and 38–48 hours, almost complete feeding cessation was detected in the 1.6 (3.2 ng/µl) and 8.0 ng dsRNA treated larvae (Fig. 2d, Table S5). The consumed area of 0.32 ng (0.64 ng/µl) dsRNA treated larvae was narrower to 39.2–73.5% than the control larvae during 12–48 hours after treatment, although, this treatment did not show any significant and feeding cessation effect. The 0.016 ng (0.032 ng/µl) dsRNA treated larvae showed up to only 97% less consumed area than the control larvae and no feeding cessation effect. This result shows that the feeding disorder effect of *diap1* f-RNAi has dose-sensitivity and only 1.6 ng dsRNA is sufficient to induce acute feeding cessation effect in the 3rd instar larva of *H. vigintioctopunctata* during 24–48 hours.

### Non-recoverability, lethality and growth-inhibition of *diap1* f-RNAi effect in *H. vigintioctopunctata*

To verify recoverable, lethal and/or growth-inhibitory effect of *diap1* f-RNAi in *H. vigintioctopunctata* larvae after the feed cessation effect, we observed the effects from 48 hours after the treatment with the 8 or 50 ng of *diap1* dsRNA (*N* = 7 in each target). Following our previous results, the *diap1* f-RNAi larvae exhibited feeding-cessation and non-lethality within 48 hours after the treatment with both 8 and 50 ng of dsRNA (Fig. 3a–c, Table S6). We found that the 50 and 8 ng of *diap1* dsRNA caused death of larvae from 48–72 hours (2–3 days) and 72–96 hours (3–4 days) after the treatment, respectively (Fig. 3c). In addition, all of the treated larvae exhibited lethal effects up to 4–5 and 5–6 days after the treatment with 50 and 8 ng of dsRNA, respectively (Fig. 3c). The *diap1* f-RNAi larvae did not show molting into the 4th instar (Fig. 3d). In contrast, the control larvae fed with 50 ng of *egfp* dsRNA molted into 4th instar larvae and pupae in 2–4 and 8–10 days after the treatment, respectively (Fig. 3d). We also found that almost all of the *diap1* f-RNAi larvae continued to exhibit feeding-cessation up to their death (Fig. 3a, b, Table S6). Only one larva fed with 8 ng *diap1* dsRNA showed slight feeding (0.38 mm^2^) during the day 3–4, but was dead by day 5 (Fig. 3a–c, Table S6). These results demonstrated that the f-RNAi of *diap1* causes lethality and growth-inhibition, but not conspicuous recovery of crop damage, in *H. vigintioctopunctata* larvae.

**Fig. 3.**
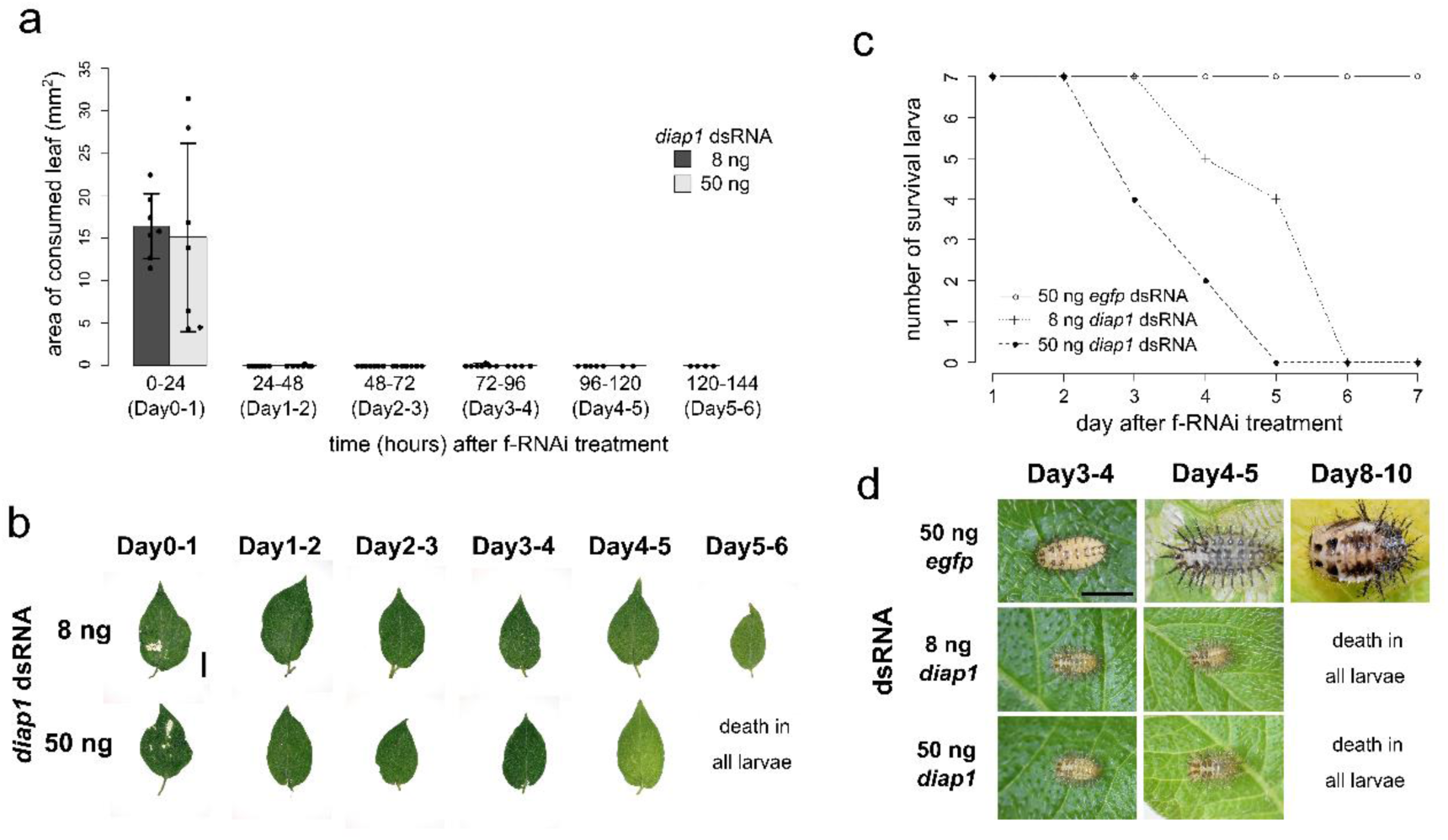
Recoverability, lethality and growth-inhibition of *diap1* f-RNAi in *H. vigintioctopunctata* larva. **a** Non-recoverability of feeding cessation of the *diap1* f-RNAi. The feeding cessation was caused within 48 hours after *diap1* f-RNAi with both the 50 and 8 ng dsRNA and persists until death of the larvae. One larva fed with the 8 ng dsRNA showed slightly feeding in the day 3, but led to death in the day 4. The data shows mean ± s.d. The starting number of larvae was 7 for each f-RNAi treatment. According to their death, the number of larvae gradually reduced during the experiment. Black circles show the number of larvae in each day. Information is provided in Table S6. **b** The leaf consumed by *diap1* f-RNAi larvae. Scale, 10 mm. **c** Lethality of the *diap1* f-RNAi larvae. All of the *diap1* f-RNAi larvae led to death up to the day 6, in contrast to surviving of all of the control larvae fed with *egfp* dsRNA during the experiment. **d** Growth-inhibitory effect of *diap1* f-RNAi. The control larvae molted into 4th instar, but not the *diap1* f-RNAi larvae. The photos show the surviving larvae at the day3-4 and 4-5 in both *diap1* and *egfp* f-RNAi. The *egfp* f-RNAi larvae further grew into pupal stage in the day 8–10, in contrast to death of all *diap1* f-RNAi larvae up to the day 6. Scale, 3 mm.

## Discussion

Searching for and selecting appropriate target gene is the keystone for effective RNAi-based pest control. In this study, we evaluated the potential of *diap1* gene as a target of RNAi-based pest control in the context of rapid reduction of crop damage by *H. viginctiopunctata*.

Our f-RNAi assay reveals that silencing of the *diap1* gene causes acute feeding disorder/cessation in *H. vigintioctopunctata* after only 24–48 hours f-RNAi (Fig. 1). In this study, the mechanism of the acute feeding disorder by *diap1* silencing is unclear. However, it is known that Diap1 promotes regeneration of gut via intestinal stem cell proliferation when intestinal tissue is injured in *Drosophila melanogaster*^26,27^. In addition, solanaceous plants produce glycoalkaloids, such as the solanine produced by potatoes, which induces cell damage and apoptosis^28^. Considering this, one possibilities is that *diap1* silencing by f-RNA suppresses intestinal stem cell proliferation via promotion of apoptosis and increases the damage caused by the defensive compounds such as solanine in *H. vigintioctopunctata* larvae. Further research is needed to elucidate the mechanism of induction of acute feeding disorders by *diap1* f-RNAi.

We clarified the species-specificity of the significant feeding disorder effect by *diap1* f-RNAi (Fig. 2). Cross-species assay using *diap1* dsRNA of other species showed a weak cross-species effect in the feeding disorder effect. Previous studies using dipteran insects have reported that *diap1* dsRNA with an identical 15 bp sequence region exhibits a cross-species effect in lethality^21^. In this study, the region of the *diap1* dsRNA had a contiguous identical 16 bp sequence (Figs. 2b, S10). Based on these facts, the cross-species effect in the feeding disorder observed in this study is considered to be due to this short identical sequence. On the other hand, the cross-species effect in this study was not significant and did not cause any feeding cessation (Fig. 2c). Therefore, we consider that *diap1* f-RNAi effect has species-specificity in the view of feeding cessation. In the practical application, the cross-species effect in feeding disorder may be completely eliminated by selecting the *diap1* dsRNA regions without >15 bp sequence identical among the target pest and the non-target insects in surrounding habitat.

Similar to the induction of the lethal effect of *diap1* f-RNAi in previous studies^21,23,24,25^, we showed that in *H. vigintioctopunctata*, the effect of *diap1* f-RNAi on feeding disorder has dose-sensitivity for dsRNA (Fig. 2d). We found that only 1.6 ng (3.2 ng/µl) of dsRNA is sufficient to induce feeding cessation in *H. vigintioctopunctata* larvae within 48 hours after the f-RNAi treatment (Fig. 2d). In our experiment, the cost of *diap1* dsRNA *in vitro* synthesis was about $0.05/µg dsRNA. Therefore, the lowest cost of feeding cessation effect of *diap1* f-RNAi is theoretically about ¢1 per a thousand *H. viginctiopunctata* larvae in our protocol. These suggest that by *diap1* silencing, induction of acute feeding disorder is achieved using very small amounts of dsRNA, and thus can be utilized as a low cost and high efficiency of RNAi-based pest control.

We showed that all of the *diap1* f-RNAi larvae lead to death during the days 2–6 days (Fig. 3). This lethal effect is rapid compared to that of the previous study in *A. planipennis* (10 days for 78% mortality)^24^. Since all of the *diap1* f-RNAi larvae were dead within 1-4 days after the feeding cessation, this rapid lethality may have occurred as a result of the acute feeding cessation, either indirectly or directly.

We compared the feeding disorder effect between *diap1* and *v-ATPase A* and *v-ATPase E*. The *v-ATPase* genes are reported as efficient targets of f-RNAi in various pests^8,14^. Significantly, we show that the acute feeding cessation can be caused by ingestion of 50 ng *diap1* dsRNA, but not that of the same dose of both *v-ATPase A* and v-*ATPase E*, in *H. vigintioctopunctata* (Fig. 1). In addition, the feeding cessation effect persists until death in the larvae fed with 50 or 8 ng *diap1* dsRNA. While we have not analyzed feeding disorder and lethality of silencing of *v-ATPase* genes from 48 hours after f-RNAi, our results suggest that the crop damage of *H. vigintioctopunctata* larvae would be more reduced by f-RNAi of *diap1* than *v-ATPase* genes and *diap1* is a more appropriate target of RNAi-based pest control in the view of crop damage reduction. We also propose that the selection of target genes for RNAi-based pest control should be evaluated not only for induction of lethality/growth-inhibition but also for induction of acute feeding disorders. In conclusion, we show that *diap1* f-RNAi in *H. vigintioctopunctata* causes the species-specific and highly efficient at inducing acute feeding disorder/cessation, which is a crucial aspect for crop damage reduction. To date, innovative application technologies for the practice of RNAi-based pest control have been developed^29^. The main application technologies are as follows: 1) Utilization of *in vitro* synthesized dsRNA: for example, a method of drying industrially produced dsRNA and spraying it^5^, 2) Utilization of *in planta* synthesized dsRNA: for example, a method for producing transgenic plants that synthesizes dsRNA targeting the pest genes in cell^8,11,12^ or chloroplast^4^, 3) Use of *in vivo* synthesized dsRNA: for example, spraying a large amount of dsRNA synthesized efficiently by bacteria^30,31^. In this study, we provided the larvae with a droplet containing dsRNA as an experimental model, but it is our methods are applicable for any of the technologies described above. Recently, it has been clarified that silencing of a dsRNA binding protein-coding gene, *Staufen* homolog, improves RNAi efficiency via reducing resistance for RNAi in the Colorado potato beetle^32^. Combining *Staufen* homolog and *diap1* silencing may cause the feeding disorder effect more efficiently in some pests.

It is expected that searching and selecting target genes inducing acute feeding disorder/cessation by f-RNAi in various herbivorous pests will realize efficient RNAi-based pest control. This will be able to solve problems such as the dsRNA synthesis cost, or persistence of crop damage until lethal or growth-inhibitory effect appears.

## Material and Methods

### Insects

*Henosepilachna vigintioctopunctata* were collected from potato leaves at either Nagoya University or National Institute for Basic Biology, in Japan and used these animals to establish a laboratory line. Laboratory stocks of *Harmonia axyridis* were derived from field collections in Aichi, Japan. They were reared as described by Niimi *et al*.^34^ *Helicoverpa armigera* was kindly provided by Dr. Chie Goto (NARO Agricultural Research Center). *Blattela lateralis* were purchased from Remix. The larvae of *Oxya yezoensis* were collected from rice field at Nagoya University.

### Cloning of partial *diap1* sequences from insects

Total RNA was extracted from the gonad of *H. vigintioctopunctata*, the embryos of *H. axyridis*, the anterior wing primordia of *H. armigera*, the posterior wing primordia of *B. lateralis* or the larval posterior legs of *O. yezoensis* using TRIzol (Invitrogen, California, USA) according to the manufactural protocol. The first-strand cDNA was synthesized using the SuperScript II Reverse Transcriptase (Life Technologies Japan Ltd., Tokyo, Japan) with SMART RACE cDNA Amplification kit (Clontech, Mountain View, California, USA) from 1 µg total RNA. *diap1* cDNA fragment of each species was amplified using degenerated primer sets designed from conserved amino acid sequences among insects (diap1_F1 or diap1_F2/diap1_R in Table S8). The RT-PCR were performed using AmpliTaq Gold DNA polymerase (Perkin Elmer, Boston, USA). PCR fragments were separated with electrophoresis using 1% agarose gel in tris-borate-EDTA (TBE) buffer and extracted using MagExtractor (TOYOBO Co. Ltd., Osaka, Japan). Each PCR fragment was subcloned into the EcoRV recognition site of pBluescript KS (+) vector (Stratagene, La Jolla, CA, USA) using DNA Ligation kit Ver. 2 (TaKaRa bio. Inc., Shiga, Japan). The vector was transformed into XL1-Blue *Escherichia coli* competent cells (GMbiolab Co., Ltd, Taichung, Taiwan). The plasmid was extracted from the transformed colony using FlexiPrep Kit (Amersham Pharmacia Biotech Inc., New Jersey, USA). The nucleotide sequences of the PCR fragment inserted into vectors were confirmed using the dideoxy chain-termination method using an automatic DNA sequencer (CEQ 2000XL; Beckman Coulter, California, USA). Sequence analysis was carried out using a DNASIS system (Hitachi Software Engineering, Tokyo, Japan). The partial length *diap1* nucleotide sequences of the insects were deposited in DNA Data Bank of Japan (DDBJ) (accession numbers: LC473085 in *H. axyridis*, LC473088 in *H. armigera*, LC473087 in *B. lateralis* and LC473086 in *O. yezoensis*).

### Rapid amplification of cDNA end (RACE) of *diap1* in *H. vigintioctopunctata*

To obtain a full-length *diap1* cDNA in *H. vigintioctopunctata*, the 5’- and 3’- rapid amplification of cDNA end (RACE) were performed according to the manufacturer’s protocol of the SMART RACE cDNA Amplification kit (Clontech, Mountain View, California, USA). The gene-specific primers are described in Table S8 (1st PCR: Hvig_diap1_1 for 5’-RACE and Hvig_diap1_3 for 3’-RACE; nested PCR: Hvig_diap1_2 for 5’ RACE and Hvig_diap1_4 for 3’RACE). These primers were designed from the *diap1* cDNA fragment cloned above. The cloning and confirmation of nucleotide sequences of RACE fragment were performed according to the same way as the above method. The full length *diap1* nucleotide sequences of *H. vigintioctopunctata* was deposited in DDBJ (accession numbers: LC473084). The *diap1* full-length nucleotide sequence is shown in Supplementary Sequences.

### Transcriptomic analysis

In order to obtain *v-ATPase A* and *v-ATPase E* sequences of *H. vigintioctopunctata*, transcriptomic analysis was performed using total RNA from the whole body of *H. vigintioctopunctata* 3rd instar larvae. Total RNAs were extracted using the RNeasy Mini kit (QIAGEN, Tokyo, Japan). The integrity of extracted RNAs were confirmed using a Bioanalyzer 2000 (Agilent Technologies) and these RNAs were used to construct cDNA libraries using the Truseq™ RNA Sample Prep Kit (Illumina) according to the manufacture’s instruction. Pair-end sequences with 200 bp were obtained from these cDNA libraries through Hiseq2000 (Illumina). The raw reads were deposited in DDBJ Sequence Read Archive (accession number: DRA008720). We rearranged pair-end reads in order and excluded unpaired reads using cmpfastq_pe (http://compbio.brc.iop.kcl.ac.uk/software/cmpfastq_pe.php). Low quality reads and adapter sequences were trimmed from obtained short-reads using Cutadapt v1.15^35^. Then, *de novo* assembly of trimmed short-read sequence was performed using Trinity-v4.2.0 (https://github.com/trinityrnaseq/trinityrnaseq/)^36^. We also analyzed the RNA-seq data of *H. axyridis* obtained from National Center for Biotechnology Information (NCBI) Sequence Read Archive database (run number.: ERR1309559) in the same way as *H. vigintioctopunctata*.

### BLAST search and phylogenetic analysis of target genes

BLAST databases of the assembled transcriptomic sequences in *H. vigintioctopunctata* and *H. axyridis* were established by makeblastdb program in BLAST+ ver. 2.7.1^37^. We searched homologous genes of *v-ATPase A, v-ATPase E* and *diap1* in the *H. vigintioctopunctata* BLAST database with the orthologues in *Drosophila melanogaster* (accession number: Diap1, NP_001261916.1; V-ATPase A, NP_652004.2; V-ATPase E, NP_524237.1) obtained from NCBI database as query sequences using the tblastn program in BLAST+ ver. 2.7.1. Then, we confirmed the orthology of these obtained sequences by molecular phylogenetic analysis (Fig. S4). The homologous sequences of V-ATPase A, V-ATPase E and Diap1 in other animals were obtained from NCBI database (Table S7). All sequences including *H. vigintioctopunctata* homologues were aligned using MAFFT ver. 7^38^ with the L-INS-i program. The using sites for phylogenetic construction were selected by trimAl v1.2^39^ with a gap threshold value of 0.7. The multiple alignments were shown in Figs. S7-S9. The selection of best-hit substitution model and construction of maximal-likelihood phylogeny were performed using MEGA X^40^. Bootstrap values were calculated after 1000 replications. BLAST search and multiple alignment were performed under Super Computer Facilities of National Institute of Genetics. The *diap1, v-ATPase A* and *v-ATPase E* nucleotide sequences of *H. vigintioctopunctata* and *H. axyridis* are shown in Supplementary Sequences.

### Expression profile analysis of target genes in larval tissues

Total RNA was extracted from nerve cord, legs, wing discs fat body, Malpighian tubules, gut and carcass of 4th instar larvae of *H. vigintioctopunctata* which is the dissectable stage for each tissue using the RNeasy Mini kit (QIAGEN, Tokyo, Japan). The first-strand cDNA was synthesized from the 1 µg total RNA using SuperScript III Reverse Transcriptase (Life Technologies Japan Ltd., Tokyo, Japan). RT-PCR for the target genes and the internal control (*ribosomal protein 49* (*rp49*), accession number: AB480201) was performed using the first-strand cDNA and Q5 High-Fidelity DNA Polymerase (New England Biolabs Japan Inc., Tokyo, Japan) with 30 cycles. Primers for RT-PCR were designed by Primer3web version 4.1.0^41^ and are shown in Table S8 (Hvig_diap1_RT-PCR_F/Hvig_diap1_RT-PCR_R, Hvig_v-ATPase_A_RT-PCR_F/Hvig_v-ATPase_A_RT-PCR_R, Hvig_v-ATPase_E_RT-PCR_F/Hvig_v-ATPase_E_RT-PCR_R and Hvig_rp49_RT-PCR_F/Hvig_rp49_RT-PCR_R). The annealing temperatures for RT-PCR were calculated by NEB Tm Calculator (https://tmcalculator.neb.com/). The expression pattern was confirmed by agarose gel electrophoresis of the PCR products with 2% Agarose S (NIPPON GENE Co. Ltd., Toyama, Japan).

### Double strand RNA synthesis

For comparison analysis of RNAi efficiency of target genes, the total RNA was extracted from 3rd instar larvae of *H. vigintioctopunctata* using TRI Reagent (Molecular Research Center Inc., Ohio, USA) according to the manufactural protocol. The first-strand cDNA was synthesized from 1 µg total RNA using SuperScript III Reverse Transcriptase (Life Technologies Japan Ltd., Tokyo, Japan). The templates of dsRNA were initially amplified from the first-strand cDNA and secondary amplified from the initial PCR products by RT-PCR using Q5 High-Fidelity DNA Polymerase (New England Biolabs Japan Inc., Tokyo, Japan). The initial PCR primers are described as Hvig_diap1_F/Hvig_diap1_R, Hvig_v-ATPase_A_F/Hvig_v-ATPase_A_R and Hvig_v-ATPase_E_F/Hvig_v-ATPase_E_R in Table S8. The nested primers were flanked with T7 promoter sequences on the 5’ ends (Hvig_T7-diap1_F/Hvig_T7-diap1_R, Hvig_T7-v-ATPase_A_F/Hvig_T7-v-ATPase_A_R and Hvig_T7-v-ATPase_E_F/Hvig_T7-v-ATPase_E_R in Table S8). The nested PCR products were purified using MagExtractor (TOYOBO Co. Ltd., Osaka, Japan). The sequences of interest were confirmed by sanger DNA sequencing service at FASMAC Co. Ltd. (Kanagawa, Japan). The dsRNAs of target genes were synthesized from the purified PCR products using AmpliScribe T7-Flash Transcription Kit (Epicentre Technologies, Co., Wisconsin, USA).

For Species-specificity and dose analysis of *diap1*, the templates of dsRNA were amplified from the above pBluescript KS (+) vectors inserted *diap1* sequences in each insect by PCR using AmpliTaq Gold (Perkin Elmer, Boston, USA). The PCR primers are described in Table S8 (T7-KS/T7-SK). The PCR products were purified by the same ways to the above method. The dsRNAs were synthesized using MEGAscript T7 Transcription Kit (Ambion, Texas, USA).

### Feeding RNAi assay

Synthesized dsRNA was artificially fed to 3rd instar larvae of *H. vigintioctopunctata* within 24 hours after molting (Fig. S5). The 0.5 µl solution including dsRNA of each gene was dropped on slide glasses in front of the larvae which had been starved after molting to the 3rd instar. The larvae completely consumed the dsRNA solution. The elapsed time in consumed the solution was 114–542 seconds (median time = 206.5 seconds). After the larvae had consumed the entire droplet, each larva was put on a fresh potato leaf and kept in a plastic container at room temperature. Body weight of the larvae after 48 hours were measured using a precision electronic balance (Table S1). The potato leaves were collected and exchanged for new potato leaves up to 48 hours every 12 or 24 hours (Fig. 1e). dsRNA of *enhanced green fluorescent protein* (*egfp*) were fed to 3rd instar larvae as the negative control.

### Evaluation of knockdown of mRNAs expression by qRT-PCR

The total RNA was extracted from the whole-body of *H. vigintioctopunctata* 3rd instar larvae 48-hours after f-RNAi treatment using TRI Reagent (Molecular Research Center Inc., Ohio, USA). The first-strand cDNA was synthesized from 1 µg total RNA using SuperScript III Reverse Transcriptase (Life Technologies Japan Ltd., Tokyo, Japan). qRT-PCR was performed on LightCycler 96 instrument (Roche, Basel, Swizerland) using THUNDERBIRD SYBR qPCR Mix (TOYOBO Co. Ltd., Osaka, Japan) according to the manufactural protocol with the first-strand cDNA as the template. The same primer sets as the above expression profile analysis of each gene were used. The expression level of target genes relative to *rp49* was calculated by the 2^-ΔΔCt^ method^42^.

### Measurements of leaf area eaten by feeding RNAi larvae

We measured the potato leaf area eaten by f-RNAi larvae using the following method with the digital microscope system (VHX-5000, KEYENCE, Osaka, Japan). A leaf was put on the glass plate on the stage and was covered with a plastic wrap. Then, the region of leaf eaten was manually bordered with a black pen. After focusing on the leaf using 10x magnification, we automatically binarized the image and measured the area of the region eaten by larvae with the digital microscope system.

### Statistical analysis

We performed the Welch’s *t*-test to evaluate the differences of effects of f-RNAi assay. We adjusted *P*-value with the Holm’s methods for the multiple-comparisons. The significance level used in our analysis was *P*-value < 0.05. All statistical analyses were performed by R-3.6.0 (https://cran.r-project.org/).

### Expression level of *diap1* in tissues of *Drosophila melanogaster*

We obtained the expression dataset of *diap1* (FBgn0260635) in tissue of *Drosophila melanogaster* from FlyAtlas Anatomical Expression Data^43^. Then, we created the histogram based on the expression data (Supplementary Fig. 1).

### Sequence analysis of target genes among insects

The nucleotide and amino acid sequences of target genes in 9 insect species were obtained from NCBI’s database (Table S7). Then, multiple alignments were performed for each pair of species using MAFFT ver.7 with L-INS-i program. The proportions of identical sites for length of coding sequence region of each gene were manually calculated. The identical sites of partial sequences of *diap1* between *H. vigintioctopunctata* and other insects using for f-RNAi assay were identified in the same way (alignment: Fig. S10).

## Acknowledgements

We would like to thank to R. A. Zinna (Mars Hill University) for his critical read of this manuscript. We also thank J. Yatomi for helping in a part of this work and T. Ando, T. Nakamura and H. Sakai (National Institute for Basic Biology, Japan) for helpful discussions. We express our gratitude to C. Goto (NARO Agricultural Research Center, Japan) for providing *Helicoverpa armigera* and T. Konagaya (National Institute for Basic Biology, Japan) for the critical advice to statistical analysis. We thank the Model Plant Research Facility, NIBB BioResource Center for their technical support. This work was supported in part by the Center for the Promotion of Integrated Sciences (CPIS) of SOKENDAI (to T.N. and Y.S.).

## Contributions

Y.C., H.Y., Y.S., T.Y. and T.N. conceived and designed the study. Y.C., H.K. and T.N. performed the majority of experiments. Y.C., T.S. and Y.S. analyzed RNA-seq data. H.K. maintained *H. vigintioctopunctata* and potato in laboratory. T.N. supervised the study. Y.C. and T.N. wrote the manuscript. All authors discussed the manuscripts.

