## supplementary file for "Oral RNAi of *diap1* in a pest results in rapid reduction of crop damage"

### Supplementary Figures

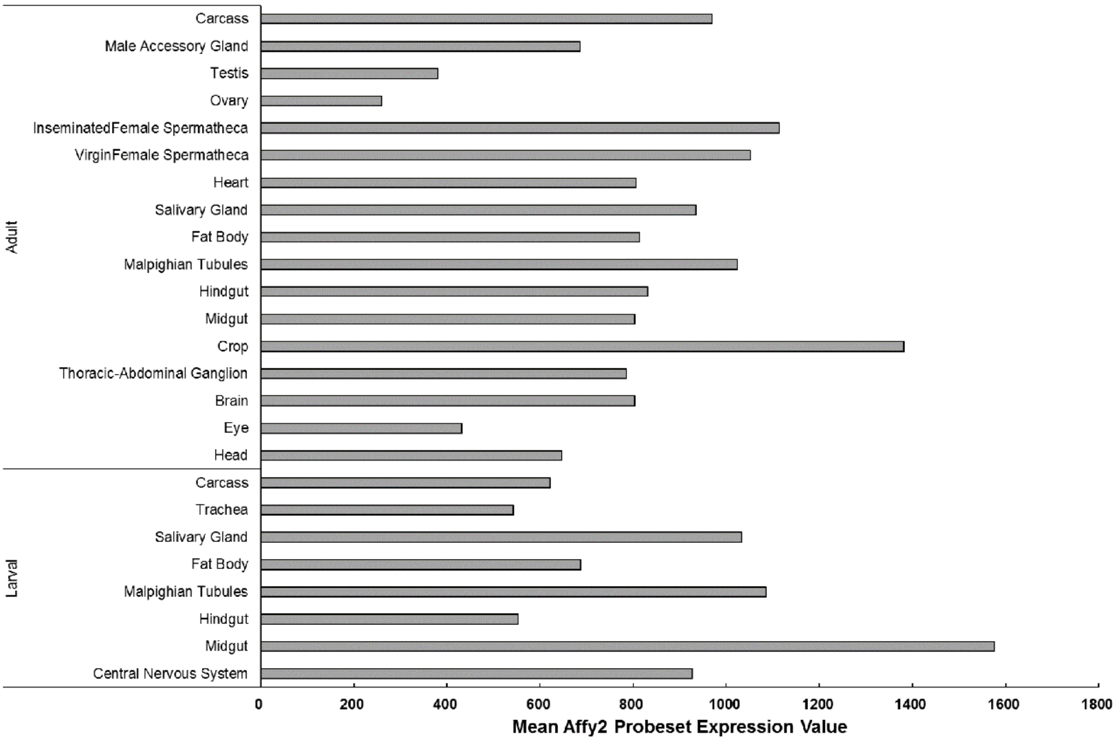


**Fig. S1. Expression of *diap1* in various tissues in *Drosophila melanogaster*.**

Expression dataset obtained from flybase (http://flybase.org/) shows that *diap1* ubiquitously expresses in *Drosophila* tissues, and is detected the highest expression in larval gut.

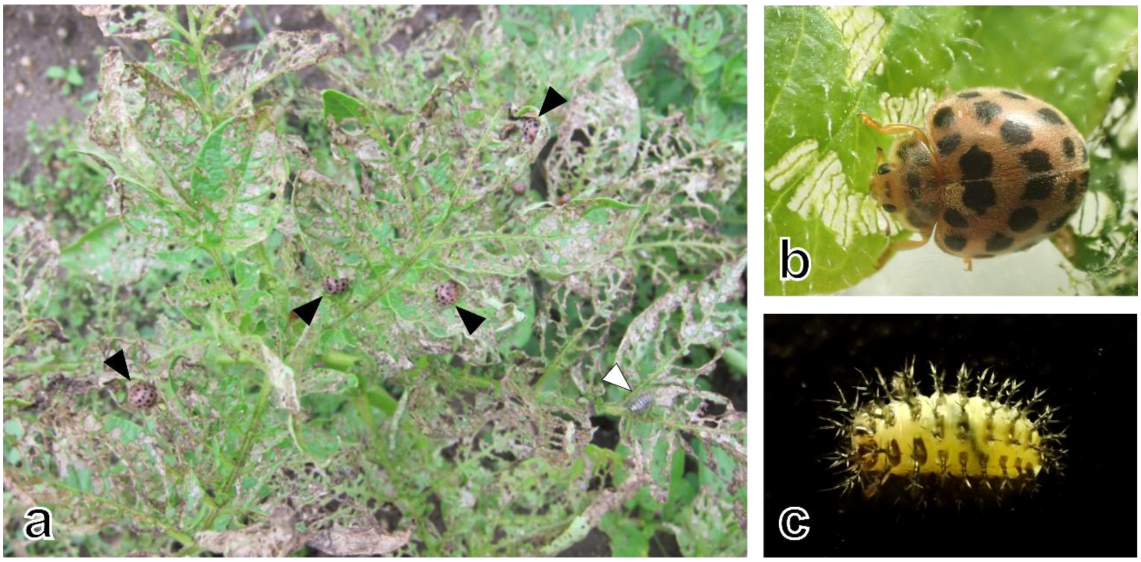


**Fig. S2.** ***Henosepilachna vigintioctopunctata*****, a global pest for Solanaceae plants.** *H. vigintioctopunctata* eats leaves of Solanaceae plants during their all life stage other than the embryo. (a) *H. vigintioctopunctata* on the potato leaves. Black arrowhead shows adults. White arrowhead shows larvae. (b) adult of *H. vigintioctopunctata.* (c) larvae of *H. vigintioctopunctata*.

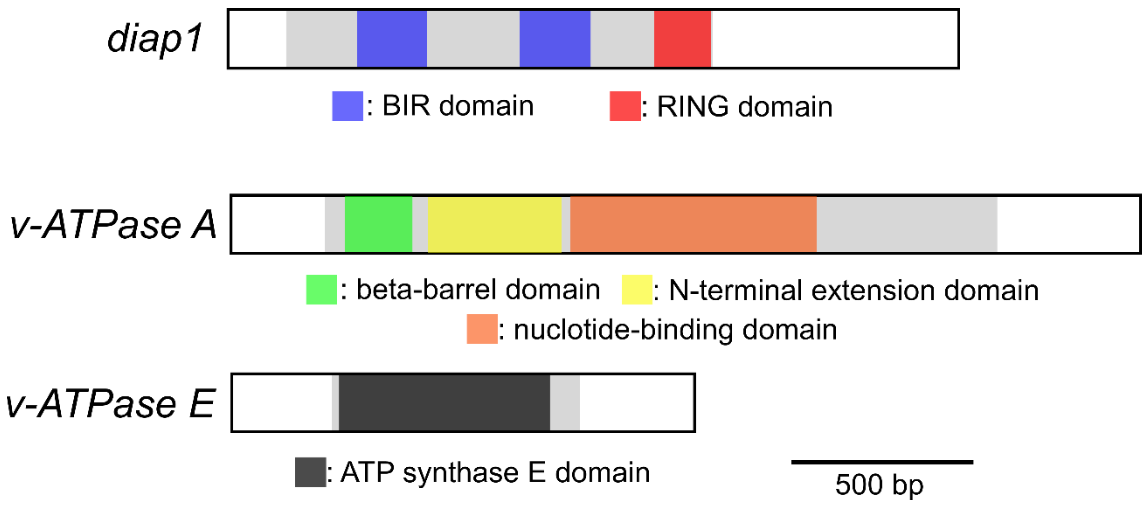


**Fig. S3. Gene structure of target genes in *H. vigintioctopunctata*.**

*diap1* has three conserved domains, two BIR and one RING domain. *v-ATPase A* has three ATP synthase alpha/beta family domain, one beta-barrel, one N-terminal extension and one nucleotide-binding domain. Overall *v-ATPase E* sequence is strongly conserved. White and gray boxes show untranslated and coding region, respectively.

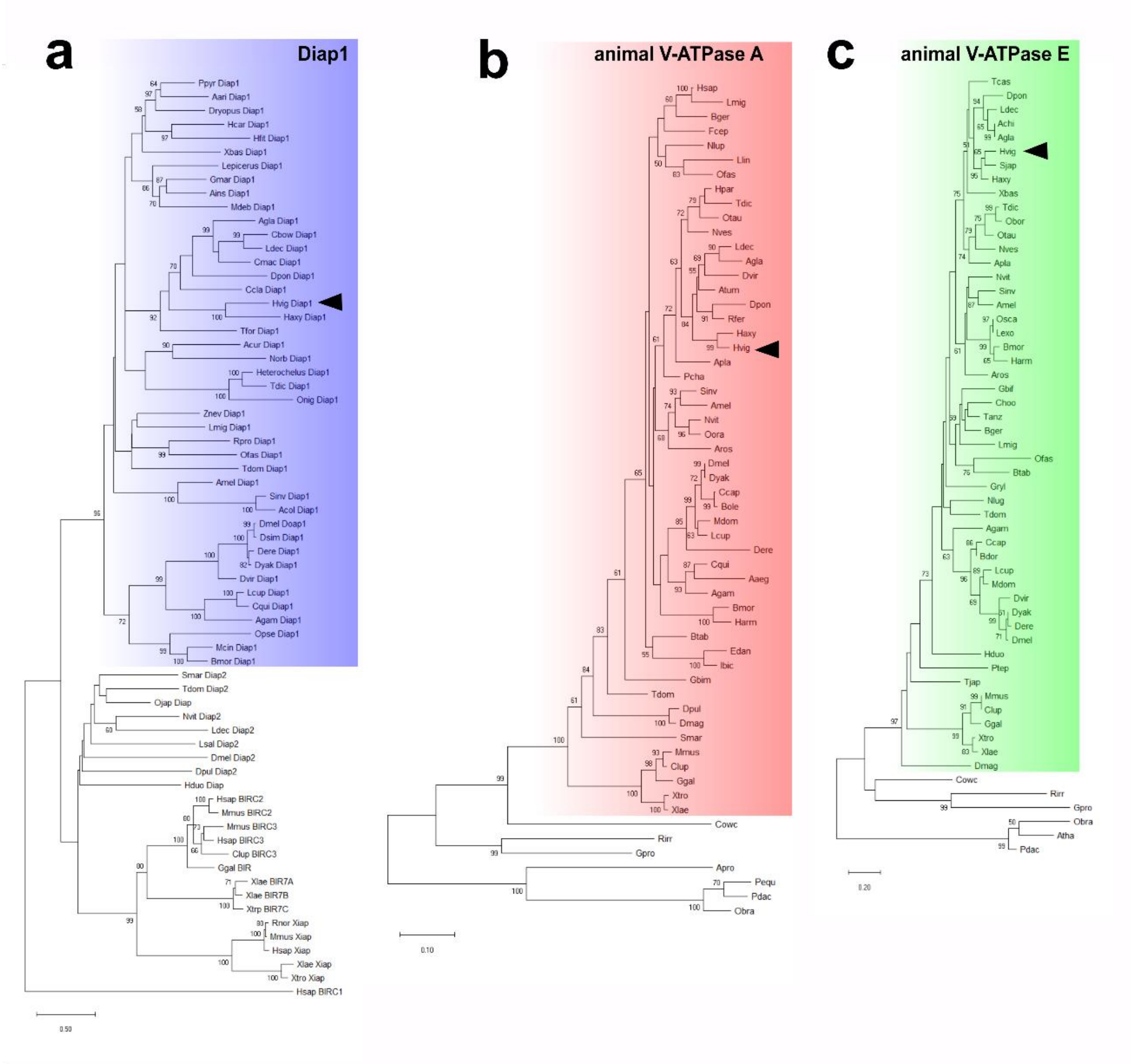


**Fig. S4. Maximum-likelihood phylogenetic analysis of target genes.**

(a) Iap phylogeny. The amino acid sequences of Iap family among Arthropoda and Vertebrata were obtained from NCBI and Swissprot database. Blue area indicates insect Diap1 clade. This clade is robustly supported by bootstrap value (96%). Our sequence from *H. vigintioctopunctata* (black arrowhead) locates in the Diap1 clade. LG+G+I model. (b) V-ATPase A phylogeny. The amino acid sequences of V-ATPase A among Eukaryote were obtained from NCBI, Swissprot and i5k database. Red area shows animal V-ATPase A clade, which is strongly supported by bootstrap value (100%). Our sequence of *H. vigintioctopunctata* (black arrowhead) puts in the clade. LG+G+I model. (c) V-ATPase E phylogeny. The amino acid sequences of V-ATPase E among Eukaryote were obtained from NCBI, Swissprot and i5k database. Green area shows animal V-ATPase E clade, which is strongly supported by bootstrap value (97%). Our sequence of *H. vigintioctopunctata* (black arrowhead) puts in the clade. LG+G+I model. The bootstrap value <50% is not shown. Datasets used in the phylogenetic analysis show in Table S7.

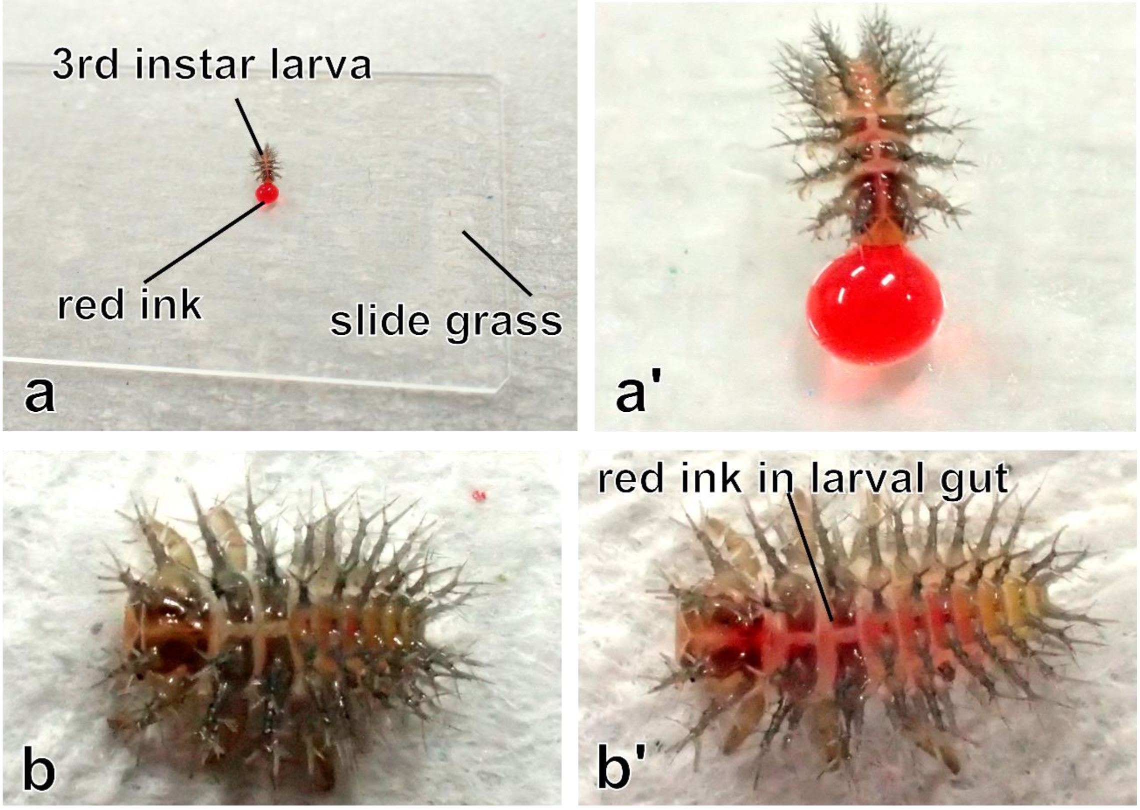


**Fig. S5. Feeding RNAi assay of *H. vigintioctopunctata*.**

These figures show the demonstration of feeding assay using food color. (a) the 3rd instar larvae were put in front of droplet of the food color on a slide grass, and immediately ate or drank the food color. (a') high-magnified image of a. (b) 3rd instar larva before feeding the food color. (b') 3rd instar larva after feeding the food color. The food color was taken up into the midgut of larva. F-RNAi assays were performed as the same way using double-stranded (ds) RNA of each gene instead of a food color.

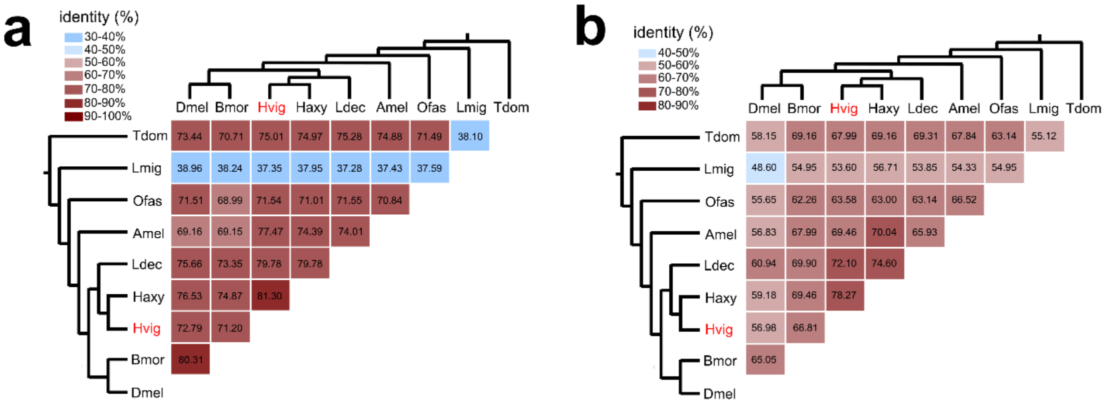


**Fig. S6. The nucleotide identities of *v-ATPase A* and *v-ATPase E* among 9 insect species.**

(a) *v-ATPase A*. *v-ATPase A* shows 37.28-81.3% identity. (b) *v-ATPase E*. *v-ATPase E* shows 48.6-78.27% identity. *v-ATPase A and v-ATPase E* sequences was obtained from RNA-seq data of NCBI database. Numbers in each column indicates identity between species. Branches show the phylogenetic relationship[33] among species. Amel, *Apis melifera*; Bmor, *Bombyx mori*; Caqu, *Catajapyx aquilonaris*; Dmel, *Drosophila melanogaster*; Hvig, *Henosepilachna vigintioctopunctata*; Haxy, *Harmonia axyridis*; Ldec, *Leptinotarsa decemlineata*; Lmig, *Locusta migratoria*; Ofas, *Oncopeltus fasciatus*.

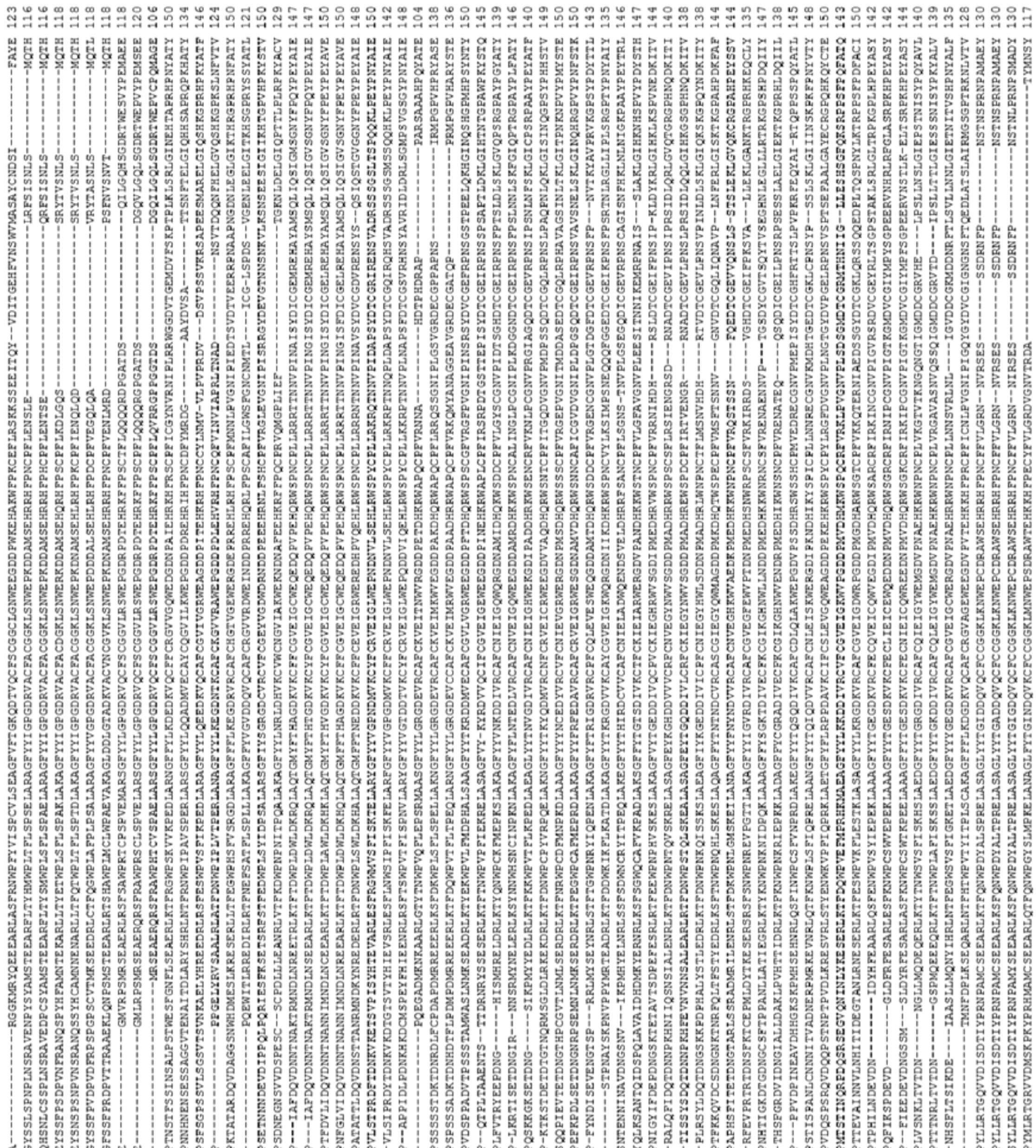

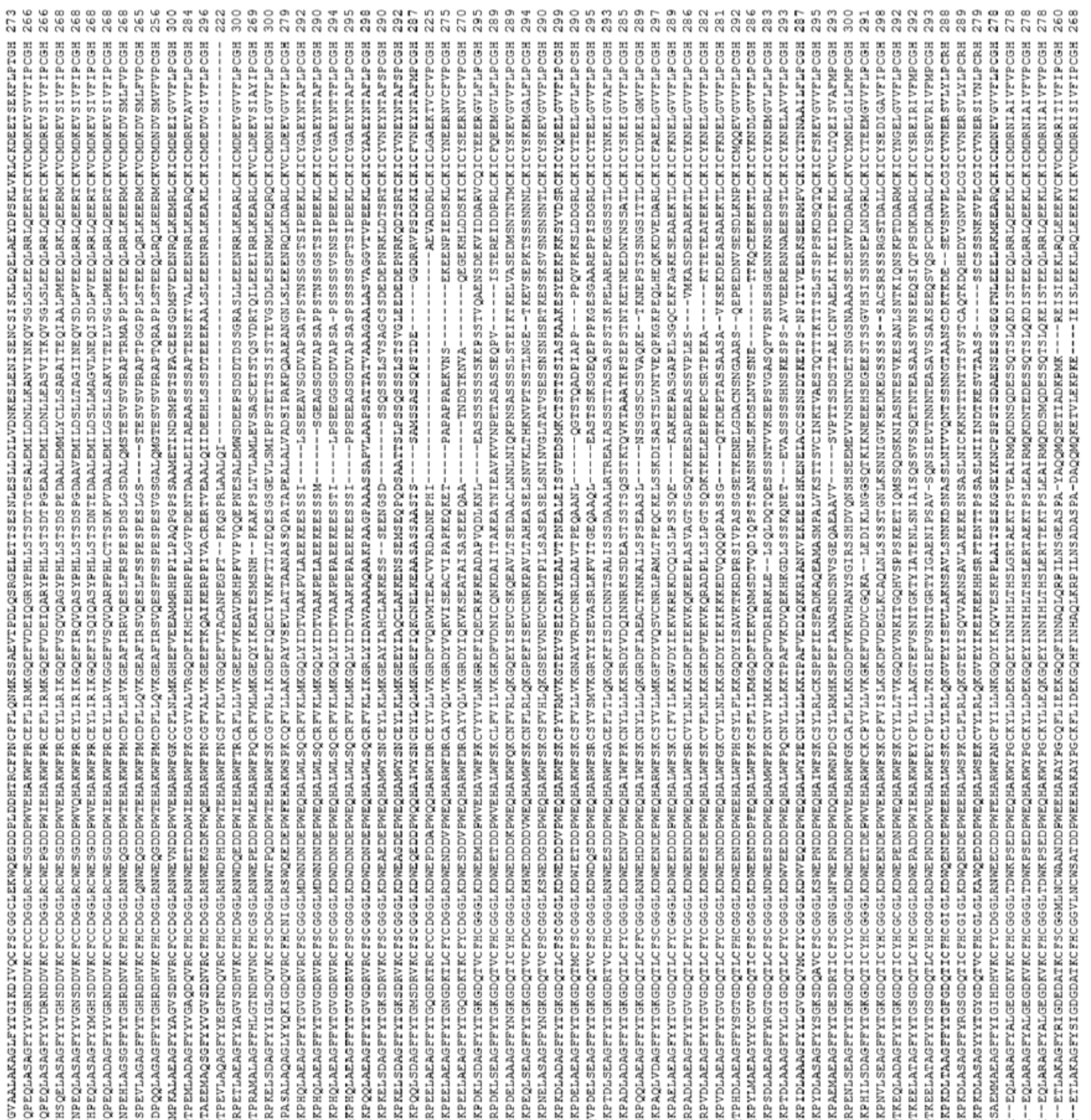

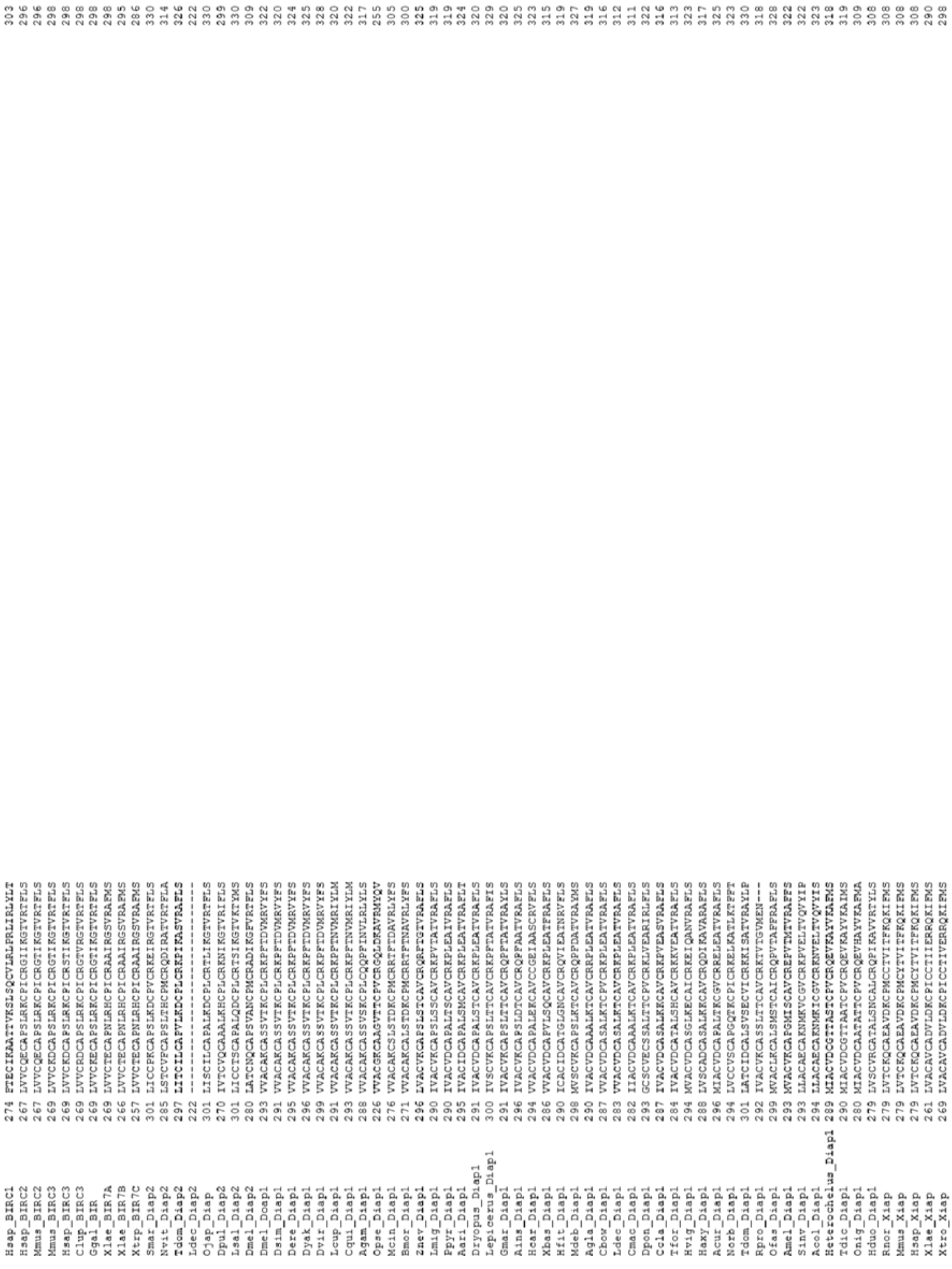

**Fig. S7. Multiple alignments of amino acid of Iap used to phylogenetic analysis.**

Amino acid sequences were obtained from 67 animal species. 330 residues were used to phylogenetic reconstruction. Reconstructed phylogenetic trees show in Fig. S4a. Datasets used in the phylogenetic analysis show in Table S7.

64

```
Hsap  1 TSTLIKTSDEDRESKFGFVFAVSGPVVTAERMAGSAMYELVRVGYYELVGEIIRLEGDMATIQVYEDTSGVTVGDPVLRTGKPLSVELGPGIMGSIFDGIQRPLKDINELSNSIYIPKGVNVPALSRTAQWDFSPVSVKVGSHITGGDLY 150
Bger  1 SPGLPKMNDEDRESKYGYVFAVSGPVVTAERMSGSAMYELVRVGYFELVGEIIRLEGDMATIQVYEETSGVTVGDPVLRTGKPLSVELGPGILGSIFDGIQRPLKDINELTQSIYIPKGVNVPALSRTQQWDFAPAVVKVGSHITGGDMY 150
Hpar  1 MSRLPKIGDEERESNYGYVFAVSGPVVTAEKMSGSAMYELVRVGYYELVGEIIRLEGDMATIQVYEETSGVTVGDPVLRTGKPLSVELGPGILGSIFDGIQRPLKDINDLTQSIYIPKGVNVPALSKTTAWEFNPLNIKAGAHLTGGDLY 150
Otau  1 MSKLAKLSDEDKESTYGYVYAVSGPVVTAEKMSGSAMYELVRVGYHELVGEIIRLEGDMATIQVYEETSGVTVGDPVLRTGKPLSVELGPGILGSIFDGIQRPLKDINDLTQSIYIPKGINVPALSRTINWEFNPLNIKVGAHLTGGDLY 150
Tdic  1 MSRLAKMTDEERESYYGYVFAVSGPVVTAQKMSGSAMYELVRVGYFELVGEIIRLEGDMATIQVYEETSGVTVGDPVLRTGKPLSVELGPGILGSIFDGIQRPLKDISDLMQSIYIPKGINIPALSRTQTWDFNPLNIKAGAHLTGGDLY 150
Nves  1 -MSLPKIRDEEKESNYGYVYAVSGPVVTAEKMSGSAMYELVRVGYYELVGEIIRLEGDMATIQVYEETSGVTVGDPVLRTGKPLSVELGPGILGSIFDGIQRPLKDINDLTQSIYIPKGVNVPALSKTTQWEFMPHNIKMGSHLTGGDLY 149
Pcha  1 MSRLPKIADEEREGKYGYVFAVSGPVVTAEKMSGSAMYELVRVGYYELVGEIIRLEGDMATIQVYEETSGVTVGDPVLRTGKPLSVELGPGILGSIFDGIQRPLKDINELTQSIYIPKGVNVPALSRTQKWEFNPLNIKNGAHITGGDLY 150
Apla  1 MSKLIKVGDVEKETTYGYVHAVSGPVVTAEKMSGSAMYELVRVGYYELVGEIIRLEGDMATIQVYEETSGVTVGDPVLRTGKPLSVELGPGIMGSIFDGIQRPLKDINELTQSIYIPKGVNIPALSRTASWEFNPLNIKLGSHMTGGDLY 150
Sinv  1 NHSLAKIAHEEREGKFGYVYAVSGPVVTAEKMSGSAMYELVRVGYYELVGEIIRLEGDMATIQVYEETSGVTVGDPVLRTGKPLSVELGPGILGSIFDGIQRPLKDINELTSSIYIPKGINVPALSRTASWEFNPHNVKNGSHITGGDLY 150
Nvit  1 -MTLPKIANVEKEAKFGYVYAVSGPVVTAEKMSGSAMYELVRVGYDELVGEIIRLEGDMATIQVYEETSGVTVGDPVLRTGKPLSVELGPGILGSIFDGIQRPLKDINELTGSIYIPKGVNTPALSRTIAWEFNPLYIKNGSHITGGDMY 149
Oora  1 -------------------------------MSGSAMYELVRVGYHELVGEIIRLEGDMATIQVYEETSGVTVGDPVLRTGKPLSVELGPGILGSIFDGIQRPLKDINELTSSIYIPKGVNVPCLSRTTSWEFNPLYIKTGSHITGGDVY 119
Amel  1 SQGLLKISNEERETKFGYVYAVSGPVVTAEQMSGSAMYELVRVGYYELVGEIIRLEGDMATIQVYEETSGVTVGDPVLRTGKPLSVELGPGILGSIFDGIQRPLKDINELTNSIYIPKGINVPALSRTAAWEFNPSNIKNGSHITGGDLF 150
Aros  1 SKGLSKGRDGEKESEYGYVYAVSGPVVTAEKMAGSAMYELVRVGYYKLVGEIIRLEGDMATIQVYEETSGVTVGDPIVRTGKPLSVELGPGILGSIFDGIQRPLKDINELTSSIYIPKGVNMPALSRTAQWEFNPLNIKKGSHITGGDLY 150
Dmel  1 MSNLKRFDDEERESKYGRVFAVSGPVVTAEAMSGSAMYELVRVGYYELVGEIIRLEGDMATIQVYEETSGVTVGDPVLRTGKPLSVELGPGIMGSIFDGIQRPLKDINELTESIYIPKGVNVPSLSRVASWEFNPLNVKVGSHITGGDLY 150
Dyak  1 MSNLKRFDDEERESKYGRVFAVSGPVVTAEAMSGSAMYELVRVGYYELVGEIIRLEGDMATIQVYEETSGVTVGDPVLRTGKPLSVELGPGIMGSIFDGIQRPLKDINELTESIYIPKGVNVPSLSRVASWEFNPLNVKVGSHITGGDLY 150
Ccap  1 MSNLKRFDDEDNEGKYGRVFAVSGPVVTAERMSGSAMYELVRVGYYELVGEIIRLEGDMATIQVYEETSGVTVGDPVLRTGKPLSVELGPGIMGSIFDGIQRPLKDINELTESIYIPKGVNVPSLSRTQAWEFNPLNVKVGSHITGGDLY 150
Bole  1 MSNLKRFDDEDNESKYGRVFAVSGPVVTAERMAGSAMYELVRVGYYELVGEIIRLEGDMATIQVYEETSGVTVGDPVLRTGKPLSVELGPGIMGSIFDGIQRPLKDINELTESIYIPKGVNVPSLSRTQAWEFNPLNVKVGSHITGGDLY 150
Mdom  1 MSNLKRFDDEERESRYGRVYAVSGPVVTAECMSGSAMYELVRVGYYELVGEIIRLEGDMATIQVYEETSGVTVGDPVLRTGKPLSVELGPGIMGSIFDGIQRPLKDINELTSSIYIPKGVNVPCLSRTATWEFNPLNVKVGSHITGGDLY 150
Lcup  1 MSNLKRFDDVEEESKYGRVFAVSGPVVTAECMAGSAMYELVRVGYYELVGEIIRLEGDMATIQVYEETSGVTVGDPVLRTGKPLSVELGPGIMGSIFDGIQRPLKDINELTNSIYIPKGVNVPCLSRTASWEFNPLNVKVGSHITGGDLY 150
Cqui  1 MSNLKKIADEDRESKFGYVFAVSGPVVTAEKMAGSAMYELVRVGYYELVGEIIRLEGDMATIQVYEETSGVTVGDPVLRTGKPLSVELGPGIMGSIFDGIQRPLKDINELTQSIYIPKGINTPSLSRTQSWGFNPMNVKVGSHITGGDLY 150
Agam  1 MSNLKKISDEDRESKFGYVFAVSGPVVTAERMSGSAMYELVRVGYYELVGEIIRLEGDMATIQVYEETSGVTVGDPVLRTGKPLSVELGPGIMGSIFDGIQRPLKDINEMTSSIYIPKGINVPCLSRTQSWSFNPLNVKAGSHITGGDLY 150
Fcep  1 --MLPKIRDEDKESRFGYVYAVSGPVVTAERMSGSAMYELVRVGYHELVGEIIRLEGDMATIQVYEETSGVTVGDPVLRTGKPLSVELGPGILGSIFDGIQRPLKDISELAGSIYIPKGVNVPALSRASSWEFNPLSIKIGSHMTGGDLY 148
Ldec  1 MSRL--IADEEQESQYGYVHAVSGPVVTAEKMSGSAMYELVRVGYYELVGEIIRLEGDMATIQVYEETSGVTVGDPVLRTGKPLSVELGPGIMGSIFDGIQRPLKDICDMTESIYIPKGVNVPSLSRTIKWEFNPINIKLGSHLTGGDIY 148
Agla  1 MSKLPKIGDEEREQQYGYVHAVSGPVVTANEMSGSAMYELVRVGYSELVGEIIRLEGDMATIQVYEETSGVTVGDPVLRTGKPLSVELGPGIMGSIFDGIQRPLKDICDMTESIYIPKGVNVPALSRTQKWEFSPLNIKLGSHLTGGDIY 150
Dvir  1 MSKV-RIGDEEKEGQYGYVHAVSGPVVTAEKMSGSAMYELVRVGYYELVGEIIRLEGDMATIQVYEETSGVTVGDPVLRTGKPLSVELGPGIMGSIFDGIQRPLKDICDATDSIYIPKGINVPSLSRTAKWDFNPINIKLGSHLTGGDIY 149
Haxy  1 MSNLRLIRDEEKESEYGYVHAVSGPVVTAEKMCGSAMYELVRVGYFELVGEIIRLEGDMATIQVYEETSGVTVGDPVLRTGKPLSVELGPGIMGSIFDGIQRPLKDINVLTESIYIPKGINVPCLSRTAKWDFNPINIKLGSHLTGGDIY 150
Hvig  1 MSNLPKISDEEREANYGYVHAVSGPVVTAEKMCGSAMYELVRVGYFELVGEIIRLEGDMATIQVYEETSGVTVGDPVLRTGKPLSVELGPGIMGSIFDGIQRPLKDINDLTQSIYIPKGVNVPALSRTSKWDFNPYNIKLGSHLTGGDIY 150
Dpon  1 MSKLIKLGDEESDGRFGYVHMVSGPVVTAEKMSGSAMYELVRVGYNELVGEIIRLEGDMATIQVYEETSGVTVGDPVLRTGKPLSVELGPGIMGSIFDGIQRPLKDIHDLTQSIYIPKGVNVPALSRTAQWDFSPINIKIGSHLTGGDIY 150
Rfer  1 MSKLAKIGDEEREGLFGYVHMVSGPVVTAEKMSGSAMYELVRVGYHELVGEIIRLEGDMATIQVYEETSGVTVGDPVLRTGKPLSVELGPGIMGSIFDGIQRPLKDINDLTQSIYIPKGVNVPALSRTAQWEFNPVNIKLGSHLTGGDIY 150
Atum  1 MSKI----DDEREGQYGYVHAVSGPVVTAEKMSGSAMYELVRVGYHELVGEIIRLEGDMATIQVYEETSGVTVGDPVLRTGKPLSVELGPGIMGSIFDGIQRPLKDINDLTQSIYIPKGVNVPALSRTAKWEFNPWNIKLGAHLTGGDIY 146
Btab  1 MSNLAKIRDEERESKFGYVFAVSGPVVTAEKMSGSAMYELVRVGYFELVGEIIRLEGDMATIQVYEETSGVTVGDPVLRTGKPLSVELGPGILGSIFDGIQRPLKDINEINQSIYIPKGVNIPALSKSHAWEFNPLNIKIGSHITGGDLY 150
Nlup  1 -MTLPKVKDVEKEGQYGYVFAVSGPVVTAEKMSGSAMYELVRVGHYELVGEIIRLEGDMATIQVYEETSGVTVGDPVLRTGKPLSVELGPGIMGSIFDGIQRPLKDINELSNSIYIPKGVNVPALSRTTAWEFHPLNIKVGSHITGGDEY 149
Bmor  1 KGGLRTIANEENEERFGYVFAVSGPVVTAEKMSGSAMYELVRVGYNELVGEIIRLEGDMATIQVYEETSGVTVGDPVLRTGKPLSVELGPGILGSIFDGIQRPLKDINELTQSIYIPKGINVPSLAREVDWEFNPLNVKVGSHITGGDLY 150
Harm  1 GGGLRTIADEEIEERFGFVFAVSGPVVTAEKMSGSAMYELVRVGYNELVGEIIRLEGDMATIQVYEETSGVTVGDPVLRTGKPLSVELGPGILGSIFDGIQRPLKDINELTQSIYIPKGVNVPSLARDVSWEFVPMNVKTGSHITGGDLY 150
Lmig  1 TSTLIKTSDEDRESKFGFVFAVSGPVVTAERMAGSAMYELVRVGYYELVGEIIRLEGDMATIQVYEDTSGVTVGDPVLRTGKPLSVELGPGIMGSIFDGIQRPLKDINELSNSIYIPKGVNVPALSRTAQWDFSPVSVKVGSHITGGDLY 150
Tdom  1 MSKLPLIRDEDRESKYGYVFAVSGPVVTAEHMAGSAMYELVRVGYSELVGEIIRLEGDMATIQVYEDTSGVTVGDPVLRTGKPLSVELGPGIMNSIFDGIQRPLQDINELTQSIYIPKGVNVPALSRTQKWDFTPHNVKVGSHITGGDIY 150
Gbim  1 SRNLAKLADEDREGKYGYVYAVSGPVVTAEHMSGSAMYELVRVGYFELVGEIIRLEGDLATIQVYEDTSGVTVGDPVLRTGKPLSVELGPGIMGSIFDGIQRPLKDIHELTHSIYIPKGVNVPALSRTEHWEFTPGDIRVGSHITGGDIY 150
Edan  1 SRGLPKIQNEEREKKFGYVFAVSGPVVTAERMAGSAMYELVRVGYSELVGEIIRLEGDMATIQVYEETSGVTVGDPVLRTGKPLSVELGPGIMGSIFDGIQRPLQDIHEISQSIYIPKGVNIPALNDKASWDFNPVHIKLGSHITGGDMY 150
Ibic  1 SRGLPKIQNEEREEKFGYVFAVSGPVVTAERMSGSAMYELVRVGYSELVGEIIRLEGDMATIQVYEETSGVTVGDPVLRTGKPLSVELGPGILGSIFDGIQRPLQDIHEIAQSIYIPKGVNIPALNDKSNWEFNPINIKMGSHITGGDVY 150
Dere  1 MSNLRKFKDEERESEYGRVYAVSGPVVTAEAMSGSAMYELVRVGYYELVGEIIRLEGDMATIQVYEETSGVTVGDPVLRTGKPLSVELGPGIMGSIFDGIQRPLRDIGVMTNSIYIPKGVNTTALSRSEMWEFNPLNVRVGSHITGGDLY 150
Llin  1 -MALPRIRDEEKESRFGYVFAVSGPVVTAEKMSGSAMYELVRVGYFELVGEIIRLEGDMATIQVYEETSGVTVGDPVLRTGKPLSVELGPGIMSSIFDGIQRPLKDICELTQSIYIPKGVNVPALSRSIAWDFTPSNIKVGAHITGGDLY 149
Ofas  1 -MALPRIRDEEHESKFGYVFAVSGPVVTAEKMSGSAMYELVRVGYFELVGEIIRLEGDMATIQVYEETSGVTVGDPVLRTGKPLSVELGPGILGSIFDGIQRPLKDINELSNSIYIPKGVNIPALSRSAKWDFSPLDVKVGSHITGGDLY 149
Aaeg  1 MSTLKKISDEDRESKFGYVFAVSGPVVTAERMSGSAMYELVRVGYYELVGEIIRLEGDMATIQVYEETSGVTVGDPVLRTGKPLSVELGPGIMGSIFDGIQRPLKDINELTSSIYIPEGCEHSLLVPYRRLGIQPLERKLGSHITGRDLY 150
Dpul  1 ASKLPKLEDEEKESNYGYVYAVSGPVVTAQKMSGSAMYELVRVGYSELVGEIIRLEGDMATIQVYEETSGVTVGDPVLRTGKPLSVELGPGIMESIFDGIQRPLKDINELTQSIYIPRGVNTPALSVTAEWDFKPANIRIGSHVTGGDIY 150
Dmag  1 AAKLPKLEDEEMESKYGYVYAVSGPVVTAQKMSGSAMYELVRVGYSELVGEIIRLEGDMATIQVYEETSGVTVGDPVLRTGKPLSVELGPGIMESIFDGIQRPLKDINELTQSIYIPRGVNTPALSVTAEWDFKPCNIRVGSHVTGGDIY 150
Smar  1 MEHLDKIGDVEKESIYGYVFAVSGPVVTANQMAGAAMYELVRVGHSELVGEIIRLEGDMATIQVYEETSGVTVGDPVLRTGKPLSVELGPGIMNNIFDGIQRPLQDIHDATQSIYIPRGVNIPALIRSKQWDFKPSNLKIGSHITGGDIY 150
Mmus  1 FSKLPKIRDEDKESTFGYVHGVSGPVVTACDMAGAAMYELVRVGHSELVGEIIRLEGDMATIQVYEETSGVSVGDPVLRTGKPLSVELGPGIMGAIFDGIQRPLSDISSQTQSIYIPRGVNVSALSRDIKWEFIPSNLRVGSHITGGDIY 150
Clup  1 FSKLPKILDEDKESTFGYVHGVSGPVVTACDMAGAAMYELVRVGHSELVGEIIRLEGDMATIQVYEETSGVSVGDPVLRTGKPLSVELGPGIMGAIFDGIQRPLSDISSQTQSIYIPRGVNVSALSRDVKWDFTPCNLRVGSHITGGDIY 150
Ggal  1 FSKLPKIRDEDREAFVGYVHGVSGPVVTACNMAGAAMYELVRVGHSELVGEIIRLEGDLATVQVYEETSGVSVGDPVLRTGKPLSVELGPGIMGAIFDGIQRPLSDISTLTKSIYIPRGVNVSALSRDVKWDFTPSNLRVGSHITGGDIY 150
Xtro  1 FSKLPKIRDEERESQFGYVHGVSGPVVTASNMAGAAMYELVRVGHCELVGEIIRLEGDMATIQVYEETSGVCVGDPVLRTGKPLSVELGPGIMGSIFDGIQRPLKDISDSTQSIYIPRGINVSALSKDIKWEFNPSSIRVGSHITGGDIY 150
Xlae  1 FSKLPKIRDEERESQFGYVHGVSGPVVTASNMAGAAMYELVRVGHSELVGEIIRLEGDMATIQVYEETSGVCVGDPVLRTGKPLSVELGPGIMGSIFDGIQRPLKDISDSTQSIYIPRGINVSALSKDIKWEFNPSSIRVGSHVTGGDIY 150
Cowc  1 NARKVKMSAEERADRFGYVFSVSGPVVVADHMNGAAMYELVRVGHDELVGEIIRLEGDTATIQVYEETSGVTVGDPVLRTGKPLSVELGPGLLASIYDGIQRPLRDISDLSNSIYIPRGVNTPALNKTAQWHFVPNKIRVGDHVTGGDQY 150
Rirr  1 KDNLPKIKDVERESEFGYVYGVSGPVVVAEQMNGSAMYELVRVGHEELVGEVIRIDGDKATIQVYEETSGVTVGDPVLKTGKPLSVELGPGLMSNIYDGIQRPLRSIREISDSIYIPRGIATPALDKKIQWDFEPLNYKVGDHITGGDIF 150
Gpro  1 LAKKRGPHDDPPESEYGYIFSVSGPVVVAEGMTGSAMYELVRVGHQELVGEVIRIEGDKATIQVYEETSGLTVGDPVLKTGKPLSVELGPGIMGDIFDGIQRPLKKIEEIAKGIFIPRGINTPSLDKKLNWEFEPQNFKVGDHITGGDVY 150
Apro  1 ----MGRDVETSERAYGYVRKVSGPVVVASSMSGSAMYELVRVGADKLIGEIIRLEGDTATVQVYEDTSGLTVGDPVTRNGKPLSVELGPGIMGTIFDGIQRPLKSIAIDSDSCFIPRGVDVPALDRTKAWQFDPLAFKVGDRITGGDIY 146
Pequ  1 GDRLTSFEDSEKESEYGYVRKVSGPVVVADGMGGAAMYELVRVGHDNLIGEIIRLEGDSATIQVYEETAGLTVNDPVLRTRKPLSVELGPGILGNIFDGIQRPLKTIAIKSGDVYIPRGVAVPALDKDFLWDFEPKKFSVGDHLTGGDLY 150
Pdac  1 SDRMTTFEDSEKESEYGYVRKVSGPVVVADGMGGAAMYELVRVGHDNLIGEIIRLEGDSATIQVYEETAGLTVNDPVLRTHKPLSVELGPGILGNIFDGIQRPLKTIALKSGDVYIPRGVSVPALDKDILWEFEPKKLAVGDLLTGGDLY 150
Obra  1 GDRITTFEDSEKESEYGYVRKVSGPVVVADGMGGAAMYELVRVGHDNLIGEIIRLEGDSATIQVYEETAGLMVNDPVLRTRKPLSVELGPGILGNIFDGIQRPLKTIAIKSGDVYIPRGVSVPALDKDVLWEFNPKNLGVGDAITGGDIY 150
```

65

Hsap 151 GLVHENTLVKHKLLLPPRAKGTVTYIAEPGNYTVDDVVLETEFDGERSKFTMLQVWPVRQPRPVTEKLPANYPLLTGQRVLDSLFPCVQGGTTAIPGAFGCGKTVISQSLSKYSNSDVIIYVGCGERGNEMSEVLRDFPQLSLEIDGVTE 300
Bger 151 GIVHENTLVKHKLLLPPRAKGTVTYIAPPGNYTVDDIVLETEFDGERSKFTMLQVWPVRQPRPVTEKLPANHPLLCGQRVLDSLFPCVQGGTTAIPGAFGCGKTVISQSLSKYSNSDVIIYVGCGERGNEMSEVLRDFPELTVEIEGNTE 300
Hpar 151 GIVHENTLVKHKLILPPRAKGTVVSIAPPGNYTVDDIVLETEFDGERTKFTMLQVWPVRQPRPVSEKLPANHPLLTGQRVLDSLFPCVQGGTTAIPGAFGCGKTVISQSLSKYSNSDVIIYVGCGERGNEMSEVLRDFPELTVEIDGQTE 300
Otau 151 GIVHENTLVKHKLILPPRAKGTVVSIAPPGNYTVDDIVLETEFDGERSKYTMLQVWPVRQPRPVSEKLPANHPLLTGQRVLDSLFPCVQGGTTAIPGAFGCGKTVISQSLSKYSNSDVIIYVGCGERGNEMSEVLRDFPELTVEIDGQTE 300
Tdic 151 GIVHENTLIKHKLILPPRAKGTVVSIAPPGNYTVDDVVLETEFDGEKTKYTMLQVWPVRQPRPVSEKLPANHPLLTGQRVLDSLFPCVQGGTTAIPGAFGCGKTVISQSLSKYSNSDVIVYVGCGERGNEMSEVLRDFPELSVEIDGQTE 300
Nves 150 GIVHENTLVKHKMILPPKAKGTVTYIASPGNYTVDDVVLETEFDGERQKFTMLQVWPVRQPRPVSEKLPANHPLLTGQRVLDSLFPCVQGGTTAIPGAFGCGKTVISQSLSKYSNSDVIIYVGCGERGNEMSEVLRDFPELTVEIDGVTE 299
Pcha 151 GIVHENTLVKHKMILPPRAKGTVTYIADPGNYTVDEVVLETEFDGERTKFTMLQVWPVRQPRPVTEKLPANHPLLTGQRVLDSLFPCVQGGTTAIPGAFGCGKTVISQSLSKYSNSDVIIYVGCGERGNEMSEVLRDFPELSVEIDGVTE 300
Apla 151 GIVHENTLVKHKLLLPPRSKGTVTYIASPGNYTVDDVILETEFDGERTKYTMLQVWPVRQPRPVAEKLPANHPLLTGQRVLDALFPCVQGGTTAIPGAFGCGKTVISQSLSKYSNSDVIIYVGCGERGNEMSEVLRDFPELTVEIEGATE 300
Sinv 151 GVVHENTLVKHWMILPPKCKGTVTYIAPIGNYTVDDVVLETEFDGERQKYTMLQVWPVRQPRPVTEKLPANHPLLTGQRVLDSLFPCVQGGTTAIPGAFGCGKTVISQALSKYSNSDVIIYVGCGERGNEMSEVLRDFPELTVEIEGVTE 300
Nvit 150 GIVHENTLVKHKMILPPKAKGTVTYIAPAGNYTVEDVVLETEFDGEKTKYTMLQVWPVRQPRPVTEKLPANHPLLTGQRVLDSLFPCVQGGTTAIPGAFGCGKTVISQALSKYSNSDVIIYVGCGERGNEMSEVLRDFPELTVEIDGVTE 299
Oora 120 GIVHENTLVKHRMILPPKSKGTVTYIAPAGNYTVEDIVMETEFDGEKTKYTMLQVWPVRQPRPVTEKLPANHPLLTGQRVLDSLFPCVQGGTTAIPGAFGCGKTVISQALSKYSNSDVIIYVGCGERGNEMSEVLRDFPELTVEIDGVTE 269
Amel 151 GVVYENTLVKHKMILPPKSKGTVTYIAPAGNYTVSDVILETEFDGERHKYTMLQVWPVRQPRPVTEKLPANHPLLTGQRVLDSLFPCVQGGTTAIPGAFGCGKTVISQALSKYSNSDVIIYVGCGERGNEMSEVLRDFPKLTVEIDGITE 300
Aros 151 GIVHENTLVKHKMILPPRSKGTVTYIAPPGNYTVNDIVLETEFDGEKNKFTMLQVWPVRQPRPVTEKLPANHPLLTGQRVLDSLFPCVQGGTTAIPGAFGCGKTVISQALSKYSNSDVIIYVGCGERGNEMSEVLRDFPELSVEIDGVTE 300
Dmel 151 GLVHENTLVKHKMIVNPRAKGTVRYIAPSGNYKVDDVVLETEFDGEITKHTMLQVWPVRQPRPVTEKLPANHPLLTGQRVLDSLFPCVQGGTTAIPGAFGCGKTVISQALSKYSNSDVIIYVGCGERGNEMSEVLRDFPELSVEIDGVTE 300
Dyak 151 GLVHENTLVKHKMIVNPRAKGTVRYIAPSGNYKVDDVVLETEFDGEITKHTMLQVWPVRQPRPVTEKLPANHPLLTGQRVLDSLFPCVQGGTTAIPGAFGCGKTVISQALSKYSNSDVIIYVGCGERGNEMSEVLRDFPELSVEIDGVTE 300
Ccap 151 GLVHENTLVKHKMIVNPRTKGTVRYIAPTGNYKVDDVVLETEFDGEITKHTMLQVWPVRQPRPVTEKLPANHPLLTGQRVLDSLFPCVQGGTTAIPGAFGCGKTVISQALSKYSNSDVIIYVGCGERGNEMSEVLRDFPELSVEIDGVTE 300
Bole 151 GLVHENTLVKHKMIVNPRTKGTVRYIAPTGNYKVDDVVLETEFDGEITKHTMLQVWPVRQPRPVTEKLPANHPLLTGQRVLDSLFPCVQGGTTAIPGAFGCGKTVISQALSKYSNSDVIIYVGCGERGNEMSEVLRDFPELSVEIDGVTE 300
Mdom 151 GLVHENTLVKHKMIVGPRSKGTVRYIAPAGNYHVDDVVLETEFDGEVTKHTMLQVWPVRQPRPVAEKLPANHPLLTGQRVLDSLFPCVQGGTTAIPGAFGCGKTVISQALSKYSNSDVIIYVGCGERGNEMAEVLGDFPELSVEIDGVTE 300
Lcup 151 GLVHENTLVKHKMIVGPRAKGTVRYIAPAGNYNVEDVVLETEFDGEITKHTMLQVWPVRQPRPVTEKLPANHPLLTGQRVLDSLFPCVQGGTTAVPGAFGCGKTVISQALSKYSNSDVIIYVGCGERGNEMSEVLRDFPELSVEIDGVTE 300
Cqui 151 GLVHENTLVKHKLLVPPRAKGTVRYIAPPGNYTVEDIILETEFDGEVNKYSMLQVWPVRQPRPVTEKMPANHPLLTGQRVLDSLFPCVQGGTTAIPGAFGCGKTVISQALSKYSNSDVIVYVGCGERGNEMSEVLRDFPELSVEIDGVTE 300
Agam 151 GIVHENTLVKHKLMVPPRAKGTVKYIAPSGNYTVDDVILETEFDGEISKFTMLQVWPVRQPRPVTEKLPANHPLLTGQRVLDSLFPCVQGGTTAIPGAFGCGKTVISQSLSKYSNSDVIVYVGCGERGNEMSEVLRDFPELSVEIDGVTE 300
Fcep 149 GVVHENTLVKHKILLPPRAKGTVTYIAEPGNYTVDDVILETEFDGEKSKFTMLQVWPVRQPRPVTEKLPANYPLLTGQRVLDSLFPCVQGGTTAIPGAFGCGKTVISQALSKYSNSDVIIYVGCGERGNEMSEVLRDFPELSVEIDGQTE 298
Ldec 149 GMVHENTLVKHKMILPPKSKGTVTYVAEPGNYTVDEVVLETEFDGERSKYTMLQVWPVRQARPVSEKLPANHPLLTGQRVLDSLFPCVQGGTTAIPGAFGCGKTVISQSLSKYSNSDVIVYVGCGERGNEMSEVLRDFPELTVEIGGETE 298
Agla 151 GIVHENTLVKHKMLLPPRAKGTVTYVADPGNYSVDEVVLETEFDGDRTRFTMLQVWPVRQPRPVSEKLPANHPLLTGQRVLDSLFPCVQGGTTAIPGAFGCGKTVISQSLSKYSNSDVIVYVGCGERGNEMSEVLRDFPELTVEIGGETE 300
Dvir 150 GLVHENTLVKHKMILPPRAKGTVTYIAEPGNYTVDEVVLETEFDGDRTKYTMLQVWPVRQARPVSEKLPANHPLLTGQRVLDALFPCVQGGTTAIPGAFGCGKTVISQSLSKYSNSDVIIYVGCGERGNEMSEVLRDFPELTVEIDGHTE 299
Haxy 151 GIVHENTLVKQKLMLPPKSKGTVTYVAEPGSYTVDDIVLETEFDGERSKYTMLQVWPVRQPRPVSEKLPANHPLLTGQRVLDSLFPCVQGGTTAIPGAFGCGKTVISQSLSKYSNSDVIIYVGCGERGNEMSEVLRDFPELTVEIEGQTE 300
Hvig 151 GLVHENTLVKQKLMLPPKSKGTVTYIAEPGSYTVDDIVLETEFDGERSKYTMLQVWPVRQPRPVSEKLPANHPLLTGQRVLDSLFPCVQGGTTAIPGAFGCGKTVISQSLSKYSNSDVIVYVGCGERGNEMSEVLRDFPELTVEIEGQTE 300
Dpon 151 GMVHENTLVKHKLILPPKAKGTITYIAEPGNYTVDDIVLETEFDGERTKFTMLQVWPVRQPRPCSEKLQANHPLLTGQRVLDALFPCVQGGTTAIPGAFGCGKTVISQALSKYSNSDVIVYVGCGERGNEMSEVLRDFPELTVEIEGATE 300
Rfer 151 GMVHENTLVKQKLILPPKAKGTVRFIAEPGNYTVNDVVLETEFDGERTEYTMLQVWPVRQPRPVSEKLPANHPLLTGQRVLDSLFPCVQGGTTAIPGAFGCGKTVISQALSKYSNSDVIIYVGCGERGNEMSEVLRDFPELTVEIDGHTE 300
Atum 147 GIVHENTLVKHKIVLPPKAKGTVTYVAEPGNYTVDEVVLETEFDGERTKYSMLQVWPVRQPRPVSEKLPANHPLLTGQRVLDSLFPCVQGGTTAIPGAFGCGKTVISQSLSKYSNSDVIIYVGCGERGNEMSEVLRDFPELSVEIDGHTE 296
Btab 151 GIVFENTLVKHKLMLPPKAKGTVTYIAPPGNYKVDDIVLETEFDGEQSKYTMLQIWPVRQPRPVTEKLPANHPLLTGQRVLDSLFPCVLGGTTAIPGAFGCGKTVISQSLSKYSNSDVIVYVGCGERGNEMSEVLRDFPELTVEIDGVTE 300
Nlup 150 AIVHENTLVKHKLILPPRSKGTVKYLAPPGNYTVDDIVLETEFDGERSKFTMLQVWPVRQPRPVTEKLPANYPLLTGQRVLDSLFPCVQGGTTAIPGAFGCGKTVISQALSKYSNSDVIVYVGCGERGNEMSEVLRDFPELTVEIDGVTE 299
Bmor 151 GIVHENTLVKHRMLVPPKAKGTVTYIAPAGNYKVTDVVLETEFDGERQKYSMLQVWPVRQPRPVTEKLPANHPLLTGQRVLDSLFPCVQGGTTAIPGAFGCGKTVISQALSKYSNSDVIIYVGCGERGNEMSEVLRDFPELTVEIEGVTE 300
Harm 151 GLVHENTLVKHRMLIPPKAKGTVTYIAPAGNYKVTDVVLETEFDGEKEKYTMLQVWPVRQPRPVTEKLPANHPLLTGQRVLDSLFPCVQGGTTAIPGAFGCGKTVISQALSKYSNSDVIVYVGCGERGNEMSEVLRDFPELTVEIEGVTE 300
Lmig 151 GLVHENTLVKHKLLLPPRAKGTVTYIAEPGNYTVDDVVLETEFDGERSKFTMLQVWPVRQPRPVTEKLPANYPLLTGQRVLDSLFPCVQGGTTAIPGAFGCGKTVISQSLSKYSNSDVIIYVGCGERGNEMSEVLRDFPELSVEIDGVTE 300
Tdom 151 GMVFENTLIKHKVMLPPRAKGTVRYIAEPGSYDINDVILETEFDGERSKYTMLQVWPVRQPRPVTEKLPANHPLLTGQRVMDSLFPCVQGGTTAIPGAFGCGKTVISQSLSKFSNSDVIIYVGCGERGNEMSEVLRDFPELTTEIEGVTE 300
Gbim 151 GVVFENTLVKHKIMLPPRSKGTVQYVAPAGNYTVEDVVLETEFDGERTKYTMLQVWPVRQPRPVTEKLPANYPLLCGQRVLDSLFPCVQGGTTAIPGAFGCGKTVISQSLSKYSNSDIIVYVGCGERGNEMSEVLRDFPELTIEVDGKTE 300
Edan 151 GIVHENTLVKHKLLLPPRDKGTVTYIAEPGSYTVKDVILETEFDGERKKYTMLQVWPVRQPRPCTEKLPANYPLLTGQRVLDSLFPCVQGGTTAIPGAFGCGKTVISQSLSKYSNSDVIVYVGCGERGNEMSEVLRDFPELTCEIDGVTE 300
Ibic 151 GLVHENTLVKHKLMLPPRDKGTVTYVAEPGSYTVKDVVLETEFDGERTKYTMLQVWPVRQPRPVTEKLPANWPLLTGQRVLDSLFPCVQGGTTAIPGAFGCGKTVISQSLSKYSNSDVIIYVGCGERGNEMSEVLRDFPELTCEIDGVTE 300
Dere 151 GVVHENTLVKQRMIVAPRAKGTVRYIAPAGNYNLEDIVLETEFDGEITKHTMLQVWPVRQPRPVTEKLPANHPLFTGQRVLDSLFPCVQGGTTAIPGAFGCGKTVISQALSKYSNSDVIIYVGCGERGNEMSEVLRDFPELTCEIDGVTE 300
Llin 150 AVVHENTLVKQKMIMPARGRGTVKYIAPPGNYTVDDVVMETEFDGEKTEIKMLQVWPVRQPRPVAEKLPANYPLLTGQRVLDALFPCVQGGTTAIPGAFGCGKTVISQALSKYSNSDVIIYVGCGERGNEMSEVLRDFPELTVEIDGVTE 299
Ofas 150 GVVHENTLVKHKMILPPRAKGTVTYLAPPGSYTVDDVILETEFDGEKSEFTMLQVWPVRQPRPVTEKLPANYPLLTGQRVLDALFPCVQGGTTAIPGAFGCGKTVISQALSKYSNSDVIIY------------VLRDFPELSVEIDGVTE 287
Aaeg 151 GLVHENTLVKHKLLVPPRAKGTVRYIAPPGNYTVDDIILETEFDGEINKWSMLQVWPVRQPRPVTEKLPANHPLLTGQRVLDSLFPCVQGGTTAIPGAFGCGKTVISQALSKYSNSDVIIYVGCGERGNEMSEVLRDFPELSVEIDGVTE 300
Dpul 151 GTVFENRMVKHKMMLPPRARGTVTWVASPGHYRVTDKILQIEFDGEKTDYTMLQVWPVRQPRPVVEKLPANYPLLTGQRVLDALFPCVQGGTTAIPGAFGCGKTVISQSLSKYSNSDVIIYVGCGERGNEMSEVLRDFPELTCEVDGITE 300
Dmag 151 GTVFENRMVKHKMMLPPRARGTVTWIASPGHYHVTDKILEVEFDGEKTGYSMLQVWPVRQPRPVVEKLPANYPLLTGQRVLDALFPCVQGGTTAIPGAFGCGKTVISQSLSKYSNSDVIIYVGCGERGNEMSEVLRDFPELTCEVDGITE 300
Smar 151 GIVFENTLIKHKVLLPPKARGSILYIAPPGNYDITDTILETEFDGERQKFSMLQVWPVRQPRPVTEKLPANYPLLTGQRVLDTLFPCVQGGTTAIPGAFGCGKTVISQALSKYSNSDVIIYVGCGERGNEMSEVLRDFPELTTEIDGVNE 300
Mmus 151 GIVNENSLIKHKIMLPPRNRGSVTYIAPPGNYDASDVVLELEFEGVKEKFSMVQVWPVRQVRPVTEKLPANHPLLTGQRVLDALFPCVQGGTTAIPGAFGCGKTVISQSLSKYSNSDVIIYVGCGERGNEMSEVLRDFPELTMEVDGKVE 300
Clup 151 GIVNENSLIKHKIMLPPRNRGTVTYIAPPGNYDTSDVVLELEFEGIKEKFSMVQVWPVRQVRPVTEKLPANHPLLTGQRVLDALFPCVQGGTTAIPGAFGCGKTVISQSLSKYSNSDVIIYVGCGERGNEMSEVLRDFPELTMEVDGKVE 300
Ggal 151 GVVNENSLIKHKIMLPPRNRGTVTYIAPPGNYDTSDVVLELEFEGVKEKFTMVQVWPVRQVRPVTEKLPANHPLLTGQRVLDALFPCVQGGTTAIPGAFGCGKTVISQSLSKYSNSDVIIYVGCGERGNEMSEVLRDFPELTMEVDGKVE 300
Xtro 151 GAVMENSLIRHKIMLPPRNRGTVTYIAPPGHYDTSDVVLELEFEGVKDKFTMVQVWPVRSIRPVTEKLPANHPLLTGQRVLDALFPCVQGGTTAIPGAFGCGKTVISQSLSKYSNSDVIIYVGCGERGNEMSEVLRDFPELTMEVDGKTE 300
Xlae 151 GMVMENSLIRHKIMLPPRNRGTVTYIAPPGHYDTSDVVLELEFEGVKDKFTMVQLWPVRSIRPVTEKLPANHPLLTGQRVLDALFPCVQGGTTAIPGAFGCGKTVISQSLSKYSNSDVIIYVGCGERGNEMSEVLRDFPELTMEVDGKVE 300
Cowc 151 GTVKENSLIEHRILVPPLARGTVKFIASEGDYSLEDVVLETEFAGKASQFTMMQVWPVRTRRPDAEKLPANYPLLTGQRVLDALFPCVQGGTTAIPGAFGCGKTVISQALSKFSNSDVIIYVGCGERGNEMSEVLRDFPELEMEVDGRKE 300
Rirr 151 GKVYENSLVNHNIMLSPKALGTITHISGKGSYTLEDIVLETEFEGQKSKHTMMQIWPVRSPRPVTEKLAANYPLLTGQRVLDALFPCVQGGTTAIPGAFGCGKTVISQSLSKYSNSDIIIYVGCGERGNEMAEVLMDFPELSINIDGRQE 300
Gpro 151 GIVRENTLIRHRIMLPPKSRGTITYIAPKGNYTLKDEVLEIEFEGEKSKYTMLQIWAVRSPRPVTEKIAGDHPLLTGQRVLDGLFPCVQGGTTAIPGAFGCGKTVISQSLSKYSNSDVIVYVGCGERGNEMAEVLMEFPELTMTVEGREE 300
Apro 147 GEVHENTLMQHKILLPPGARGTVTYIAPAGEYTITDPVIEVEFGGAKKEYSMLQLWPVRAPRPVAQKLLANTPLLTGQRVLDGLFPGVLGGTCAIPGAFGCGKTVISQALSKYSNSDGIIYVGCGERGNEMAEVLMDFPALTMTLDGREE 296
Pequ 151 ATVFENTLMQHHVALPPGAMGKISYIAPSGQYSLKDTVLEIEFQGVKKEYTMLQTWPVRSPRPVASKLAADTPLLTGQRVLDALFPSVLGGTCAIPGAFGCGKTVISQALSKYSNSDIVVYVGCGERGNEMAEVLMDFPQLTMTLDGREE 300
Pdac 151 ATVFENTLMQHHVALPPGSMGKITYVAPAGHYSLRDTVLELEFQGVKKQFTMLQTWPVRTPRPVAGKLAADTPLLTGQRVLDALFPSVLGGTCAIPGAFGCGKTVISQALSKYSNSDTVVYVGCGERGNEMAEVLMDFPQLTMTLDGREE 300
Obra 151 AAVHENTLMEHRVALPPGSMGKISYIAPAGQYSLQDTVLELEFQGIKKQFTMLQTWPVRSPRPVASKLAADTPLLTGQRVLDALFPSVLGGTCAIPGAFGCGKTVISQALSKYSNSEAVVYVGCGERGNEMAEVLMDFPQLTMTYEGREE 300

66

```
Hsap 301 SIMKRTALVANTSNMPVAAREASIYTGITLSEYFRDMGYNVSMMADSTSRWAEALREISGRLAEMPADSGYPAYLGARLASFYERAGRVKCLGNPDREGSVSIVGAVSPPGGDFSDPVTTATLGIVQVFWGLDKKLAQRKHFPSINWLIS 450
Bger 301 SIMKRTALVANTSNMPVAAREASIYTGITLSEYFRDMGYNVSMMADSTSRWAEALREISGRLAEMPADSGYPAYLGARLASFYERAGRVKCLGNPDREGSVSIVGAVSPPGGDFSDPVTSATLGIVQVFWGLDKKLAQRKHFPSINWLIS 450
Hpar 301 SIMKRTALVANTSNMPVAAREASIYTGITLSEYFRDMGYNVSMMADSTSRWAEALREISGRLAEMPADSGYPAYLGARLASFYERAGRVKCLGNPDREGSVSIVGAVSPPGGDFSDPVTSATLGIVQVFWGLDKKLAQRKHFPSINWLIS 450
Otau 301 SIMKRTALVANTSNMPVAAREASIYTGITLSEYFRDMGYNVSMMADSTSRWAEALREISGRLAEMPADSGYPAYLGARLASFYERAGRVKCLGNPDREGSVSIVGAVSPPGGDFSDPVTSATLGIVQVFWGLDKKLAQRKHFPSINWLIS 450
Tdic 301 SIMKRTALVANTSNMPVAAREASIYTGITLSEYFRDMGYNVSMMADSTSRWAEALREISGRLAEMPADSGYPAYLGARLASFYERAGRVKCLGNPDREGSVSIVGAVSPPGGDFSDPVTSATLGIVQVFWGLDKKLAQRKHFPSINWLIS 450
Nves 300 SIMKRTALVANTSNMPVAAREASIYTGITLSEYFRDMGYNVSMMADSTSRWAEALREISGRLAEMPADSGYPAYLGARLASFYERAGRVKCLGNPDREGSVSIVGAVSPPGGDFSDPVTSATLGIVQVFWGLDKKLAQRKHFPSINWLIS 449
Pcha 301 SIMKRTALVANTSNMPVAAREASIYTGITLSEYFRDMGYNVSMMADSTSRWAEALREISGRLAEMPADSGYPAYLGARLASFYERAGRVKCLGNPDREGSVSIVGAVSPPGGDFSDPVTSATLGIVQVFWGLDKKLAQRKHFPSINWLIS 450
Apla 301 SIMKRTALVANTSNMPVAAREASIYTGITLSEYFRDMGYNVSMMADSTSRWAEALREISGRLAEMPADSGYPAYLGARLASFYERAGRVKCLGNPDREGSVSIVGAVSPPGGDFSDPVTSATLGIVQVFWGLDKKLAQRKHFPSINWLIS 450
Sinv 301 SIMKRTALVANTSNMPVAAREASIYTGITLSEYFRDMGYNVSMMADSTSRWAEALREISGRLAEMPADSGYPAYLGARLASFYERAGRVKCLGNPDREGSVSIVGAVSPPGGDFSDPVTSATLGIVQVFWGLDKKLAQRKHFPSINWLIS 450
Nvit 300 SIMKRTALVANTSNMPVAAREASIYTGITLSEYFRDMGYNVSMMADSTSRWAEALREISGRLAEMPADSGYPAYLGARLASFYERAGRVKCLGNPDREGSVSIVGAVSPPGGDFSDPVTSATLGIVQVFWGLDKKLAQRKHFPSINWLIS 449
Oora 270 SIMKRTALVANTSNMPVAAREASIYTGITLSEYFRDMGYNVSMMADSTSRWAEALREISGRLAEMPADSGYPAYLGARLASFYERAGRVKCLGNPDREGSVSIVGAVSPPGGDFSDPVTSATLGIVQVFWGLDKKLAQRKHFPSINWLIS 419
Amel 301 SIMKRTALVANTSNMPVAAREASIYTGITLSEYFRDMGYNVSMMADSTSRWAEALREISGRLAEMPADSGYPAYLGARLASFYERAGRVKCLGNPDREGSVSIVGAVSPPGGDFSDPVTSATLGIVQVFWGLDKKLAQRKHFPSINWLIS 450
Aros 301 SIMKRTALVANTSNMPVAAREASVYTGITLSEYFRDMGYNVSMMADSTSRWAEALREISGRLAEMPADSGYPAYLGARLASFYERAGRVQCLGNPDREGSVSIVGAVSPPGGDFSDPVTTATLGIVQVFWGLDKKLAQRKHFPSINWLIS 450
Dmel 301 SIMKRTALVANTSNMPVAAREASIYTGITLSEYFRDMGYNVSMMADSTSRWAEALREISGRLAEMPADSGYPAYLGARLASFYERAGRVKCLGNPEREGSVSIVGAVSPPGGDFSDPVTSATLGIVQVFWGLDKKLAQRKHFPSINWLIS 450
Dyak 301 SIMKRTALVANTSNMPVAAREASIYTGITLSEYFRDMGYNVSMMADSTSRWAEALREISGRLAEMPADSGYPAYLGARLASFYERAGRVKCLGNPEREGSVSIVGAVSPPGGDFSDPVTSATLGIVQVFWGLDKKLAQRKHFPSINWLIS 450
Ccap 301 SIMKRTALVANTSNMPVAAREASIYTGITLSEYFRDMGYNVSMMADSTSRWAEALREISGRLAEMPADSGYPAYLGARLASFYERAGRVKCLGNPEREGSVSIVGAVSPPGGDFSDPVTSATLGIVQVFWGLDKKLAQRKHFPSINWLIS 450
Bole 301 SIMKRTALVANTSNMPVAAREASIYTGITLSEYFRDMGYNVSMMADSTSRWAEALREISGRLAEMPADSGYPAYLGARLASFYERAGRVKCLGNPEREGSVSIVGAVSPPGGDFSDPVTSATLGIVQVFWGLDKKLAQRKHFPSINWLIS 450
Mdom 301 SIMKRTALVANTSNMPVAAREASIYTGITLSEYFRDMGYNVSMMADSTSRWAEALREISGRLAEMPADSGYPAYLGARLASFYERAGRVKCLGNPEREGSVSIVGAVSPPGGDFSDPVTSATLGIVQVFWGLDKKLAQRKHFPSINWLIS 450
Lcup 301 SIMKRTALVANTSNMPVAAREASIYTGITLSEYFRDMGYNVSMMADSTSRWAEALREISGRLAEMPADSGYPAYLGARLASFYERAGRVKCLGNPEREGSVSIVGAVSPPGGDFSDPVTSATLGIVQVFWGLDKKLAQRKHFPSINWLIS 450
Cqui 301 SIMKRTALVANTSNMPVAAREASIYTGITLSEYFRDMGYNVSMMADSTSRWAEALREISGRLAEMPADSGYPAYLGARLASFYERAGRVKCLGNPEREGSVSIVGAVSPPGGDFSDPVTSATLGIVQVFWGLDKKLAQRKHFPSINWLIS 450
Agam 301 SIMKRTALVANTSNMPVAAREASIYTGITLSEYFRDMGYNVSMMADSTSRWAEALREISGRLAEMPADSGYPAYLGARLASFYERAGRVKCLGNPEREGSVSIVGAVSPPGGDFSDPVTSATLGIVQVFWGLDKKLAQRKHFPSINWLIS 450
Fcep 299 SIMKRTALVANTSNMPVAAREASIYTGITLSEYFRDMGYNVSMMADSTSRWAEALREISGRLAEMPADSGYPAYLGARLASFYERAGRVKCLGNPDREGSVSIVGAVSPPGGDFSDPVTSATLGIVQVFWGLDKKLAQRKHFPSINWLIS 448
Ldec 299 SIMKRTALVANTSNMPVAAREASIYTGITLSEYFRDMGYNVSMMADSTSRWAEALREISGRLAEMPADSGYPAYLGARLASFYERAGRVKCLGNPDREGSVSIVGAVSPPGGDFSDPVTSATLGIVQVFWGLDKKLAQRKHFPSINWLIS 448
Agla 301 SIMKRTALVANTSNMPVAAREASIYTGITLSEYFRDMGYNVSMMADSTSRWAEALREISGRLAEMPADSGYPAYLGARLASFYERAGRVKCLGNPDREGSVSIVGAVSPPGGDFSDPVTSATLGIVQVFWGLDKKLAQRKHFPSINWLIS 450
Dvir 300 SIMKRTALVANTSNMPVAAREASIYTGITLSEYFRDMGYNVSMMADSTSRWAEALREISGRLAEMPADSGYPAYLGARLASFYERAGRVKCLGNPDREGSVSIVGAVSPPGGDFSDPVTTATLGIVQVFWGLDKKLAQRKHFPSVDWLGS 449
Haxy 301 SIMKRTALVANTSNMPVAAREASIYTGITLSEYFRDMGYNVSMMADSTSRWAEALREISGRLAEMPADSGYPAYLGARLASFYERAGRVKCLGNPDREGSVSIVGAVSPPGGDFSDPVTSATLGIVQVFWGLDKKLAQRKHFPSINWLIS 450
Hvig 301 SIMKRTTLVANTSNMPVAAREASIYTGITLSEYFRDMGYNVSMMADSTSRWAEALREISGRLAEMPADSGYPAYLGARLASFYERAGRVKCLGNPDREGSVSIVGAVSPPGGDFSDPVTSATLGIVQVFWGLDKKLAQRKHFPSINWLIS 450
Dpon 301 SIMKRTALVANTSNMPVAAREASIYTGITLSEYFRDMGYNVSMMADSTSRWAEALREISGRLAEMPADSGYPAYLGARLASFYERAGRVKCLGNPDREGSVSIVGAVSPPGGDFSDPVTSATLGIVQVFWGLDKKLAQRKHFPSVNWLIS 450
Rfer 301 SIMKRTALVANTSNMPVAAREASIYTGITLSEYFRDMGYNVSMMADSTSRWAEALREISGRLAEMPADSGYPAYLGARLASFYERAGRVKCLGNPDREGSVSIVGAVSPPGGDFSDPVTSATLGIVQVFWGLDKKLAQRKHFPSVNWLIS 450
Atum 297 SIMKRTALVANTSNMPVAAREASIYTGITLSEYFRDMGYNVSMMADSTSRWAEALREISGRLAEMPADSGYPAYLGARLASFYERAGRVKCLGNPDREGSVSIVGAVSPPGGDFSDPVTSATLGIVQVFWGLDKKLAQRKHFPSINWLIS 446
Btab 301 SIMKRTALVANTSNMPVAAREASIYTGITLSEYFRDMGYNVSMMADSTSRWAEALREISGRLAEMPADSGYPAYLGARLASFYERAGRVKCLGNPDREGSVSIVGAVSPPGGDFSDPVTSATLGIVQVFWGLDKKLAQRKHFPSINWLIS 450
Nlup 300 SIMKRTALVANTSNMPVAAREASIYTGITLSEYFRDMGYNVSMMADSTSRWAEALREISGRLAEMPADSGYPAYLGARLASFYERAGRVKCLGNPDREGSVSIVGAVSPPGGDFSDPVTSATLGIVQVFWGLDKKLAQRKHFPSINWLIS 449
Bmor 301 SIMKRTALVANTSNMPVAAREASIYTGITLSEYFRDMGYNVSMMADSTSRWAEALREISGRLAEMPADSGYPAYLGARLASFYERAGRVKCLGNPDREGSVSIVGAVSPPGGDFSDPVTAATLGIVQVFWGLDKKLAQRKHFPSINWLIS 450
Harm 301 SIMKRTALVANTSNMPVAAREASIYTGITLSEYFRDMGYNVSMMADSTSRWAEALREISGRLAEMPADSGYPAYLGARLASFYERAGRVKCLGNPDREGSVSIVGAVSPPGGDFSDPVTAATLGIVQVFWGLDKKLAQRKHFPSINWLIS 450
Lmig 301 SIMKRTALVANTSNMPVAAREASIYTGITLSEYFRDMGYNVSMMADSTSRWAEALREISGRLAEMPADSGYPAYLGARLASFYERAGRVKCLGNPDREGSVSIVGAVSPPGGDFSDPVTTATLGIVQVFWGLDKKLAQRKHFPSINWLIS 450
Tdom 301 SIMKRTALVANTSNMPVAAREASIYTGITLSEYFRDMGYNVSMMADSTSRWAEALREISGRLAEMPADSGYPAYLGARLASFYERAGRVKCLGNPEREGSVSIVGAVSPPGGDFSDPVTSATLGIVQVFWGLDKKLAQRKHFPSINWLIS 450
Gbim 301 SIMKRTALVANTSNMPVAAREASIYTGITLSEYFRDMGYNVSMMADSTSRWAEALREISGRLAEMPADSGYPAYLGARLASFYERAGRTKCLGNPEREGSVSIVGAVSPPGGDFSDPVTSATLGIVQVFWGLDKKLAQRKHFPSINWLIS 450
Edan 301 SIMKRTALVANTSNMPVAAREASIYTGITLSEYFRDMGYNVSMMADSTSRWAEALREISGRLAEMPADSGYPAYLGARLASFYERAGRVKCLGNPEREGSVSIVGAVSPPGGDFSDPVTSATLGIVQVFWGLDKKLAQRKHFPSINWLIS 450
Ibic 301 SIMKRTALVANTSNMPVAAREASIYTGITLSEYFRDMGYNVSMMADSTSRWAEALREISGRLAEMPADSGYPAYLGARLASFYERAGRVKCLGNPEREGSVSIVGAVSPPGGDFSDPVTSATLGIVQVFWGLDKKLAQRKHFPSINWLIS 450
Dere 301 SIMKRTALVANTSNMPVAAREASIYTGITLSEYFRDMGYNVAMMADSTSRWAEALREISGRLAEMPADSGYPAYLGARLATFYERAGRVKCLGNPEREGSVSIVGAVSPPGGDFSDPVTSATLGIVQVFWGLDKKLAQRKHFPSINWLIS 450
Llin 300 SIMKRTALVANTSNMPVAAREASIYTGITLSEYFRDMGYNVSMMADSTSRWAEALREISGRLAEMPADSGYPAYLGARLASFYERAGRVKCLGSPDREGSVSIVGAVSPPGGDFSDPVTSATLGIVQVFWGLDKKLAQRKHFPSINWLIS 449
Ofas 288 SIMKRTALVANTSNMPVAAREASIYTGITLSEYFRDMGYNVSMMADSTSRWAEALREISGRLAEMPADSGYPAYLGARLASFYERAGRVKCLGNPEREGSVSIVGAVSPPGGDFSDPVTSATLGIVQVFWGLDKKLAQRKHFPSINWLIS 437
Aaeg 301 SIMKRTALVANTSNMPVAAREASIYTGITLSEYFRDMGYNVSMMADSTSRWAEALREISGRLAEMPADSGYPAYLGARLASFYERAGRVKCLGNPEREGSVSIVGAVSPPGGDFSDPVTSATLGIVQVFWGLDKKLAQRKHFPSINWLIS 450
Dpul 301 SIMKRTSLVANTSNMPVAAREASIYTGITLSEYFRDMGYNVSMMADSTSRWAEALREISGRLAEMPADSGYPAYLGARLASFYERAGRVKCQGSPEREGSVSIVGAVSPPGGDFSDPVTSATLGIVQVFWGLDKKLAQRKHFPSVNWLIS 450
Dmag 301 SIMKRTSLVANTSNMPVAAREASIYTGITLSEYFRDMGYNVSMMADSTSRWAEALREISGRLAEMPADSGYPAYLGARLASFYERAGRVKCQGSPEREGSVSIVGAVSPPGGDFSDPVTSATLGIVQVFWGLDKKLAQRKHFPSVNWLIS 450
Smar 301 SIMKRTALVANTSNMPVAAREASIYTGITLSEYFRDMGYNVSMMADSTSRWAEALREISGRLAEMPADSGYPAYLGARLASFYERAGRVKCLGNPEREGR------FSPPGGDFSDPVTSATLGIVQVFWGLDKKLAQRKHFPSVNWLIS 444
Mmus 301 SIMKRTALVANTSNMPVAAREASIYTGITLSEYFRDMGYHVSMMADSTSRWAEALREISGRLAEMPADSGYPAYLGARLASFYERAGRVKCLGNPEREGSVSIVGAVSPPGGDFSDPVTSATLGIVQVFWGLDKKLAQRKHFPSVNWLIS 450
Clup 301 SIMKRTALVANTSNMPVAAREASIYTGITLSEYFRDMGYHVSMMADSTSRWAEALREISGRLAEMPADSGYPAYLGARLASFYERAGRVKCLGNPEREGSVSIVGAVSPPGGDFSDPVTSATLGIVQVFWGLDKKLAQRKHFPSVNWLIS 450
Ggal 301 SIMKRTALVANTSNMPVAAREASIYTGITLSEYFRDMGYHVSMMADSTSRWAEALREISGRLAEMPADSGYPAYLGARLASFYERAGRVKCLGNPEREGSVSIVGAVSPPGGDFSDPVTSATLGIVQVFWGLDKKLAQRKHFPSVNWLIS 450
Xtro 301 SIMKRTALVANTSNMPVAAREASIYTGITLSEYFRDMGYHVSMMADSTSRWAEALREISGRLAEMPADSGYPAYLGARLASFYERAGRVKCLGNPEREGSVSIVGAVSPPGGDFSDPVTSATLGIVQVFWGLDKKLAQRKHFPSVNWLIS 450
Xlae 301 SIMKRTALVANTSNMPVAAREASIYTGITLSEYFRDMGYHVSMMADSTSRWAEALREISGRLAEMPADSGYPAYLGARLASFYERAGRVKCLGNPEREGSVSIVGAVSPPGGDFSDPVTSATLGIVQVFWGLDKKLAQRKHFPSVNWLIS 450
Cowc 301 TIMKRTTLVANTSNMPVAAREASIYTGITLSEYFRDMGYNVSMMADSTSRWAEALREISGRLAEMPADSGYPAYLGARLASFYERAGKVKCLGNPSREGSVSIVGAVSPPGGDFSDPVTSATLGIVQVFWGLDKKLAQRKHFPSVNWLMS 450
Rirr 301 SIMKRTTLVANTSNMPVAAREASIYTGITLSEYFRDQGKNVAMMADSTSRWAEALREISGRLAEMPADSGYPAYLGARLASFYERAGKVTCLGNPKRDGSVTVVGAVSPPGGDFSDPVTSSTLGIVQVFWGLDKKLAQRKHFPSVNWSLS 450
Gpro 301 SIMLRTTLVANTSNMPVAAREASIYTGITLAEYFRDQGMNVAMMADSTSRWAEALREISGRLAEMPADSGYPAYLGARLASFYERSGKVICLGSPKRNGSVTIVGAVSPPGGDFSDPVTSSTLSIVQVFWGLDKKLAQRKHFPSVNWNIS 450
Apro 297 SIMQRTTLVANTSNMPVAAREASIYTGITLAEYFRDMGYNFSMMADSTSRWAEALREISGRLAEMPADSGYPAYLSARLASFYERAGRVKCLGSPAREGSVTIVGAVSPPGGDFSDPVTTATLSIVQVFWGLDKKLAQRKHFPSVNWLIS 446
Pequ 301 SVMKRTTLVANTSNMPVAAREASIYTGITIAEYFRDMGYNVSMMADSTSRWAEALREISGRLAEMPADSGYPAYLAARLASFYERAGKVKCLGGPDRTGSVTIVGAVSPPGGDFSDPVTSATLGIVQVFWGLDKKLAQRKHFPSVNWLIS 450
Pdac 301 SVMKRTTLVANTSNMPVAAREASIYTGITIAEYFRDMGYNVSMMADSTSRWAEALREISGRLAEMPADSGYPAYLAARLASFYERAGKVKCLGGPDRTGSVTIVGAVSPPGGDFSDPVTSATLGIVQVFWGLDKKLAQRKHFPSVNWLIS 450
Obra 301 SVMKRTTLVANTSNMPVAAREASIYTGITIAEYFRDMGYNVSMMADSTSRWAEALREISGRLAEMPADSGYPAYLAARLASFYERAGKVRCLGSPDRDGSVTIVGAVSPPGGDFSDPVTSATLSIVQVFWGLDKKLAQRKHFPSVNWLIS 450
```

```
Hsap 451 YSKYMRALDDFYDKNFPEFVPLRTKVKEILQEEEDLSEIVQLVGKASLAETDKITLEVAKLLKDDFLQQNSYSPYDRFCPFYKTVGMLKNMIAFYDMSRHAVESTAQSENKITWNVIRDSMGNILYQLSSMKFKDPVKDGEAKIKADFEQ 600
Bger 451 YSKYMRALDDFYDKNFPEFVPLRTKVKEILQEEEDLSEIVQLVGKASLAETDKITLEVAKLLKDDFLQQNSYSPYDRFCPFYKTVGMLKNIIAFYDMSRHAVESTAQSENKITWGVIRDSMGNILYQLSSMKFKDPVKDGEAKIKADFEQ 600
Hpar 451 YSKYTRALDDFYDRNFAEFVSLRTKVKEILQEEEDLSEIVQLVGKASLAESDKITLEVAKLLKEDFLQQNSYSSYDRFCPFYKTVGMLKNMIGLYDMSRHAVETTAQSENKITWNVIRDSMGNILYQLSSMKFKDPVKDGEAKIKADFDQ 600
Otau 451 YSKYTRALDDFYDKNFAEFVALRTKVKEILQEEEDLSEIVQLVGKASLAESDKITLEVAKLLKEDFLQQNSYSAYDRFCPFYKTVGMLKNMIGLYDMARHAVESTAQSENKITWNVIRDSMGNILYQLSSMKFKDPVKDGEAKIKADFDQ 600
Tdic 451 YSKYTRALDDFYDKNFAEFVPLRTKVKEILQEEEDLSEIVQLVGKASLAESDKITLEVAKLLKEDFLQQNSYSTYDRFCPFYKTVGMLKNMIGLYDMARHAVETTAQSENKITWNVIRDSMGNILYQLSSMKFKDPVKDGEAKIKADFDQ 600
Nves 450 YSKYTRALDDFYDKNFAEFVSLRTKVKEILQEEEDLSEIVQLVGKASLAETDKITLEVAKLLKEDFLQQNSYSSYDRFCPFYKTVGMLRNMIGLYDMSRHAVESTAQSENKITWNVIRDSMGNILYQLSSMKFKDPVKDGEAKIKADFEQ 599
Pcha 451 YSKYMRALDDFYDKNFVEFVPLRTKVKEILQEEEDLSEIVQLVGKASLAETDKITLEVAKLLKDDFLQQNSYSSYDRFCPFYKTVGMLRNMIAFYDMSRHAVETTAQSENKITWNVIRDSMGNILYQLSSMKFKDPVKDGEAKIKADFDQ 600
Apla 451 YSKYIRALDDFYDKNFAEFVPLRTKVKEILQEEEDLSEIVQLVGKASLAETDKITLEVAKLLKEDFLQQNSYSSYDRFCPFYKTVGMLRNMIGFYDMSRHAVESTAQSDNKITWNIIRDAMGNILYQLSSMKFKDPVKDGEAKIKSDFDQ 600
Sinv 451 YSKYLRALDDFYDKNFPEFVPLRTKVKEILQEEEDLSEIVQLVGKASLAETDKITLEVAKLLKDDFLQQNSYSPYDRFCPFYKTVGMLRNMIAFYDMARHAVESTAQSDNKITWNVIRDSMGNILYQLSSMKFKDPVKDGEAKIRADFDQ 600
Nvit 450 YSKYLRALDDFYDKNFQEFVPLRTKVKEILQEEEDLSEIVQLVGKASLAETDKITLEVAKLLKDDFLQQNSYSPYDRFCPFYKTVGMLRNMIAFYDMARHAVESTAQSDNKITWNVIRDSMGNILYQLSSMKFKDPVKDGEAKIRSDFDQ 599
Oora 420 YSKYLRALDDFYDKNFQEFVPLRTKVKEILQEEEDLSEIVQLVGKASLAETDKITLEVAKLLKDDFLQQNSYSPYDRFCPFYKTVGMLRNMIAFYDMARHAVESTAQSDNKITWNVIRDGMGNILYQLSSMKFKDPVKDGEAKIRSDFDQ 569
Amel 451 YSKYLRALDDFYDKNFAEFVPLRTKVKEILQEEEDLSEIVQLVGKASLAETDKITLEVAKLLKDDFLQQNSYSPYDRFCPFYKTVGMLRNMIAFYDMARHAVESTAQSDNKITWNVIKDSMYNILYQLSSMKFKDPVQDGEAKIRADFDQ 600
Aros 451 YSNYMRALDDFYDKNFQEFVPLRTKVKEILQEEEDLSEIVQLVGKASLAETDKITLEVAKLLKDDFLQQNSYSAYDRFCPFYKTVGMLRNMIAFYDMARHAVESTAQSDNKITWNVIRDSMGNILYQLSSMKFKDPVKDGEAKIRADFEQ 600
Dmel 451 YSKYMRALDDFYDKNFPEFVPLRTKVKEILQEEEDLSEIVQLVGKASLAETDKITLEVAKLLKDDFLQQNSYSSYDRFCPFYKTVGMLRNIIDFYDMARHSVESTAQSENKITWNVIREAMGNIMYQLSSMKFKDPVKDGEAKIKADFEQ 600
Dyak 451 YSKYMRALDDFYDKNFPEFVPLRTKVKEILQEEEDLSEIVQLVGKASLAETDKITLEVAKLLKDDFLQQNSYSSYDRFCPFYKTVGMLRNIIDFYDLARHSVESTAQSENKITWNVIREAMGNIMYQLSSMKFKDPVKDGEAKIKADFEQ 600
Ccap 451 YSKYMRALDDFYEKNFAEFVPLRTKVKEILQEEEDLSEIVQLVGKASLAETDKITLEVAKLLKDDFLQQNSYSSYDRFCPFYKTVGMLKNIISFYDMARHSVESTAQSENKITWNVIREAMGNIMYQLSSMKFKDPVKDGEAKIKADFEQ 600
Bole 451 YSKYMRALDDFYEKNFAEFVPLRTKVKEILQEEEDLSEIVQLVGKASLAETDKITLEVAKLLKDDFLQQNSYSSYDRFCPFYKTVGMLKNIISFYDLARHSVESTAQSENKITWNVIREAMGNIMYQLSSMKFKDPVKDGEAKIKGDFEQ 600
Mdom 451 YSKYMRALDDFYDKNFPEFVPLRTKVKEILQEEEDLSEIVQLVGKASLAETDKITLEVAKLLKDDFLQQNSYSSYDRFCPFYKTVGMLKNIIAFYDMARHSVESTAQSENKITWNVIREAMGNIMYQLSSMKFKDPVKDGEAKIKADFEQ 600
Lcup 451 YSKYMRALDDFYDKNFPEFVPLRTKVKEILQEEEDLSEIVQLVGKASLAETDKITLEVAKLLKDDFLQQNSYSSYDRFCPFYKSVGMLKNIIAFYDLARHSVESTAQSENKITWNVIREAMGNIMYQLSSMKFKDPVKDGEAKIKADFEQ 600
Cqui 451 YSKYMRALDDFYDKNFQEFVPLRTKVKEILQEEEDLSEIVQLVGKASLAETDKITLEVAKLLKDDFLQQNSYSAYDRFCPFYKTVGMLRNIIGFYDMARHAVETTAQSENKITWNVIREAMGNILYQLSSMKFKDPVKDGEAKIKADFDQ 600
Agam 451 YSKYMRALEDFYDKNFPEFVPMRTKVKEILQEEEDLSEIVQLVGKASLAETDKITLEVAKLLKDDFLQQNSYSAYDRFCPFYKTVGMLKNMIGFYDMARHAVETTAQSENKITWNVIRDSMGQILYQLSSMKFKDPVKDGEPKIKADFDQ 600
Fcep 449 YSKYMRALDDFYDKNFPEFVPLRTKVKEILQEEEDLSEIVQLVGKASLAEADKITLEVAKLLKDDFLQQNSYSAYDRFCPFYKTSGMLKNMIAFYDMARHAVESTAQSENKITWNVIRDSMGQIMYQLSSMKFKDPHKDGEAKIKADFEQ 598
Ldec 449 YSKYMRALDDFYDKNFPEFVSLRTKVKEILQEEEDLSEIVQLVGKASLAEADKITLEIAKLLKEDFLQQNSYSSYDRFCPFYKTVGMLKNMIGLYDMSRHAVETTAQSENKITWNVIRDSMSNILYQLSSMKFKDPVKDGEAKIKADFDQ 598
Agla 451 YSKYTRALDDFYDKNFPEFVSLRTKVREILQEEEDLSEIVQLVGKASLAEADKITLEIAKLLKEDFLQQNSYSSYDRFCPFYKTVGMLKNMIGLYDMSRHAVETTAQSENKITWTVIRDSMSNILYQLSSMKFKDPVKDGEAKIKADFDQ 600
Dvir 450 YSKYLRALDDFYDKNFQEFIPLRTKVKEILQEEDDLAEIVQLVGKASLAETDKITLEIARLLKEDFLQQNSYSSYDRFCPFYKTVGMLRNMIGLYDMARHAVESTAQSENKITWNVIRDSMSGILYQLSSMKFKDPVKDGEAKIKADFDQ 599
Haxy 451 YSKYTRALDDFYDKNFAEFVPLRTKVKEILQEEEDLSEIVQLVGKASLAETDKITLEVAKLLKEDFLQQNSYSSYDRFCPFYKTVGMLKNMIGLYDMARHAVETTAQSENKITWNVIREAMNNILYQLSSMKFKDPVKDGEAKIKADFDQ 600
Hvig 451 YSKYTRALDDFYDKNFPEFVPLRTKVKEILQEEEDLSEIVQLVGKASLAETDKITLEVAKLLKEDFLQQNSYSAYDRFCPFYKTVGMLKNMIGLYDMSRHAVETTAQSENKITWNVIREAMNNILYQLSSMKFKDPVKDGEAKIKAEFDQ 600
Dpon 451 YSKYMRALDDFYDKNFPEFVPLRIKAKEILQEEEDLSEIVQLVGKASLAEGDKITLEVAKLLKEDFLQQNSYSAYDRFCPFYKTVGMLKNMIGLYDMARHAVEITAQSENKITWSVIRDSMSNILYQLSSMKFKDPVKDGEAKIKADFDQ 600
Rfer 451 YSKYMRALDDFYDKNFSEFVPLRTKAKEILQEEEDLSEIVQLVGKASLAETDKITLEVAKLLKEDFLQQNSYSSYDRFCPFYKTVGMLKNMIGLYDMARHAVESTAQSENKITWAVIRDSMSNILYQLSSMKFKDPVKDGEAKIKADFDQ 600
Atum 447 YSKYTRALDDFYDKNFPEFVPLRTKVKEILQEEEDLSEIVQLVGKASLAETDKITLEVAKLLKEDFLQQNSYSAYDRFCPFYKTVGMLRNMIGLYDMARHAVESTAQSENKITWNVIRDSMSNILYQLSSMKFKDPVKDGEAKIKADFDQ 596
Btab 451 YSKYTRALDDFYDKNFPEFVALRSKVKEILQEEEDLSEIVQLVGKASLAESDKITLEVAKLLKDDFLQQNSYSPYDRFCPFYKTVGMLRNMIAFYDMSRHAVESTAQSENKISWSVIKDSMGNILYQLSSMKFKDPVKDGEQKIKADFDQ 600
Nlup 450 YSKYMRALDDFYDKNYAEFVPLRTKVKEILQEEEDLSEIVQLVGKASLAESDKITLEVAKVLKDDFLQQNSYSAYDRFCPFYKTVGMLKNMIAFYDMARHAVESTAQSENKITWSVIKDSMGNILYQLSSMKFKDPQKEGETKLKADFEQ 599
Bmor 451 YSKYMRALDDFYEKNYPEFVPLRTKVKEILQEEEDLSEIVQLVGKASLAETDKITLEVAKLLKDDFLQQNSYSSYDRFCPFYKTVGMLKNIITFYDMSRHAVESTAQSDNKVTWNVIRDAHGHVLYQLSSMKFKDPVKDGEPKIKADFDQ 600
Harm 451 YSKYMRALDDFYDKNYVEFVPLRTKVKEILQEEEDLSEIVQLVGKASLAETDKITLEVAKLLKDDFLQQNSYSAYDRFCPFYKTVGMLKNIIAFYDMSRHAVESTAQSDNKVTWNVIRDAMGNVLYQLSSMKFKDPVKDGEAKIKADFDQ 600
Lmig 451 YSKYMRALDDFYDKNFPEFVPLRTKVKEILQEEEDLSEIVQLVGKASLAETDKITLEVAKLLKDDFLQQNSYSPYDRFCPFYKTVGMLKNMIAFYDMSRHAVESTAQSENKITWNVIRDSMGNILYQLSSMKFKE--RSGSNSFDHNTVT 598
Tdom 451 YSKYMRALDDFYEKNFPEFVPLRTKVKEILQEEEDLSEIVQLVGKASLAETDKITLEVARLIKDDFLQQNGYTPYDRFCPFYKTVGMLKNIVAFYDMSRHAVESTAQSENKITWAIIRESMGNILYQLSSMKFKDPVKDGEAKIKADFEQ 600
Gbim 451 YSKYLRALDDFYEKNYPEFVPLRTKVKEILQEEEDLSEIVQLVGKASLAETDKITLEVAKLLKDDFLQQNSYSPYDRFCPFYKTVGMLKNMISFYDMARHAVESTAQTENKITWNIIRENMSNIIYQLSSMKFKDPVKDGEQKIKADFEQ 600
Edan 451 YSKYMRALDDYYDKNFPEFVSLRTKVKEILQEEEDLSEIVQLVGKASLAETDKITLEVAKLLKDDFLQQNSYSAYDRFCPFYKTVGMLRNMMGLYDMSRHAVESTAQSENKITWSTIKDSMGAILYQLSSMKFKDPQKEGEAKIKADFDQ 600
Ibic 451 YSKYMRALDDYYDKNFPEFVTLRTKVKEILQEEEDLSEIVQLVGKASLAETDKITLEVAKLLKDDFLQQNSYSPYDRFCPFYKTVGMLRNIMGLYDMARHAVESTAQSENKITWSTIKDSMGSILYQLSSMKFKDPVKDGEAKIKADFDQ 600
Dere 451 YSKYMRALDEYYDKNYPEFVPLRTKVKEILQEEEDLSEIVQLVGKASLAETDKVTLEVAKLLKDDFLQQNSYSPYDRVCPFYKTVGMLRNIMAFYETARHAVETTAQSDNKITWNTIRESMGGIMYQLSSMKFKDPVKDGEQKIKADYDQ 600
Llin 450 YSKYMRALDDFYDKRYPEFVPLRTKVKEILQEEEDLAEIVQLVGKGSLAESDKITLEIAKILKDDFLQQNSYSPYDRFCPFYKTVGMLKNMISFYDLARHTVESTAQSDNKITWTVIKESMGNILYQLSSMKFKDPVKDGEAKIKADFEQ 599
Ofas 438 YSKYMRALDDFYDKNFPEFVPLRTKVKEILQEEEDLSEIVQLVGKGSLAESDKITLEIAKLLKDDFLQQNSYSPYDRFCPFYKTVGMLKNMIAFYDAARHSCETTAQSEKKITWSVIKESMGNILYQLSSMKFKDPVKDGEAKIRSDFEQ 587
Aaeg 451 YSKYMRALDDFYDKNFQEFVPTAYKVKEILQEEEDLSEIVQLVGKASLAETDKITLEVAKLLKDDFLQQNSYSAYDRFCPFYKTVGMLRNMIGFYDMARHAVETTAQSENKITWNVIRDSMGNILYQLSSMKFKDPGRMAK-KIKADFDQ 599
Dpul 451 YSKYMRALDDHYERHFPEFVPLRTKVKEILQEEEDLSEIVQLVGKASLAETDKITLEVAKLLKDDFLQQNGYSAYDRFCPFYKTVGMLRNMVGFYDMARHAVESTAQSDNKITWGTIRESMGNILYQLSSMKFKDPVKDGEDKIKQDYDE 600
Dmag 451 YSKYMRALDDHYERHFPEFVPLRTKVKEILQEEEDLSEIVQLVGKASLAETDKITLEVAKLLKDDFLQQNGYSAYDRFCPFYKTVGMLRNMVGFYDMARHAVESTSQSDNKITWGTIRESMGNILYQLSSMKFKDPVKDGEAKIKQDYDE 600
Smar 445 YSKYMRALDEFYEKSFPEFVPLRTKCKEILQEEEDLSEIVQLVGKASLAETDKITLEVAKLIKDDFLQQNGYSAYDRFCPFYKSVGMMKNMIAFYDLSRHAVETTSQSDNKITWAIIRESMGDVLYQLSTMKFKDPIAEGETKIKADYEE 594
Mmus 451 YSKYMRALDEYYDKHFTEFVPLRTKAKEILQEEEDLAEIVQLVGKASLAETDKITLEVAKLIKDDFLQQNGYTPYDRFCPFYKTVGMLSNMISFYDMARRAVETTAQSDNKITWSIIREHMGEILYKLSSMKFKDPVKDGEAKIKADYAQ 600
Clup 451 YSKYTRALDEYYDKHFTEFVPLRTKAKEILQEEEDLAEIVQLVGKASLAETDKITLEVAKLIKDDFLQQNGYTPYDRFCPFYKTVGMLSNMIAFYDMARRAVETTAQSDNKITWSIIREHMGEILYKLSSMKFKDPVKDGEAKIKADYAQ 600
Ggal 451 YSKYTRALDEYYDKHFTEFVPLRTKAKEILQEEEDLAEIVQLVGKASLAETDKITLEVAKLIKDDFLQQNGYTPYDRFCPFYKTVGMLSNMIAFYDMARRAVETTAQSDNKITWSIIRENMSEILYRLTSMKFKDPVKDGETKIKADYAQ 600
Xtro 451 YSKYTRALDEYYDKHFQEFVPLRTKAKEILQEEEDLAEIVQLVGKASLAETDKITLEVAKLIKDDFLQQNGYTPYDRFCPFYKTVGMLYNMIAFYDMARRAVDTTAQSDNKITWSIIREHMGEILYRLTSMKFKDPVKEGEAKIKADYAQ 600
Xlae 451 YSKYTRALDEYYDKHFPEFVPLRTKAKEILQEEEDLAEIVQLVGKASLAESDKITLEVAKLIKDDFLQQNGYTPYDRFCPFYKTVGMLYNMIAFYDMARRAVDTTAQSDNKITWSIIREHMGEILYRLTSMKFKDPVKEGEAKIKADYAQ 600
Cowc 451 YSKYERALDPFYEDFDPEFVSFRTKCKEILQLEEDLSEIVQLVGKGSLAESDKITLEVAKIIRDDFLQQNGYSNYDRFCPFYKTVGMMRNICLFFDLARRAVESTAQSDNKITFAIIKDKMGGLMHELSSMKFLDPI-NGEASLKESFKQ 599
Rirr 451 YSKYTKVLETFYESNEPDFIALRDKTKEILQKEEDLSEIVQLVGKSALGEADKITLEIAKIIKDDFLQQNGYSSYDRYCPFYKTTWLLKNMIGFYNSATHAVEVT---NNEITWAKIRESMNDLIYKLSSMKFEDPA-EGETALRERYQQ 596
Gpro 451 YSKYTKALDPWYEQFDPEFTSLRTKCKEILQMEEDLSEIVQLVGKSALAETDKITLEIAKIIKEDFLQQNGYSSYDRYCPFFKTVGMLRNMMGFYDLATHAVEASS---NQITWAKIRESMSDILYKLSSMKFEDPI-EGEAVIRERYQK 596
Apro 447 YSKYTKGLEPFYDKFDPDFVSVRKSAREILQKEDDLNEIVQLVGKDALAEGDKITLETARLIKDDFLQQNSFTPYDKYCPFYKSVDMLRNMIAFHNAANAAVERSAASDSKLTFNVIKTRLSDVLYKLSSQKFEDPA-E-EEALRAKFAA 594
Pequ 451 YSKYTKALESFYEKFDPDFIDIRTKAREVLQREDDLNEIVQLVGKDALAETDKIILETAKLLREDYLAQNAFTPYDKFCPFYKSVWMMRNIIHFHTLANQAVERAAGSDQKITYSVIKHHLGDLFYRLVSQKFEDPA-EGEEALSAKFKK 599
Pdac 451 YSKYSKALESFYEKFDPDFIDIRTKAREVLQREDDLNEIVQLVGKDALAETDKIILETAKLLREDYLAQNAFTPYDKFCPFYKSVWMMRNIIHFNTLANQAVERGAGSEQKITYSVIKHRLGDLFYRLVSQKFEDPS-EGEEVLVDKFKK 599
Obra 451 YSKYSKALESFYEKFDPDFIDIRTKAREVLQREDDLNEIVQLVGKDALAESDKITLETAKLLREDYLAQNAFTSYDKYCPFYKSVWMMRNIIHFNTLANQAVERAAGVDQKITYSVIKHRMGDLFYRLVSQKFEDPA-EGEEVLVVKFQK 599
```

```
Hsap 601 LHEDIQQAFRNLED 614
Bger 601 LHEDIQQAFRNLED 614
Hpar 601 LYEDIQQAFRNLED 614
Otau 601 LYEDIQQAFRNLED 614
Tdic 601 LYEDIQQAFRNLED 614
Nves 600 LYEDIQQAFRNLED 613
Pcha 601 LYEDIQQAFRNLED 614
Apla 601 LYEDMQQTFRNLED 614
Sinv 601 LHEDIQQAFRNLED 614
Nvit 600 LHEDIQQAFRNLED 613
Oora 570 LHEDIQQAFRNLED 583
Amel 601 LHEDIQQAFRNLED 614
Aros 601 LHEDIQQAFRNLED 614
Dmel 601 LHEDLQQAFRNLED 614
Dyak 601 LHEDLQQAFRNLED 614
Ccap 601 LHEDLQQAFRNLED 614
Bole 601 LHEDLQQAFRNLED 614
Mdom 601 LHEDLQQAFRNLED 614
Lcup 601 LHEDLQQAFRNLED 614
Cqui 601 LYEDLQQAFRNLED 614
Agam 601 LYEDMQQAFRNLED 614
Fcep 599 LHEDLQQAFRNLED 612
Ldec 599 LYEDIQQAFRNLED 612
Agla 601 LYEDIQQAFRNLED 614
Dvir 600 LYEDIQQAFRNLED 613
Haxy 601 LYEDIQQAFRNLED 614
Hvig 601 LYEDIQQAFRNLED 614
Dpon 601 LYEDIQQAFRNLED 614
Rfer 601 LYEDIQQAFRNLED 614
Atum 597 LYEDIQQAFRNLED 610
Btab 601 LHEDIQQAFRNLED 614
Nlup 600 LHEDIQQAFRNLED 613
Bmor 601 LLEDMSAAFRNLED 614
Harm 601 LLEDMAAAFRNLED 614
Lmig 599 LKYKLDNEFELLED 612
Tdom 601 LNEEIQQAFRNLED 614
Gbim 601 LHEDMQQAFRNLED 614
Edan 601 LYEDIQQAFRNLED 614
Ibic 601 LYEDIQQAFRNLED 614
Dere 601 LYEDLQQAFRNLED 614
Llin 600 LHEDMQQAFRNLED 613
Ofas 588 LYEDIQQAFRNLED 601
Aaeg 600 LYEDLQQAFRNLED 613
Dpul 601 LHEQMQQAFRNLED 614
Dmag 601 LHEQMQQAFRNLED 614
Smar 595 LHERMQQEFRNLED 608
Mmus 601 LLEDMQNAFRSLED 614
Clup 601 LLEDMQNAFRSLED 614
Ggal 601 LFEDMQNAFRSLED 614
Xtro 601 LLEDMQNAFRSLED 614
Xlae 601 LLEDMQNAFRSLED 614
Cowc 600 LAEKLQNAFRNLED 613
Rirr 597 LQTEIQERFRQLLE 610
Gpro 597 LYKEMEERFRQMLD 610
Apro 595 LSDEIRDRFRTLEE 608
Pequ 600 LYEDLTIGFRNLED 613
Pdac 600 LYDDLTAGFRNLED 613
Obra 600 LYDDLTTGFRNLED 613
```

**Fig. S8. Multiple alignments of amino acid of V-ATPase A used to phylogenetic analysis.**

Amino acid sequences were obtained from 57 eukaryotes. 614 residues were used to phylogenetic reconstruction. Reconstructed phylogenetic trees show in Fig. S4b. Datasets used in the phylogenetic analysis show in Table S7.

75

```
Dmel            1 MALSDADVQKQIKHMMAFIEQEANEKAEEIDAKAEEEFNIEKGRLVQQQRLKIMEYYEKKEKQVELQKKIQSSNMLNQARLKVLKVREDHVSSVLDDARKRLGEVTKNQSEYETVLTKLIVQGLFQIMEPKVILRCREVDVPLVRNVLPA 150
Dyak            1 MALSDADVQKQIKHMMAFIEQEANEKAEEIDAKAEEEFNIEKGRLVQQQRLKIMEYYEKKEKQVELQKKIQSSNMLNQARLKVLKVREDHVSSVLDDARKRLGEVTKNQSEYQTVLTKLIVQGLFQIMEPKVILRCREVDVPLVRDVLPN 150
Dere            1 MALSDADVQKQIKHMMAFIEQEANEKAEEIDAKAEEEFNIEKGRLVQQQRLKIMEYYEKKEKQVELQKKIQSSNMLNQARLKVLKVREDHVSSVLDDARKRLGEVTQKQSEYQTVLTKLIVQGLFQIMEPKVILRCREVDVPLVRDVLPT 150
Dvir            1 MALSDADVQKQIKHMMAFIEQEANEKAEEIDAKAEEEFNIEKGRLVQQQRLKIMEYYEKKEKQVELQKKIQSSNMLNQARLKVLKVREDHVGSVLDDARKRLGEVTQNESEYKTVLEKLIVQGLYQVMEPKVTLRCRQVDVPLVRAVIPS 150
Mdom            1 MALSDADVQKQIKHMMAFIEQEANEKAEEIDAKAEEEFNIEKGRLVQQQRLKIMEYYEKKEKQVELQKKIQSSNMLNQARLKVLKVREDHVASVLEDARRRLGEVTKNPAEYKVVLEKLILQALFQTMEPTVILRCRQVDVGLVNEVLPA 150
Lcup            1 MALSDADVQKQIKHMMAFIEQEANEKAEEIDAKAEEEFNIEKGRLVQQQRLKIMEYYEKKEKQVELQKKIQSSNMLNQARLKVLKIREDHVASVLEDARRRLGEVTKNPAEYKVVLQKLILQALYQTMEPTVILRCRQVDVGLVNEALPV 150
Bdor            1 MALSDADVQKQIKHMMAFIEQEANEKAEEIDAKAEEEFNIEKGRLVQQQRLKIMEYYEKKEKQVELQKKIQSSNMLNQARLKVLKVREDHVRNVLDEARRRLGEVTKNPSEYKEVLQKLILQALYQAMEPSVTLRCRQADIGLINEVLPA 150
Ccap            1 MALSDADVQKQIKHMMAFIEQEANEKAEEIDAKAEEEFNIEKGRLVQQQRLKIMEYYEKKEKQVELQKKIQSSNMLNQARLKVLKVREDHVRNVLDEARRRLGEVTKNPGEYKEVLQKLILQALYQAMEQNVTLRCRQADIGLINEVLPA 150
Agam            1 MALSDADVQKQIKHMMAFIEQEANEKAEEIDAKAEEEFNIEKGRLVQQQRLKIMEYYEKKEKQVELQKKIQSSNMLNQARLKVLKVREDHVSNVLDECRRRLGEVTKDPARYGEILTALITQGLLQLMEAKVLIRGRQADAQVIQNVLPA 150
Bmor            1 MALSDADVQKQIKHMMAFIEQEANEKAEEIDAKAEEEFNIEKGRLVQQQRLKIMEYYEKKEKQVELQKKIQSSNMLNQARLKVLKVREDHVRNVLDEARKRLAEVPKDTKLYSELLVTLIVQALFQLMEPTVTIRVRQTDKALVESLLGK 150
Lexo            1 MALSDADVQKQIKHMMAFIEQEANEKAEEIDAKAEEEFNIEKGRLVQQQRLKIMEYYEKKEKQVELQKKIQSSNMLNQARLKVLKVREDHVRNVLDEARKRLAEVPSDAKLYSDLLVTLIVQALFQLMESTVTLRVRQADKSLVESLLGK 150
Osca            1 MALSDADVQKQIKHMMAFIEQEANEKAEEIDAKAEEEFNIEKGRLVQQQRLKIMEYYEKKEKQVELQKKIQSSNMLNQARLKVLKVREDHVRNVLDEARKRLAEVPSDAKLYSDLLVTLIVQALFQLMESTVTLRVRQADKSLVESLLGK 150
Harm            1 MALSDADVQKQIKHMMAFIEQEANEKAEEIDAKAEEEFNIEKGRLVQQQRLKIMEYYEKKEKQVELQKKIQSSNMLNQARLKVLKVREDHVRNLLDEARKRLAEVPNDQKMYSELLTTLIVQALFQIMEASVTLRVRQADKALVDSVLGK 150
Aros            1 MALSDADVQKQIKHMMAFIEQEANEKAEEIDAKAEEEFNIEKGRLVQQQRLKIMEFFEKKEKQVELQKKIQSSNMLNQARLKVLKVREDHVRNVLDEARRRLGEVTRDQSRYSQVLKSLIVQSLYQLMEANIVVRVRQNDVALTESILAT 150
Amel            1 MALSDADVQKQINHMMAFIEQEANEKAEEIDAKAEEEFNIEKGRLVQQQRLKIMEYYEKKEKQVELQKKIQSSNMLNQARLKVLKVREDHVRNVLDEARKRLGEVTRDISRYREILKLLIVQGLCQLTENHVTIRVRQVDLPLVESLLDS 150
Sinv            1 MALSDADVQKQIKHMMAFIEQEANEKAEEIDAKAEEEFNIEKGRLVQQQRLKIMEYYEKKEKQVELQKKIQSSNMLNQARLKALKVREDHVRNVLDDARKRLSEVTHNSTRYKEVLQLLIIQGLYQLTESNITLRVRQVDLPLVESIIDS 150
Nvit            1 MALSDADVQKQIKHMMAFIEQEANEKAEEIDAKAEEEFNIEKGRLVQQQRLKIMEYYEKKEKQVELQKKIQSSNMLNQARLKVLKVREDHVRNVLDDARKKLGEVARDQGKYAEILKLLITQGLYQLIELNVVVRARQADHDLIESIMPA 150
Haxy            1 MALSDADVQKQIKHMMAFIEQEANEKAEEIDAKAEEEFNIEKGRLVQQQRLKIMEYYEKKEKQVELQKKIQSSNMLNQARLKVLKVREDHVRNVLEDARKRLGDVTRDQGRYREILESLILQGLYQLFESNVVIRVRPQDRDLVRSITSN 150
Hvig            1 MALSDADVQKQIKHMMAFIEQEANEKAEEIDAKAEEEFNIEKGRLVQQQRLKIMEYYEKKEKQVELQKKIQSSNMLNQARLKVLKVREDHVRNVLEEARKRLGDVTRDRKKYAEILESLILQGLYQLFESHVLVRVRPQDRDIVHSIIAN 150
Sjap            1 MALSDADVQKQIKHMMAFIEQEANEKAEEIDAKAEEEFNIEKGRLVQQQRLKIMEYYEKKEKQVELQKKIQSSNMLNQARLKVLKVREDHVRNVLEEARKRLGDVTRDQAKYRVILESLILQALYQLFESHVLLRVRPQDRELVQSILPK 150
Agla            1 MALSDADVQKQIKHMMAFIEQEANEKAEEIDAKAEEEFNIEKGRLVQQQRLKIMEYYEKKEKQVELQKKIQSSNMLNQARLKVLKVREDHVRSVLDEARKRLGEVTRDQGKYAQILESLILQGLYQLFEKEVTIRVRPQDRDLVKSVLPV 150
Achi            1 MALSDADVQKQIKHMMAFIEQEANEKAEEIDAKAEEEFNIEKGRLVQQQRLKIMEYYEKKEKQVELQKKIQSSNMLNQARLKVLKVREDHVRSVLDEARKRLGEVTRDQGKYAQILESLILQGLYQLFEKEVTIRVRPQDRDLVKSVLPV 150
Ldec            1 MALSDVDVQKQIKHMMAFIEQEANEKAEEIDAKAEEEFNIEKGRLVQQQRLKIMEYYEKKEKQVELQKKIQSSNMLNQARLKVLKVREDHVRTVLEEARKRLGQVTNDQGKYSQILESLILQGLYQLFEKDVTIRVRPQDRELVKSIIPT 150
Dpon            1 MALSDADVQKQIKHMMAFIEQEANEKAEEIDAKAEEEFNIEKGRLVQQQRLKIMEYYEKKEKQVELQKKIQSSNMLNQARLKVLKVREDHVRSVLEDARKRLGEVTRDQGRYAQIAESLILQALYQLFENNVIVRTRPQDRDLVKTVLPT 150
Tcas            1 MALSDVDVQKQIKHMMAFIEQEANEKAEEIDAKAEEEFNIEKGRLVQQQRLKIMEYYEKKEKQVELQKKIQSSNMLNQARLKVLKVREDHVHNVLDDARKRLGEITNDQARYSQLLESLILQSLYQLFENNIVVRVRQQDRSIIQGILPV 150
Xbas            1 MALSDADVQKQIKHMMAFIEQEANEKAEEIDAKAEEEFNIEKGRLVQQQRLKIMEYYEKKEKQVELQKKIQSSNMLNQARLKVLKVREDHVRSVLDEARKRLGQVTRDQTKYADVLEQLILQGLYQLFEEKVTVRVREEDRSLVENVLPI 150
Apla            1 MALSDADVQKQIKHMMAFIEQEANEKAEEIDAKAEEEFNIEKGRLVQQQRLKIMEYYEKKEKQVELQKKIQSSNMLNQARLKVLKVREDHVYQVLDDARKRLGEVTRNTPKYSEILKSLILQGLYQLFETNVSVRVRAQDKALVESLLPD 150
Nves            1 MALSDSDVQKQIKHMMAFIEQEANEKAEEIDAKAEEEFNIEKGRLVQQQRLKIMEYYEKREKQVELQKKIQSSNMLNQARLKVLKVREDHVRNVLDEARKRLGQVTSNHSKYQEIIKTLILQGLYQLFETNVLVRCRAQDKDLVEGCFDD 150
Otau            1 MALSDADVQKQIKHMMAFIEQEANEKAEEIDAKAEEEFNIEKGRLVQQQRLKIMEYYEKKEKQVELQKKIQSSNMLNQARLKVLKVREDHVRNVLEEARKRLGEVTNNSAKYKEILSTLILQALYQLFETNVSVRCRTQDKSLVESVFSE 150
Obor            1 MALSDADVQKQIKHMMAFIEQEANEKAEEIDAKAEEEFNIEKGRLVQQQRLKIMEYYEKKEKQVELQKKIQSSNMLNQARLKVLKVREDHVRNVLEDARKRLGEVTSNSLKYKEILSTLILQALFQLFEPAVQIRCRTQDRGLVESALPE 150
Tdic            1 MALSDADVQKQIKHMMAFIEQEANEKAEEIDAKAEEEFNIEKGRLVQQQRLKIMEYYEKKEKQVELQKKIQSSNMLNQARLKVLKVREDHVRSVLEDARKRLGEVTSNSAKYKEILSTLILQALFQLFESTVHIRCRSQDRGLVESALPE 150
Bger            1 MALSDADVQKQIKHMMAFIEQEANEKAEEIDAKAEEEFNIEKGRLVQQQRLKIMEYYEKKEKQVELLKKIQSSNMLNQARLKALKVREDHVRNVLDEARRRLGEVTRNPSKYGQILQSLILQGLLQLLELNVTLRVREVDQSAVQPILSA 150
Tanzaniophasma  1 MALSDADVQKQIKHMMAFIEQEANEKAEEIDAKAEEEFNIEKGRLVQQQRLKIMEYYERKEKQVELQKKIQSSNMLNQARLKALKVREDHVRNVLEEARRRLGEVTRNPTRYSQVLQALIVQGLLQLLEPTVVLRVREVDQSLVQSILPA 150
Choo            1 MALSDADVQKQIKHMMAFIEQEANEKAEEIDAKAEEEFNIEKGRLVQQQRLQIMEYFERKEKDVELQKKIQSSNMLNQARLKVLKMREDHVRGVLDEARRRLGEVTRNPTQYSQILKALTVQGLLQFLETAVTLRVRDVDQSLLHGILPA 150
Gbif            1 MALSDFDVQKQIKHMMAFIEQEANEKAEEIDAKAEEEFNIEKGRLVQQQRLKIMEYYERKEKQVELQKKIQSSNYLNLARLKALKVREDHVRTVLDEARRRLGEVTRDTTSYSRLLKLLAGQGLLQLLEPRVLLRVREVDQSLLQDLLPI 150
Tdom            1 MALSDADVKKQIQHMMAFIEQEANEKAEEIDAKAEEEFNIEKGRLVQQQRLKIMEYYDRKEKQVELQKKIQSSNMLNQARLKVLKYQEDHVHNVLDEARRRLGEVTRDPARYSQILQKLIIQSLFQLLEANVILRVRQADIGLIEDAIPV 150
Gryllotalpa     1 MALSDADVQKQIKHMMAFIEQEANEKAEEIDAKAEEEFNIEKGRLVQQQRLKIMEFYERKEKQVELQKKIQSSNMLNQARLKALKVREDHVESVLDDARKRLGEVTRNPQQYKKILEQLVLQGLFQLLEPIVYIRIRQTDSSMIEEILPS 150
Nlug            1 MALSDADVEKQIKHMMAFIEQEANEKAEEIDAKAEEEFNIEKGRLVQHQRLKIMEYYDRKEKQVELQKKIQSSNMLNQARLKALKVREDHVRSVL-KIQKRLGEVTRNPAKYKEVLQYLIVQGLLQLLESNVVLRVREADVSLIEGIVGS 149
Btab            1 MALSDADVQKQIKHMMAFIEQEANEKAEEIDAKAEEEFNIEKGRLVQNQRLKIMEYYERKEKQVELQKKIQSSNMLNQARLKALIVQEDHIKNVLNDARKKLGEITKDPAQYREVFEKLILQALLQIIETNVVLRVKESEVGFVEKLLPK 150
Lmig            1 SALSAYDYNGAIKHMMAFIDQEANEKAEEIDAKAEEEFNIEKGRLVQQQRLKIMEYYERKEKQVELQKKIQSSNMLNQARLKALKVREDHVRGVLEEARKRLGEITKDQSRYSHVLHQLIMQGLLQLLEPNVTVRTREVDQRIVDSILPA 150
Ofas            1 MSLSDADVQKQIKHMMAFIEQEANEKAEEIDQKAEEEFNIEKGRLVQNQRLKIMEYYERKEKQVELQKKIQSSNMLNQARLRALKVREDHVRDVLDEARKQLVHSVKDDDNYKDLLIKLLLQALMQLMESEVIIKTKESDGPIVSSVFPH 150
Hduo            1 MALSDADVMKQIKHMMAFIEQEANEKAEEIDAKAEEEFNIEKGRLVQQQRIKIMEYYDRKEKQVELQKKIQQSNMHNAARLRVLKEREDHVASVLEDARKRRLAEIVKDRNVYKKILQNLIAQGLLQLLESNVILRCRQADLSIVNESLEV 150
Tjap            1 MALSDADVSKQIKHMMAFIDQEANEKAEEIDAKAEEEFNIEKGRLVQQQRLKIMEFYERKEKQVELQKKIQSSNMLNQARLKVLKARDDHVLSVLEDGKKRMLLISQDQAKYAKLLESLIAQGLCQLIEGQVTIRCRQADKTLVENAVTP 150
Ptep            1 MSLSDADVQKQINHMMAFIEQEANEKVEEIDAKAEEEFNIEKGRLVQEQRLKIMEYYDKKEKQVELQRKIQSSNMLNQARLKVLKARDDHIRTVLEEAKKQLHGLTRDQSRYQEVMKNLVLQAVFQLLEKEVSVKCRKQDIPVVESSLKS 150
Mmus            1 MALSDADVQKQIKHMMAFIEQEANEKAEEIDAKAEEEFNIEKGRLVQTQRLKIMEYYEKKEKQIEQQKKIQMSNLMNQARLKVLRARDDLITDLLNEAKQRLSKVVKDTTRYQVLLDGLVLQGLYQLLEPRMIVRCRKQDFPLVKAAVQK 150
Clup            1 MALSDADVQKQIKHMMAFIEQEANEKAEEIDAKAEEEFNIEKGRLVQTQRLKIMEYYEKKEKQIEQQKKIQMSNLMNQARLKVLRARDDLITDLLNEAKQRLSKVVKDTTRYQVLLDGLVLQGLYQLLEPRMIVRCRKQDFPLVKAAVQK 150
Ggal            1 MALSDADVQKQIKHMMAFIEQEANEKAEEIDAKAEEEFNIEKGRLVQTQRLKIMEYYEKKEKQIEQQKKIQMSNLMNQARLRVLKARDDLISDLLNEAKLRLAKVVKDTARYQILLDGLVLQGFYQLLEPRIVVRCRKQDLPMVKAAVQK 150
Xtro            1 MALSDADVQKQIKHMMAFIEQEANEKAEEIDAKAEEEFNIEKGRLVQTQRLKIMEYYEKKEKQIEQQKKIQMSNLMNQARLKVLKARDDHISELLNEARQRLSRVVKDTARYQMLLDGLILQGLFQLLEPRVLIRCRKQDFPLISASVQK 150
Xlae            1 MALSDADVQKQIKHMMAFIEQEANEKAEEIDAKAEEEFNIEKGRLVQTQRLKIMEYYEKKEKQIEQQKKIQMSNLLNQARLKVLKARDDHISDLVNEARQRLARVVKDTARYQMLLDGLILQGLFQLLEPKVVIRCRKQDLPLITASVQK 150
Dmag            1 MALTDSDVQKQIKQMMGFIEQEANEKVEEIDAKAEEEFNIEKGRLVQQQRLKIMEFYERKEKQVELQKKIQSSNLLNQARLKVLQAQQQHIQNLLAESRTRLAKSSADRSNYTRVICDLIIQALFQIMEPVVTIRCRQVDLELVESVLPE 150
Cowc            1 MALNDEEVDKQIKHMVAFINQEANEKATEMNFKAEEEFNIEKGRLVQQEKIKISAMYERKEKQVDIQKKISYSNELNQSRLRILSERDKHIQSIFNETQQQLAGISSNPDKYRKLLEGLLGQAFHQLLEENVTVRARKVDIALVEAAIPA 150
Rirr            1 RALNDDEVLNEMKKMVAFINQEAIEKAREIKVKADEEFNIEKAKLVRQESNNIEATFQRKIKQAEIQKKITQSNMINKTRLKVLQVRQQMLEEVFNETVSQLNKVSQDEVKYRDLLKNLILQGLYQLMDDQVKAIVRKCDKELANNAIQA 150
Gpro            1 RGLNDQEVQQEMNKMTSFIKQEALEKAREIKVKADEEFNIEKAKLVRQETLNIEAFYEKKQKQAEVQKRITQSTHVNKSRLRILQSRQTLLNQLMEEAKQKLFTLSADPKSYQVLLKDLTLQGLYQLMEKDIIIQVRKKDVALLEAVLPA 150
Atha            1 --MNDGDVSRQIQQMVRFIRQEAEEKANEISVSAEEEFNIEKLQLVEAEKKKIRQDYEKKEKQADVRKKIDYSMQLNASRIKVLQAQDDIVNAMKDQAAKDLLNVSRDEYAYKQLLKDLIVQCLLRLKEPSVLLRCREEDLGLVEAVLDD 148
Pdac            1 --MNDADVSKQIQQMVRFIRQEAEEKANEISVSAEEEFNIEKLQLVEAEKRKIRQEYERKEKQVEIRRKIEYSMQLNASRIKVLQAQDDLVNSMKEAASKELLRVSNDSKAYKMLLKGLIVQSLLRLKEPSVLLRCRKVDRKVVESILNE 148
Obra            1 --MNDADVGKQIHQMVRFILQEAEEKASEISVAAEEEFNIEKLQLVESEKRRIRQEYERKVKQVDVGRKIEYSTQLNAARIKVLQAQDGVVGEMKEDAGRGLLRVTKDATAYRKVLKGLVVQSLLRLREPSVVLRCREADRGHVESVLEV 148
```

```
Dmel           151 AVEQYKAQINQNVELFIDEKDFLSADTCGGVELLALNGRIKVPNTLESRLDLISQQLVPEIRNALFGRNVNRKFTD 226
Dyak           151 AVEQYKAQIQQNVDLFIDEKDFLSADTCGGVELLALNGRIKVPNTLESRLELISQQLVPEIRNALFGRNVNRKFTD 226
Dere           151 AVEQYKAAIHQNVDLLIDEKDFLSADTCGGVELLALNGRIKVPNTLESRLELISQQLVPEIRNALFGRNVNRKFTD 226
Dvir           151 AIEQYKAAMKQDVEFYIDEKEYLSANTCGGVELLALNGRIKVPNTLESRLELISQQLVPEIRNALFGRNVNRKFTD 226
Mdom           151 AVEEYKKQMNQGVSVDVDTDNYLPADTCGGIELIALNGRIKVPNTLESRLELISQQLVPEIRNALFGRNVNRKFAD 226
Lcup           151 AIDMYRQQLNQDVNVTIDTDNFLPADTCGGIELIALNGRVKVPNTLESRLELISQQLVPEIRNALFGRNVNRKFAD 226
Bdor           151 TVEEYKQQIGKDVNVHIDTDNHLSADTCGGIELIALNGRLKVPNTLESRLELIAQQLVPEIRNALFGRNVNRKFTD 226
Ccap           151 SLEEYKNKIGKDVNVHIDTDNHLSADTCGGIELIALNGRLKVPNTLESRLELIAQQLVPEIRNALFGRNTNRKFTD 226
Agam           151 AVELYKSKCGRDVVVTLDTENFLPADTTGGVDLLAQSGRIKVANTLESRLELIAQQLVPEIRNALFGRNMNRKFND 226
Bmor           151 AQTDYKNKIKKDVVLKVDTENFLSPDTCGGIELVAARGRIKISNTLESRLELIAQQLLPEIRNALFGRNPNRKFTD 226
Lexo           151 AQQDYKAKIKKDVALKVDSENFLPADTCGGIELIAAKGRIKISNTLESRLELIAQQLLPEIRTALFGRNPNRKFTD 226
Osca           151 AQQDYKAKIKKDVALKVDSENFLPADTCGGIELIAAKGRIKISNTLESRLELIAQQLLPEIRTALFGRNPNRKFTD 226
Harm           151 CQADYKAKIMKDTVLKIDTDNWLSPDTCGGIELVAMKGRIKICNTLESRLELIAQQLLPEIRTALFGRNPNRKFTD 226
Aros           151 IQDTYKEVSGKDTVIKVDKDNFLPVDSCGGVELLAGRGRIKISNTLESRLELIAQQLVPEVRSALFGRNPNRKFTD 226
Amel           151 VQNAYKQITKKDVTIKVDQDNFLPSDSCGGVDLFAAKGRIKVSNTLETRLELIAQQLIPDIRSALFGCNPNRKFID 226
Sinv           151 VQQQYKQKTKKDVTLKIDTDNFLPTESCGGVELLASRGRIKISNTLETRLELIAQQLIPEIRSALFGQNANRKFND 226
Nvit           151 IQQEYKNVAKKEVHLKMDTDNFLPPDSCGGVELLAAKGRIKIVNTLENRLELIAQQLVPEIRTALFGRNANRRFTD 226
Haxy           151 VEEKYKDGCGKEVNLKIDDESHLNQDSTGGVELLAQRGRIKINNTLEARLELIAQQLVPEIRTALFGRNVNRKFTD 226
Hvig           151 TEKKYKDGCGKEISLKIDDQTHLAQDTTGGIELYAQKGRIKINNTLEARLELIAQQLVPEIRTALFGRNTNRKFTD 226
Sjap           151 VQDAYKSGCGKEVILKIDDDTHLAQDTTGGVELFAQKGKIKINNTLESRLELIAQQLVPEIRTALFGRNVNRKFTD 226
Agla           151 VVTKYKDATAKEIVLKLDDESHLSQETTGGIDLLAHKNKIKISNTLEARLELISQQLVPQIRTALFGRNVNRKFTD 226
Achi           151 VVTKYKDATAKEIVLKLDDESHLSQETTGGIDLLAHKNKIKISNTLEARLELISQQLVPQIRTALFGRNVNRKFTD 226
Ldec           151 VTNKYKDATGKDIHLKIDDEIHLSQETTGGIDLLAQKNKIKISNTMEARLELISQQLVPQIRTALFGRNVNRKFTD 226
Dpon           151 VATKYRDVTGRDVNVTLDDAVQLSQDITGGIDLYTRQNKIKISNTLEARLELISQQLIPQIRNALFGRNVNRKFTD 226
Tcas           151 VATKYRDATGKDVHLKIDDESHLPSETTGGVVLYAQKGKIKIDNTLEARLDLIAQQLVPEIRTALFGRNVNRKFTD 226
Xbas           151 VEKKYKDATGVNVALTIEQDSYLPRDTTGGVDLWAKNGRIKINNTLEARLELIAEQLVPEIRTALFGRNPNRRFTD 226
Apla           151 VSKKYKDAIGKEVVLKVDPENNLPADTTGGIELYAHGGRIKISNTLEARLGLISAQLLPQIRTALFGRNANRKFTD 226
Nves           151 VASLYKTAVGKEVNLKIDTENPLGANTTGGIELLAQRGKIKIDNTLEARLELIAQQLVPEIRTALFGRNVNRKFTD 226
Otau           151 TAATYKTNTGKDINLKIDNENPLPADTTGGVDLVAHQGKIRICNTLEARLELIAQQLVPEIRTALFGRNVNRKFND 226
Obor           151 IAAKYKDATGKDIFMKIDLENPLPMDTTGGIELVVHQGRIRINNTLEARLELIAQQLVPQIRAALFGRNVNRKFTD 226
Tdic           151 VAAKYKDATGKDIFMKIDSENPLPADATGGVELVVHQGRIRVNNTLEARLELIAQQLVPQIRTALFGRNVNRKFTD 226
Bger           151 VAGQYKDATGKEVYLRLDSETFLPPDTTGGVELTAQNGRIKITNTLEARLELIAQQLIPEIRGALFGRNPNRKFAD 226
Tanzaniophasma 151 VSEDYKGHTNKDITLKIDTESFLPVDTTGGVELQAQRGRIKITNTLEARLDLISQLLIPEIRGALFGRNPNRKFAD 226
Choo           151 IAQEYKNFTGKDVKLKLDTESFLPVDTTGGVELLAQQARIKISNTLEARLELIAQQLVPEIRNALFGRNPNRKFAE 226
Gbif           151 AMEEYKRATTMDVHLKVDSEYFLPADTTGGVELQAQKGRIKIANTLEARLEMIAQQLVPEVRCALFGRNPNRKFTD 226
Tdom           151 IAAQYTEATGKEVNVKVDKENFLSPDTCGGIELQAQRGKIKINNTLESRMDLIAHQMIPEVRTALFGRNPNRKFAD 226
Gryllotalpa    151 VQEEYESKTCKKVQVKIDNENYLSSETTGGIELYAQQGRIKIVNTMEARLELIAQQLVPEIRSALFGRNVNRKFTD 226
Nlug           150 CAEQYAKMTGKEVVVKLDADNFLAAETCGGVELFARNGRIKIPNTLESRLDLISQQLVPEIRVALFGRNPNRKFSD 225
Btab           151 VTEDYKKISNRDVSLKLDTESFLPAENSGGVEVLAQRGKIKVNNTLEARLSLIAAQLIPQIRVALFGRNPNRKFND 226
Lmig           151 ITQKYKEITGKDISLKVDTEAFLNPEVTGGIELLAQKGRIKIVNTLEARLELIAQQLIPEIRCALFGRNPNRKFAD 226
Ofas           151 VLVKYKEVTGKNLNLKLETGSYLPAESAGGIELIAGRGRIKVVNTLESRLDLISAQLVPQIRTALFGRNANRKFAD 226
Hduo           151 VTQQYKATTGKDVNLKIDKESFLTATITGGVELTAQKGRIKIPNTLEHRLEMLAKQMVPEIRTNLFGQNMNRKFTD 226
Tjap           151 AVNSVKDVIKRDCQVKVDNENFLPANCAGGIELYTQKGKIKVDNTLEARLAMIAQQIMPEIRGALFGANPNRKFTD 226
Ptep           151 ISDAYQQQVEKPIRITIDKDNCLPSDCAGGVEISAQHGKLKINNTLESRLDMISQQLLPEIRQTLFGSNPNRKFND 226
Mmus           151 AIPMYKIATKKDVDVQIDQEAYLPEEIAGGVEIYNGDRKIKVSNTLESRLDLIAQQMMPEVRGALFGANANRKFLD 226
Clup           151 AIPMYKIATKKDVDVQIDQESYLPEEIAGGVEIYNGDRKIKVSNTLESRLDLIAQQMMPEVRGALFGANANRKFLD 226
Ggal           151 SIPIYKNATKRDVDIHIDQDNFLPEEIAGGVEIYNSDGKIKVSNTLESRLDLVAQQMMPEIRVALFGANANRKFLD 226
Xtro           151 SIPTYKAATKQGVEVIIDQENHLPPEIAGGVELYNGDGKIKVSNTLESRLDLIAQQMMPEIRVALFGANTNRKFMD 226
Xlae           151 SIPTYKAATKQGVEVIIDQETHLTPEIAGGVELYNGNGKIKVSNTLESRLDLIAQQMMPEIRVALFGANTNRKFLD 226
Dmag           151 AIAKYSEAMHKPCQINIAKENFLPVDTCGGVELAAFHGRIRVNNTLENRLEMIAGQMLPEMRTKLFNANPNRKFFD 226
Cowc           151 AVKEYAVSTKKTVNVTVDKQNFLAADIAGGVEVSARGGKISVVNTLENRLKLVYKQMLPEIRSSMFGESTTRKFFD 226
Rirr           151 AAKIYKESTKRNINVELDKENNLPDDCAGGVILTSHFGRIRINNTLEERLTLAQEELLPEIRVLLFGPSPNKKHNH 226
Gpro           151 AKKEFEEALKIPSNATIDKNNFLPENSAGGVVLFALGNRIKCSNTLESRLEILGETALPDVRTMLFVVSTSRKFYS 226
Atha           149 AKEEYAGKAKVHAEVAVDTKIFLPPHCSGGVVLASRDGKIVCENTLDARLDVAFRMKLPVIRKSLFGQVTA----- 219
Pdac           149 AKQEYADKAAVHPKITVD-NVYLPPFCSGGVVLASQDGKIVIENTLDARLDVAFRQKLPEIRKRLFGKMEA----- 218
Obra           149 AKKEYAEKAKVNLKILIDGKVYLPPSCSGGVVLASQDGKIVCDNTLDARVEVSFKQKLPEIRKKLFSQKVSQ---- 220
```

**Fig. S9. Multiple alignments of amino acid of V-ATPase E used to phylogenetic analysis.**

Amino acid sequences were obtained from 56 eukaryotes. 226 residues were used to phylogenetic reconstruction. Reconstructed phylogenetic trees show in Fig. S4c. Datasets used in the phylogenetic analysis show in Table S7.

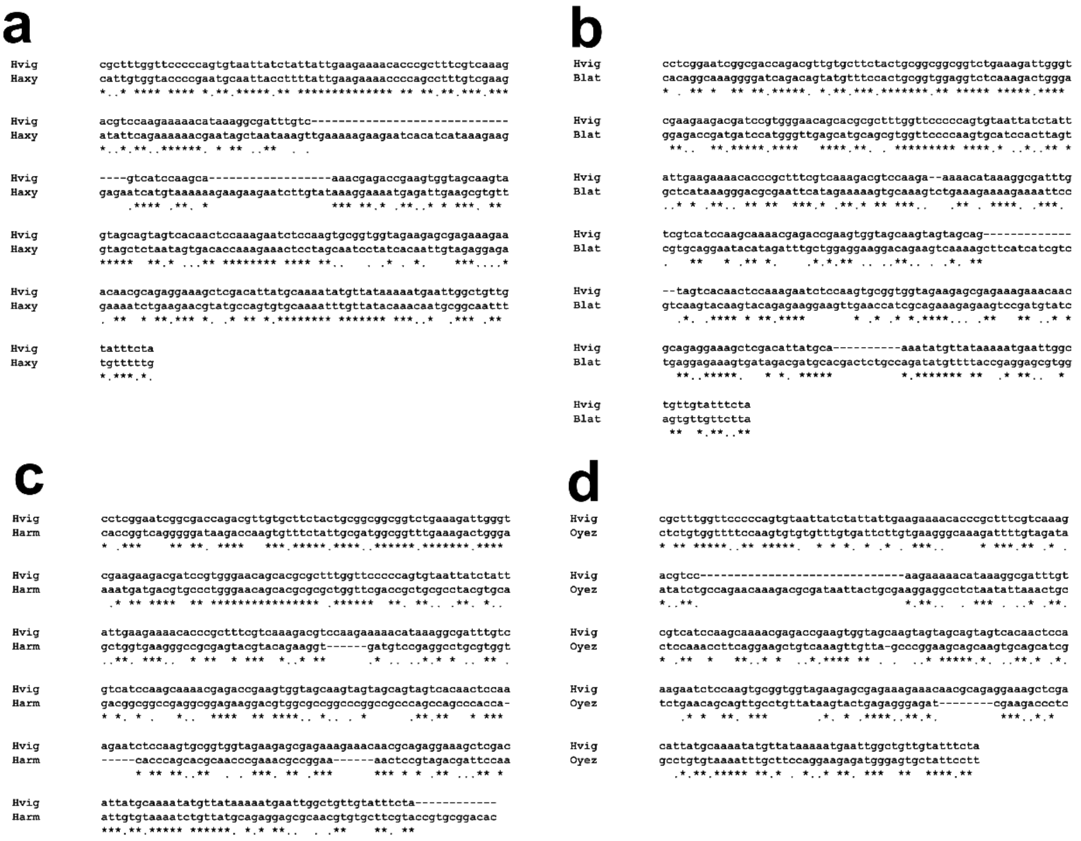


**Fig. S10. Multiple alignments of *diap1* nucleotide sequences used as dsRNA templates between *H. vigintioctopunctata* and other insects.**

In the region of dsRNA, *diap1* did not show consecutive matches over 16 bases between *H. vigintioctopunctata* and each representative insect. (a) Comparison between *H. vigintioctopunctata* (Hvig) and *Harmoni axyridis* (Haxy). (b) Comparison between Hvig and *Blattela lateralis* (Blat). (c) Comparison between Hvig and *Helicoverpa armigera* (Harm). (d) Comparison between Hvig and *Oxya yezoensis* (Oyez). This figure shows in default clustal format, that is, asterisk, colon and period indicate an identical nucleotide site, a strong-conserved site and a weak-conserved site, respectively.

### Supplementary Tables

**Table S1.** Effects of f-RNAi in *Henosepilachna vigintioctopunctata* larvae.

| target of dsRNA | amount (ng) | indivisual | weight (mg) | consumed area ($mm^2$) | |
|---|---|---|---|---|---|
| | | | | 0-24 | 24-48 |
| | | larva 1 | 7.79 | 107.48 | 159.74 |
| | | larva 2 | 9.18 | 113.77 | 142.58 |
| | | larva 3 | 7.21 | 113.65 | 117.78 |
| | | larva 4 | 9.58 | 104.58 | 159.02 |
| *egfp* | 50 | larva 5 | 7.28 | 68.06 | 99.26 |
| | | larva 6 | 6.90 | 64.73 | 150.72 |
| | | larva 7 | 8.16 | 96.48 | 108.22 |
| | | average | 8.01 | 95.54 | 133.90 |
| | | s.d. | 1.03 | 20.78 | 25.09 |
| | | larva 1 | 2.38 | 46.42 | 0.35 |
| | | larva 2 | 2.24 | 39.28 | 0.00 |
| | | larva 3 | 2.73 | 75.41 | 0.36 |
| | | larva 4 | 1.79 | 31.45 | 0.00 |
| *diap1* | 50 | larva 5 | 2.02 | 23.65 | 0.00 |
| | | larva 6 | 2.35 | 47.24 | 0.00 |
| | | larva 7 | 2.74 | 39.22 | 0.35 |
| | | average | 2.32 | 43.24 | 0.15 |
| | | s.d. | 0.35 | 16.41 | 0.19 |
| | | larva 1 | 6.10 | 126.98 | 119.73 |
| | | larva 2 | 8.39 | 150.21 | 137.06 |
| | | larva 3 | 8.01 | 134.93 | 196.36 |
| | | larva 4 | 8.07 | 106.34 | 167.18 |
| *v-ATPase A* | 50 | larva 5 | 6.88 | 121.26 | 99.00 |
| | | larva 6 | 5.30 | 106.51 | 78.72 |
| | | larva 7 | 5.40 | 69.43 | 86.68 |
| | | average | 6.88 | 116.53 | 126.39 |
| | | s.d. | 1.31 | 25.91 | 43.43 |
| | | larva 1 | 2.62 | 29.98 | 18.47 |
| | | larva 2 | 7.75 | 125.49 | 128.54 |
| | | larva 3 | 7.98 | 107.81 | 132.24 |
| | | larva 4 | 8.50 | 181.49 | 158.39 |
| *v-ATPase E* | 50 | larva 5 | 5.71 | 88.51 | 113.18 |
| | | larva 6 | 5.55 | 65.10 | 123.44 |
| | | larva 7 | 5.34 | 57.24 | 74.99 |
| | | average | 6.21 | 93.66 | 107.04 |
| | | s.d. | 2.04 | 50.25 | 46.39 |

**Table S2.** Results of statistical analysis of each f-RNAi experiment. Each *P*-value was calculated by the Welch's *t*-test. Adjusted *P*-value indicates *P*-value corrected by the Holm's method.

| experiment | | time (hrs) | comparison of dsRNA target | | *P*-value | adjusted *P*-value | significance |
|---|---|---|---|---|---|---|---|
| qPCR | | | *egfp* | *diap1* | 0.0124700 | - | yes |
| | | | *egfp* | *v-ATPase A* | 0.0006534 | - | yes |
| | | | *egfp* | *v-ATPase E* | 0.0002460 | - | yes |
| body weight | | | *egfp* | *diap1* | 0.0000015 | 0.0000045 | yes |
| | | | *egfp* | *v-ATPase A* | 0.0972000 | 0.0972000 | no |
| | | | *egfp* | *v-ATPase E* | 0.0667000 | 0.1334000 | no |
| leaf area consumed by larvae | Effects of different target dsRNA | 0-24 | *egfp* | *diap1* | 0.0002522 | 0.0007566 | yes |
| | | | *egfp* | *v-ATPase A* | 0.1216000 | 0.2432000 | no |
| | | | *egfp* | *v-ATPase E* | 0.9295000 | 0.9295000 | no |
| | | 24-48 | *egfp* | *diap1* | 0.0000079 | 0.0000237 | yes |
| | | | *egfp* | *v-ATPase A* | 0.7005000 | 0.7005000 | no |
| | | | *egfp* | *v-ATPase E* | 0.2099000 | 0.4198000 | no |
| | Effects of dsRNA from other insects | 0-24 | *egfp* | Hvig-1 | 0.0335100 | 0.1675500 | no |
| | | | *egfp* | Hvig-2 | 0.0152400 | 0.0914400 | no |
| | | | *egfp* | Haxy | 0.4756000 | 1.0000000 | no |
| | | | *egfp* | Blat | 0.6942000 | 1.0000000 | no |
| | | | *egfp* | Harm | 0.5929000 | 1.0000000 | no |
| | | | *egfp* | Oyez | 0.1155000 | 0.4620000 | no |
| | | 24-48 | *egfp* | Hvig-1 | 0.0004039 | 0.0024234 | yes |
| | | | *egfp* | Hvig-2 | 0.0004039 | 0.0024234 | yes |
| | | | *egfp* | Haxy | 0.5414000 | 1.0000000 | no |
| | | | *egfp* | Blat | 0.0429700 | 0.2148500 | no |
| | | | *egfp* | Harm | 0.1986000 | 0.7944000 | no |
| | | | *egfp* | Oyez | 0.8330000 | 1.0000000 | no |
| | Effects of different dose dsRNA | 0-12 | *egfp* | 0.064 ng dsRNA | 0.4147000 | 1.0000000 | no |
| | | | *egfp* | 0.320 ng dsRNA | 0.2819000 | 0.8457000 | no |
| | | | *egfp* | 1.600 ng dsRNA | 0.4117000 | 1.0000000 | no |
| | | | *egfp* | 8.000 ng dsRNA | 0.0388500 | 0.1554000 | no |
| | | 12-24 | *egfp* | 0.064 ng dsRNA | 0.4634000 | 0.4634000 | no |
| | | | *egfp* | 0.320 ng dsRNA | 0.2776000 | 0.5552000 | no |
| | | | *egfp* | 1.600 ng dsRNA | 0.0237300 | 0.0711900 | no |
| | | | *egfp* | 8.000 ng dsRNA | 0.0104500 | 0.0418000 | yes |
| | | 24-36 | *egfp* | 0.064 ng dsRNA | 0.2605000 | 0.5210000 | no |
| | | | *egfp* | 0.320 ng dsRNA | 0.0441500 | 0.1324500 | no |
| | | | *egfp* | 1.600 ng dsRNA | 0.0025480 | 0.0101920 | yes |
| | | | *egfp* | 8.000 ng dsRNA | 0.0025480 | 0.0101920 | yes |
| | | 36-48 | *egfp* | 0.064 ng dsRNA | 0.7062000 | 0.7062000 | no |
| | | | *egfp* | 0.320 ng dsRNA | 0.0950500 | 0.1901000 | no |
| | | | *egfp* | 1.600 ng dsRNA | 0.0071890 | 0.0215670 | yes |
| | | | *egfp* | 8.000 ng dsRNA | 0.0071720 | 0.0286880 | yes |

**Table S3.** Identity of nucleotide sequences of *diap1*, *v-ATPase A* and *v-ATPase E* among insect species. Coding sequences were used in this analysis. Total length shows the nucleotide length after each multiple alignment of each comparison. Abbreviations in the ‘combination of the comparison’ column are same as Figure 2.

| gene | combination of the comparison | | total length (bp) | No. of identical nucleotide (base) | identity |
|---|---|---|---|---|---|
| *diap1* | Tdom | Lmig | 1208 | 597 | 49.42% |
| | Tdom | Ofas | 1236 | 593 | 47.98% |
| | Tdom | Amel | 1280 | 582 | 45.47% |
| | Tdom | Ldec | 1191 | 541 | 45.42% |
| | Tdom | Haxy | 1218 | 523 | 42.94% |
| | Tdom | Hvig | 1257 | 576 | 45.82% |
| | Tdom | Bmor | 1204 | 524 | 43.52% |
| | Tdom | Dmel | 1410 | 534 | 37.87% |
| | Lmig | Ofas | 1199 | 611 | 50.96% |
| | Lmig | Amel | 1226 | 581 | 47.39% |
| | Lmig | Ldec | 1116 | 562 | 50.36% |
| | Lmig | Haxy | 1170 | 545 | 46.58% |
| | Lmig | Hvig | 1221 | 551 | 45.13% |
| | Lmig | Bmor | 1179 | 492 | 41.73% |
| | Lmig | Dmel | 1337 | 554 | 41.44% |
| | Ofas | Amel | 1241 | 611 | 49.23% |
| | Ofas | Ldec | 1183 | 525 | 44.38% |
| | Ofas | Haxy | 1207 | 549 | 45.48% |
| | Ofas | Hvig | 1249 | 533 | 42.67% |
| | Ofas | Bmor | 1239 | 491 | 39.63% |
| | Ofas | Dmel | 1359 | 537 | 39.51% |
| | Amel | Ldec | 1208 | 569 | 47.10% |
| | Amel | Haxy | 1226 | 575 | 46.90% |
| | Amel | Hvig | 1289 | 573 | 44.45% |
| | Amel | Bmor | 1287 | 512 | 39.78% |
| | Amel | Dmel | 1354 | 550 | 40.62% |
| | Ldec | Haxy | 1114 | 554 | 49.73% |
| | Ldec | Hvig | 1188 | 583 | 49.07% |
| | Ldec | Bmor | 1122 | 488 | 43.49% |
| | Ldec | Dmel | 1338 | 504 | 37.67% |
| | Haxy | Hvig | 1232 | 661 | 53.65% |
| | Haxy | Bmor | 1227 | 457 | 37.25% |
| | Haxy | Dmel | 1319 | 537 | 40.71% |
| | Hvig | Bmor | 1224 | 472 | 38.56% |
| | Hvig | Dmel | 1360 | 537 | 39.49% |
| | Bmor | Dmel | 1320 | 538 | 40.76% |

(continued)

| | | | | | |
|---|---|---|---|---|---|
| *v-ATPase A* | Tdom | Lmig | 3606 | 1374 | 38.10% |
| | Tdom | Ofas | 1887 | 1349 | 71.49% |
| | Tdom | Amel | 1851 | 1386 | 74.88% |
| | Tdom | Ldec | 1845 | 1389 | 75.28% |
| | Tdom | Haxy | 1845 | 1384 | 75.01% |
| | Tdom | Hvig | 1845 | 1397 | 75.72% |
| | Tdom | Bmor | 1854 | 1311 | 70.71% |
| | Tdom | Dmel | 1845 | 1355 | 73.44% |
| | Lmig | Ofas | 3655 | 1374 | 37.59% |
| | Lmig | Amel | 3609 | 1351 | 37.43% |
| | Lmig | Ldec | 3613 | 1347 | 37.28% |
| | Lmig | Haxy | 3613 | 1371 | 37.95% |
| | Lmig | Hvig | 3606 | 1347 | 37.35% |
| | Lmig | Bmor | 3606 | 1379 | 38.24% |
| | Lmig | Dmel | 3606 | 1405 | 38.96% |
| | Ofas | Amel | 1893 | 1341 | 70.84% |
| | Ofas | Ldec | 1884 | 1348 | 71.55% |
| | Ofas | Haxy | 1887 | 1340 | 71.01% |
| | Ofas | Hvig | 1887 | 1350 | 71.54% |
| | Ofas | Bmor | 1896 | 1308 | 68.99% |
| | Ofas | Dmel | 1892 | 1353 | 71.51% |
| | Amel | Ldec | 1851 | 1370 | 74.01% |
| | Amel | Haxy | 1851 | 1377 | 74.39% |
| | Amel | Hvig | 1851 | 1434 | 77.47% |
| | Amel | Bmor | 1854 | 1282 | 69.15% |
| | Amel | Dmel | 1855 | 1283 | 69.16% |
| | Ldec | Haxy | 1845 | 1472 | 79.78% |
| | Ldec | Hvig | 1845 | 1472 | 79.78% |
| | Ldec | Bmor | 1854 | 1360 | 73.35% |
| | Ldec | Dmel | 1845 | 1396 | 75.66% |
| | Haxy | Hvig | 1845 | 1500 | 81.30% |
| | Haxy | Bmor | 1854 | 1388 | 74.87% |
| | Haxy | Dmel | 1845 | 1412 | 76.53% |
| | Hvig | Bmor | 1854 | 1320 | 71.20% |
| | Hvig | Dmel | 1845 | 1343 | 72.79% |
| | Bmor | Dmel | 1854 | 1489 | 80.31% |

(continued)

| | | | | | |
|---|---|---|---|---|---|
| *v-ATPase E* | Tdom | Lmig | 820 | 452 | 55.12% |
| | Tdom | Ofas | 681 | 430 | 63.14% |
| | Tdom | Amel | 681 | 462 | 67.84% |
| | Tdom | Ldec | 681 | 472 | 69.31% |
| | Tdom | Haxy | 681 | 471 | 69.16% |
| | Tdom | Hvig | 681 | 463 | 67.99% |
| | Tdom | Bmor | 681 | 471 | 69.16% |
| | Tdom | Dmel | 681 | 396 | 58.15% |
| | Lmig | Ofas | 819 | 450 | 54.95% |
| | Lmig | Amel | 819 | 445 | 54.33% |
| | Lmig | Ldec | 819 | 441 | 53.85% |
| | Lmig | Haxy | 820 | 465 | 56.71% |
| | Lmig | Hvig | 819 | 439 | 53.60% |
| | Lmig | Bmor | 819 | 450 | 54.95% |
| | Lmig | Dmel | 819 | 398 | 48.60% |
| | Ofas | Amel | 681 | 453 | 66.52% |
| | Ofas | Ldec | 681 | 430 | 63.14% |
| | Ofas | Haxy | 681 | 429 | 63.00% |
| | Ofas | Hvig | 681 | 433 | 63.58% |
| | Ofas | Bmor | 681 | 424 | 62.26% |
| | Ofas | Dmel | 681 | 379 | 55.65% |
| | Amel | Ldec | 681 | 449 | 65.93% |
| | Amel | Haxy | 681 | 477 | 70.04% |
| | Amel | Hvig | 681 | 473 | 69.46% |
| | Amel | Bmor | 681 | 463 | 67.99% |
| | Amel | Dmel | 681 | 387 | 56.83% |
| | Ldec | Haxy | 681 | 508 | 74.60% |
| | Ldec | Hvig | 681 | 491 | 72.10% |
| | Ldec | Bmor | 681 | 476 | 69.90% |
| | Ldec | Dmel | 681 | 415 | 60.94% |
| | Haxy | Hvig | 681 | 533 | 78.27% |
| | Haxy | Bmor | 681 | 473 | 69.46% |
| | Haxy | Dmel | 681 | 403 | 59.18% |
| | Hvig | Bmor | 681 | 455 | 66.81% |
| | Hvig | Dmel | 681 | 388 | 56.98% |
| | Bmor | Dmel | 681 | 443 | 65.05% |

**Table S4**. Effects of *diap1* dsRNA derived from various insects. Hvig: *H. vigintioctopunctata*, Haxy: *Harmonia axyridis*, Blast: *Blattela lateralis*, Harm: *Helicoverpa armigera*, Oyez: *Oxya yezoensis.*

| target of dsRNA | | | *egfp* | leaf area consumed by larvae ($mm^2$) *diap1* Hvig-1 | Hvig-2 | Haxy | Blat | Harm | Oyez |
|---|---|---|---|---|---|---|---|---|---|
| time (hrs) | 0-24 | larva 1 | 109.62 | 44.81 | 28.73 | 50.44 | 144.88 | 141.58 | 79.14 |
| | | larva 2 | 77.48 | 38.48 | 19.71 | 84.78 | 107.41 | 83.33 | 66.02 |
| | | larva 3 | 117.15 | 37.44 | 15.50 | 115.39 | 80.54 | 115.81 | 66.97 |
| | | average | 101.42 | 40.24 | 21.31 | 83.54 | 110.94 | 113.57 | 70.71 |
| | | s.d. | 21.07 | 3.99 | 6.76 | 32.49 | 32.32 | 29.19 | 7.32 |
| | 24-48 | larva 1 | 156.25 | 0.00 | 0.00 | 91.21 | 99.63 | 99.92 | 148.88 |
| | | larva 2 | 148.26 | 0.00 | 0.00 | 181.36 | 126.17 | 185.67 | 163.90 |
| | | larva 3 | 158.48 | 0.00 | 0.00 | 133.29 | 86.02 | 91.23 | 128.91 |
| | | average | 154.33 | 0.00 | 0.00 | 135.29 | 103.94 | 125.61 | 147.23 |
| | | s.d. | 5.37 | 0.00 | 0.00 | 45.11 | 20.42 | 52.20 | 17.55 |

**Table S5.** Effects of amount of *diap1* dsRNA on the eating in *H. vigintioctopunctata*.

| | | | leaf area consumed by larvae (mm2) | | | | |
|---|---|---|---|---|---|---|---|
| target of dsRNA | | | *egfp* | *diap1* | | | |
| amount (ng) | | | | 0.064 | 0.32 | 1.60 | 8.00 |
| time (hrs) | 0-12 | larva 1 | 22.53 | 24.22 | 31.48 | 26.28 | 24.30 |
| | | larva 2 | 40.02 | 37.95 | 17.89 | 39.89 | 20.35 |
| | | larva 3 | 36.53 | 26.52 | 27.14 | 21.87 | 20.40 |
| | | larva 4 | 37.92 | 30.74 | 34.88 | 29.48 | 18.70 |
| | | average | 34.25 | 29.86 | 27.85 | 29.38 | 20.94 |
| | | s.d. | 7.94 | 6.03 | 7.36 | 7.67 | 2.38 |
| | 12-24 | larva 1 | 19.48 | 46.23 | 45.45 | 21.28 | 15.68 |
| | | larva 2 | 49.92 | 43.81 | 27.87 | 14.46 | 2.08 |
| | | larva 3 | 54.71 | 45.02 | 23.86 | 11.92 | 0.41 |
| | | larva 4 | 45.57 | 63.80 | 27.45 | 2.66 | 0.00 |
| | | average | 42.42 | 49.72 | 31.16 | 12.58 | 4.54 |
| | | s.d. | 15.74 | 9.44 | 9.70 | 7.70 | 7.48 |
| | 24-36 | larva 1 | 43.95 | 49.04 | 57.33 | 0.00 | 0.00 |
| | | larva 2 | 57.44 | 31.66 | 8.53 | 0.00 | 0.00 |
| | | larva 3 | 61.79 | 51.86 | 8.80 | 0.00 | 0.00 |
| | | larva 4 | 74.61 | 61.29 | 18.51 | 0.00 | 0.00 |
| | | average | 59.45 | 48.46 | 23.29 | 0.00 | 0.00 |
| | | s.d. | 12.64 | 12.37 | 23.16 | 0.00 | 0.00 |
| | 36-48 | larva 1 | 60.29 | 92.55 | 94.33 | 1.22 | 1.65 |
| | | larva 2 | 78.60 | 74.18 | 7.26 | 0.00 | 0.00 |
| | | larva 3 | 122.62 | 80.23 | 14.72 | 0.00 | 0.00 |
| | | larva 4 | 106.85 | 97.11 | 54.03 | 0.46 | 0.00 |
| | | average | 92.09 | 86.02 | 42.59 | 0.42 | 0.41 |
| | | s.d. | 27.95 | 10.64 | 40.14 | 0.58 | 0.83 |

**Table S6.** Consumed leaf area of *diap1* f-RNA in *H. vigintioctopunctata* until larval death. The blank fields show the death of individuals and unmeasured.

| dsRNA | dose of dsRNA | indivisual | consumed leaf area ($mm^2$) | | | | | |
|---|---|---|---|---|---|---|---|---|
| | | | day1 | day2 | day3 | day4 | day5 | day6 |
| *diap1* | 8 ng | larva1 | 15.84 | 0.00 | 0.00 | 0.00 | 0.00 | 0.00 |
| | | larva2 | 11.45 | 0.00 | 0.00 | 0.00 | | |
| | | larva3 | 17.48 | 0.00 | 0.00 | 0.00 | 0.00 | 0.00 |
| | | larva4 | 12.67 | 0.00 | 0.00 | 0.00 | | |
| | | larva5 | 19.62 | 0.00 | 0.00 | 0.38 | 0.00 | |
| | | larva6 | 22.50 | 0.00 | 0.00 | 0.00 | 0.00 | 0.00 |
| | | larva7 | 15.37 | 0.00 | 0.00 | 0.00 | 0.00 | 0.00 |
| | | average | 16.42 | 0.00 | 0.00 | 0.05 | 0.00 | 0.00 |
| | | s.d. | 3.56 | 0.00 | 0.00 | 0.13 | 0.00 | 0.00 |
| | 50 ng | larva1 | 6.47 | 0.00 | 0.00 | | | |
| | | larva2 | 4.49 | 0.00 | 0.00 | 0.00 | 0.00 | |
| | | larva3 | 28.03 | 0.00 | 0.00 | 0.00 | 0.00 | |
| | | larva4 | 16.92 | 0.23 | 0.00 | 0.00 | | |
| | | larva5 | 13.92 | 0.00 | 0.00 | 0.00 | | |
| | | larva6 | 4.31 | 0.00 | 0.00 | | | |
| | | larva7 | 31.46 | 0.00 | 0.00 | | | |
| | | average | 15.09 | 0.03 | 0.00 | 0.00 | 0.00 | |
| | | s.d. | 10.30 | 0.08 | 0.00 | 0.00 | 0.00 | |

**Table S7.** Data sets for phylogenetic analysis of Diap1, V-ATPase A and V-ATPase E. The multiple alignments and reconstructed phylogeny are shown in Fig. S7-S9 and S4, respectively.

|  | gene | Accessiion No. | data source | taxa1 | taxa2 | taxa3 | species | index |
|---|---|---|---|---|---|---|---|---|
|  | BIR | Q90660 | Swissprot | Metazoa | Vertebrate | Aves | *Gallus gallus* | Ggal_BIR |
|  | BIR7A | Q8JHV9 | Swissprot | Metazoa | Vertebrate | Amphibia | *Xenopus laevis* | Xlae_BIR7A |
|  | BIR7B | A9ULZ2 | Swissprot | Metazoa | Vertebrate | Amphibia | *Xenopus laevis* | Xlae_BIR7B |
|  | BIR7C | A9JTP3 | Swissprot | Metazoa | Vertebrate | Amphibia | *Xenopus tropicalis* | Xtrp_BIR7C |
|  | BIRC1 | Q13075 | Swissprot | Metazoa | Vertebrate | Mammalia | *Homo sapiens* | Hsap_BIRC1 |
|  | BIRC2 | Q13490 | Swissprot | Metazoa | Vertebrate | Mammalia | *Homo sapiens* | Hsap_BIRC2 |
|  | BIRC2 | Q62210 | Swissprot | Metazoa | Vertebrate | Mammalia | *Mus musculus* | Mmus_BIRC2 |
|  | BIRC3 | O08863 | Swissprot | Metazoa | Vertebrate | Mammalia | *Mus musculus* | Mmus_BIRC3 |
|  | BIRC3 | Q13489 | Swissprot | Metazoa | Vertebrate | Mammalia | *Homo sapiens* | Hsap_BIRC3 |
|  | BIRC3 | A1E2V0 | Swissprot | Metazoa | Vertebrate | Mammalia | *Canis lupus* | Clup_BIRC3 |
|  | Xiap | Q9R0I6 | Swissprot | Metazoa | Vertebrate | Mammalia | *Rattus norvegicus* | Rnor_Xiap |
|  | Xiap | Q60989 | Swissprot | Metazoa | Vertebrate | Mammalia | *Mus musculus* | Mmus_Xiap |
|  | Xiap | P98170 | Swissprot | Metazoa | Vertebrate | Mammalia | *Homo sapiens* | Hsap_Xiap |
|  | Xiap | A5D8Q0 | Swissprot | Metazoa | Vertebrate | Amphibia | *Xenopus laevis* | Xlae_Xiap |
|  | Xiap | Q5BKL8 | Swissprot | Metazoa | Vertebrate | Amphibia | *Xenopus tropicalis* | Xtro_Xiap |
|  | Diap | GFPE01022568.1 | NCBI | Metazoa | Hexapoda | Collembola | *Holacanthella duospinosa* | Hdeo_Diap |
|  | Diap | GAXJ02018588.1 | NCBI | Metazoa | Hexapoda | Diplura | *Occasjapyx japonicus* | Ojap_Diap |
|  | Diap2 | T1JFM1 | Swissprot | Metazoa | Myriapoda | Chilopoda | *Strigamia maritima* | Smar_Diap2 |
|  | Diap2 | K7IZE2 | Swissprot | Metazoa | Hexapoda | Hymenoptera | *Nasonia vitripennis* | Nvit_Diap2 |
|  | Diap2 | E9HP27 | Swissprot | Metazoa | Crustacea | Branchiopoda | *Daphnia pulex* | Dpul_Diap2 |
|  | Diap2 | A0A0K2SWB7 | Swissprot | Metazoa | Crustacea | Copepoda | *Lepeophtheirus salmonis* | Lsal_Diap2 |
|  | Diap2 | Q24307 | Swissprot | Metazoa | Hexapoda | Diptera | *Drosophila melanogaster* | Dmel_Diap2 |
|  | Diap2 | GASN02050070.1 | NCBI | Metazoa | Hexapoda | Zygentoma | *Thermobia domestica* | Tdom_Diap2 |
|  | Diap2 | GGNV01253976.1 | NCBI | Metazoa | Hexapoda | Coleoptera | *Leptinotarsa decemlineata* | Ldec_Diap2 |
|  | Diap1 | Q24306 | Swissprot | Metazoa | Hexapoda | Diptera | *Drosophila melanogaster* | Dmel_Diap1 |
|  | Diap1 | B4QLT4 | Swissprot | Metazoa | Hexapoda | Diptera | *Drosophila simulans* | Dsim_Diap1 |
|  | Diap1 | B3NDR0 | Swissprot | Metazoa | Hexapoda | Diptera | *Drosophila erecta* | Dere_Diap1 |
|  | Diap1 | B4ITU5 | Swissprot | Metazoa | Hexapoda | Diptera | *Drosophila yakuba* | Dyak_Diap1 |
|  | Diap1 | B4LGN4 | Swissprot | Metazoa | Hexapoda | Diptera | *Drosophila virilis* | Dvir_Diap1 |
|  | Diap1 | A0A0L0CRF7 | Swissprot | Metazoa | Hexapoda | Diptera | *Lucilia cuprina* | Lcup_Diap1 |
|  | Diap1 | Q16WV1 | Swissprot | Metazoa | Hexapoda | Diptera | *Aedes aegypti* | Aaeg_Diap1 |
|  | Diap1 | B0W3T6 | Swissprot | Metazoa | Hexapoda | Diptera | *Culex quinquefasciatus* | Cqui_Diap1 |
|  | Diap1 | Q7QJ55 | Swissprot | Metazoa | Hexapoda | Diptera | *Anopheles gambiae* | Agam_Diap1 |
| Iap | Diap1 | P41437 | Swissprot | Metazoa | Hexapoda | Lepidoptera | *Orgyia pseudotsugata* | Opse_Diap1 |
|  | Diap1 | MCINX007844-RA | i5k | Metazoa | Hexapoda | Lepidoptera | *Melitaea cinxia* | Mcin_Diap1 |
|  | Diap1 | Q968T8 | Swissprot | Metazoa | Hexapoda | Lepidoptera | *Bombyx mori* | Bmor_Diap1 |
|  | Diap1 | Znev_06782 | i5k | Metazoa | Hexapoda | Isoptera | *Zooterpomosis nevadaensis* | Znev_Diap1 |
|  | Diap1 | GASN02052838.1 | NCBI | Metazoa | Hexapoda | Zygentoma | *Thermobia domestica* | Tdom_Diap1 |
|  | Diap1 | OFAS004454-RA | i5k | Metazoa | Hexapoda | Hemiptera | *Oncopeltus fasciatus* | Ofas_Diap1 |
|  | Diap1 | T1HCP5 | i5k | Metazoa | Hexapoda | Hemiptera | *Rhodnius prolixus* | Rpro_Diap1 |
|  | Diap1 | A0A088A7X5 | Swissprot | Metazoa | Hexapoda | Hymenoptera | *Apis melifera* | Amel_Diap1 |
|  | Diap1 | XM_011166876.1 | NCBI | Metazoa | Hexapoda | Hymenoptera | *Solenopsis invicta* | Sinv_Diap1 |
|  | Diap1 | KQ976424.1 | NCBI | Metazoa | Hexapoda | Hymenoptera | *Atta colombica* | Acol_Diap1 |
|  | Diap1 | - | this study | Metazoa | Hexapoda | Coleoptera | *Henosepilachna vigintioctopunctata* | Hvig_Diap1 |
|  | Diap1 | "ERR1309559" | NCBI | Metazoa | Hexapoda | Coleoptera | *Harmonia axyridis* | Haxy_Diap1 |
|  | Diap1 | A0A1Y1KCG0 | NCBI | Metazoa | Hexapoda | Coleoptera | *Photinus pyralis* | Ppyr_Diap1 |
|  | Diap1 | V5GHQ7 | NCBI | Metazoa | Hexapoda | Coleoptera | *Anoplophora glabripennis* | Agla_Diap1 |
|  | Diap1 | GAZB02077490.1 | NCBI | Metazoa | Hexapoda | Coleoptera | *Lepicerus sp.* | Lepicerus_Diap1 |
|  | Diap1 | GAUY02016224.1 | NCBI | Metazoa | Hexapoda | Coleoptera | *Gyrinus marinus* | Gmar_Diap1 |
|  | Diap1 | GFUZ01020644.1 | NCBI | Metazoa | Hexapoda | Coleoptera | *Amphizoa insolens* | Ains_Diap1 |
|  | Diap1 | GDMY01023426.1 | NCBI | Metazoa | Hexapoda | Coleoptera | *Dryops sp.* | Dryopus_Diap1 |
|  | Diap1 | GDOQ01027193.1 | NCBI | Metazoa | Hexapoda | Coleoptera | *Micromalthus debilis* | Mdeb_Diap1 |
|  | Diap1 | GDNM01013720.1 | NCBI | Metazoa | Hexapoda | Coleoptera | *Xylobiops basilaris* | Xbas_Diap1 |
|  | Diap1 | GDNJ01035765.1 | NCBI | Metazoa | Hexapoda | Coleoptera | *Heterochelus sp.* | Heterochelus_Diap1 |
|  | Diap1 | GBHN01000409.1 | NCBI | Metazoa | Hexapoda | Coleoptera | *Colaphellus bowringi* | Cbow_Diap1 |
|  | Diap1 | GDPR01024911.1 | NCBI | Metazoa | Hexapoda | Coleoptera | *Hydrochara caraboides* | Hcar_Diap1 |
|  | Diap1 | GDLJ01018392.1 | NCBI | Metazoa | Hexapoda | Coleoptera | *Anorus arizonicus* | Aari_Diap1 |
|  | Diap1 | GDNW01017804.1 | NCBI | Metazoa | Hexapoda | Coleoptera | *Cucujus clavipes* | Ccla_Diap1 |
|  | Diap1 | GDLI01019913.1 | NCBI | Metazoa | Hexapoda | Coleoptera | *Hypocaccus fitchi* | Hfit_Diap1 |
|  | Diap1 | GEUD01096124.1 | NCBI | Metazoa | Hexapoda | Coleoptera | *Callosobruchus maculatus* | Cmac_Diap1 |
|  | Diap1 | GATW02015597.1 | NCBI | Metazoa | Hexapoda | Coleoptera | *Aleochara curtula* | Acur_Diap1 |
|  | Diap1 | GGNV01202648.1 | NCBI | Metazoa | Hexapoda | Coleoptera | *Leptinotarsa decemlineata* | Ldec_Diap1 |
|  | Diap1 | GAQW01003002.1 | NCBI | Metazoa | Hexapoda | Coleoptera | *Onthophagus nigriventris* | Onig_Diap1 |
|  | Diap1 | GDAR01021016.1 | NCBI | Metazoa | Hexapoda | Coleoptera | *Dendroctonus ponderosae* | Dpon_Diap1 |
|  | Diap1 | IADJ01048613.1 | NCBI | Metazoa | Hexapoda | Coleoptera | *Trypoxylus dichotomus* | Tdic_Diap1 |
|  | Diap1 | GGAA01019025.1 | NCBI | Metazoa | Hexapoda | Coleoptera | *Nicrophorus orbicollis* | Norb_Diap1 |
|  | Diap1 | GDPC01012259.1 | NCBI | Metazoa | Hexapoda | Coleoptera | *Thanasimus formicarius* | Tfor_Diap1 |
|  | Diap1 | JAMg_model_7561 | i5k | Metazoa | Hexapoda | Orthoptera | *Locusta migratoria* | Lmig_Diap1 |

(continued)

| | | | | | | | |
|---|---|---|---|---|---|---|---|
| | XP_011399845.1 | NCBI | Chlorophyta | Trebouxiophyc | Chlorellales | *Auxenochlorella protothecoides* | Apro |
| | XP_020581168.1 | NCBI | Streptophyta | Embryophyta | Tracheophyta | *Phalaenopsis equestris* | Pequ |
| | XP_006646955.1 | NCBI | Streptophyta | Embryophyta | Tracheophyta | *Oryza brachyantha* | Obra |
| | XP_008789620.1 | NCBI | Streptophyta | Embryophyta | Tracheophyta | *Phoenix dactylifera* | Pdac |
| | KXS15095.1 | NCBI | Fungi | Chytridiomycot | Monoblepharidomycetes | *Gonapodya prolifera* | Gpro |
| | PKY46007.1 | NCBI | Fungi | Mucoromycota | Glomeromycotina | *Rhizophagus irregularis* | Rirr |
| | KJE91442.1 | NCBI | Holozoa | Filasterea | Ministeriida | *Capsaspora owczarzaki* | Cowc |
| | NP_001007512.1 | NCBI | Metazoa | Vertebrate | Amphibia | *Xenpus laevis* | Xtro |
| | XP_018104475.1 | NCBI | Metazoa | Vertebrate | Amphibia | *Xenpus laevis* | Xlae |
| | XP_015149949.1 | NCBI | Metazoa | Vertebrate | Avis | *Gallus gallus* | Ggal |
| | NP_031534.2 | NCBI | Metazoa | Vertebrate | Mammalia | *Mus musculus* | Mmus |
| | XP_545103.1 | NCBI | Metazoa | Vertebrate | Mammalia | *Canis lupus* | Clup |
| | T1II23 | swissprot | Metazoa | Myriapoda | Chilopoda | *Strigamia maritima* | Smar |
| | EFX90349.1 | NCBI | Metazoa | Coleoptera | Branchiopoda | *Daphnia pulex* | Dpul |
| | GDIP01149480.1 | NCBI | Metazoa | Crustacea | Branchiopoda | *Daphnia magna* | Dmag |
| | GGQQ01154761.1 | NCBI | Metazoa | Crustacea | Malacostraca | *Oratosquilla oratoria* | Oora |
| | GASN02050071.1 | NCBI | Metazoa | Hexapoda | Zygentoma | *Thermobia domestica* | Tdom |
| | GAUK02036341.1 | NCBI | Metazoa | Hexapoda | Ephemeroptera | *Ephemera danica* | Edan |
| | GAXA02037250.1 | NCBI | Metazoa | Hexapoda | Ephemeroptera | *Isonychia bicolor* | Ibic |
| | JAMg_model_12597 | NCBI | Metazoa | Hexapoda | Orthoptera | *Locusta migratoria* | Lmig |
| | GFMG01304196.1 | NCBI | Metazoa | Hexapoda | Orthoptera | *Gryllus bimaculatus* | Gbim |
| | GBID01003875.1 | NCBI | Metazoa | Hexapoda | Blattodea | *Blattella germanica* | Bger |
| | OFAS000728 | NCBI | Metazoa | Hexapoda | Hemiptera | *Oncopeltus fasciatus* | Ofas |
| | GFXM01028507.1 | NCBI | Metazoa | Hexapoda | Hemiptera | *Bemisia tabaci* | Btab |
| | IACV01118816.1 | NCBI | Metazoa | Hexapoda | Hemiptera | *Nilaparvata lugens* | Nlup |
| | GDAW01022047.1 | NCBI | Metazoa | Hexapoda | Hemiptera | *Lygus lineolaris* | Llin |
| | GAYE02020252.1 | NCBI | Metazoa | Hexapoda | Thysanoptera | *Frankliniella cephalica* | Fcep |
| | XP_011163633.1 | NCBI | Metazoa | Hexapoda | Hymenoptera | *Solenopsis invicta* | Sinv |
| V-ATPase A | XP_623495.1 | NCBI | Metazoa | Hexapoda | Hymenoptera | *Apis melifera* | Amel |
| | XP_012267080.1 | NCBI | Metazoa | Hexapoda | Hymenoptera | *Athalia rosae* | Aros |
| | XP_001604685.1 | NCBI | Metazoa | Hexapoda | Hymenoptera | *Nasonia vitripennis* | Nvit |
| | XP_023012283.1 | NCBI | Metazoa | Hexapoda | Coleoptera | *Leptinotarsa decemlineata* | Ldec |
| | XP_019764847.1 | NCBI | Metazoa | Hexapoda | Coleoptera | *Dendroctonus ponderosae* | Dpon |
| | XP_018579543.1 | NCBI | Metazoa | Hexapoda | Coleoptera | *Anoplophora glabripennis* | Agla |
| | APB08718.1 | NCBI | Metazoa | Hexapoda | Coleoptera | *Holotrichia parallela* | Hpar |
| | XP_019879320.1 | NCBI | Metazoa | Hexapoda | Coleoptera | *Aethina tumida* | Atum |
| | XP_017776369.1 | NCBI | Metazoa | Hexapoda | Coleoptera | *Nicrophorus vespilloides* | Nves |
| | XP_022904700.1 | NCBI | Metazoa | Hexapoda | Coleoptera | *Onthophagus taurus* | Otau |
| | XP_018326761.1 | NCBI | Metazoa | Hexapoda | Coleoptera | *Agrilus planipennis* | Apla |
| | IADJ01044263.1 | NCBI | Metazoa | Hexapoda | Coleoptera | *Trypoxylus dichotomus* | Tdic |
| | JR472786.1 | NCBI | Metazoa | Hexapoda | Coleoptera | *Rhynchophorus ferrugineus* | Rfer |
| | JU407646.1 | NCBI | Metazoa | Hexapoda | Coleoptera | *Pogonus chalceus* | Pcha |
| | "ERR1309559" | NCBI | Metazoa | Hexapoda | Coleoptera | *Harmonia axyridis* | Haxy |
| | - | this study | Metazoa | Hexapoda | Coleoptera | *Henosepilachna vigintioctopunctata* | Hvig |
| | NP_001091829.1 | NCBI | Metazoa | Hexapoda | Lepidoptera | *Bombyx mori* | Bmor |
| | XP_021181051.1 | NCBI | Metazoa | Hexapoda | Lepidoptera | *Helicovarpa armigera* | Harm |
| | XP_001969693.1 | NCBI | Metazoa | Hexapoda | Diptera | *Drosophila erecta* | Dere |
| | NP_652004.2 | NCBI | Metazoa | Hexapoda | Diptera | *Drosophila melanogaster* | Dmel |
| | AKJ26286.1 | NCBI | Metazoa | Hexapoda | Diptera | *Diabrotica virgifera* | Dvir |
| | XP_001849275.1 | NCBI | Metazoa | Hexapoda | Diptera | *Culex quinquefasciatus* | Cqui |
| | XP_312843.1 | NCBI | Metazoa | Hexapoda | Diptera | *Anopheles gambiae* | Agam |
| | XP_002088521.1 | NCBI | Metazoa | Hexapoda | Diptera | *Drosophila yakuba* | Dyak |
| | XP_011291042.1 | NCBI | Metazoa | Hexapoda | Diptera | *Musca domestica* | Mdom |
| | XP_004533380.1 | NCBI | Metazoa | Hexapoda | Diptera | *Ceratitis capitata* | Ccap |
| | XP_014100281.1 | NCBI | Metazoa | Hexapoda | Diptera | *Bactrocera oleae* | Bole |
| | XP_023299314.1 | NCBI | Metazoa | Hexapoda | Diptera | *Lucilia cuprina* | Lcup |
| | AAB71659.1 | NCBI | Metazoa | Hexapoda | Diptera | *Aedes aegypti* | Aaeg |

(continued)

| | | | | | | | |
|---|---|---|---|---|---|---|---|
| | NP_192853.1 | NCBI | Streptophyta | Embryophyta | Tracheophyta | *Arabidopsis thaliana* | Atha |
| | XP_006654547.1 | NCBI | Streptophyta | Embryophyta | Tracheophyta | *Oryza brachyantha* | Obra |
| | XP_008788110.1 | NCBI | Streptophyta | Embryophyta | Tracheophyta | *Phoenix dactylifera* | Pdac |
| | KXS21230.1 | NCBI | Fungi | Chytridiomycot | Monoblepharidomycetes | *Gonapodya prolifera* | Gpro |
| | XP_025171477.1 | NCBI | Fungi | Mucoromycota | Glomeromycotina | *Rhizophagus irregularis* | Rirr |
| | XP_004347333.1 | NCBI | Holozoa | Filasterea | Ministeriida | *Capsaspora owczarzaki* | Cowc |
| | NP_031536.2 | NCBI | Metazoa | Vertebrate | Mammalia | *Mus musculus* | Mmus |
| | XP_005637441.1 | NCBI | Metazoa | Vertebrate | Mammalia | *Canis lupus* | Clup |
| | NP_001006246.1 | NCBI | Metazoa | Vertebrate | Avis | *Gallus gallus* | Ggal |
| | NP_989123.1 | NCBI | Metazoa | Vertebrate | Amphibia | *Xenopus tropicalis* | Xtro |
| | NP_001079767.1 | NCBI | Metazoa | Vertebrate | Amphibia | *Xenopus larvis* | Xlae |
| | aug3.g14276.t1 | i5k | Metazoa | Chelicerata | Aranea | *Parasteatoda tepidariorum* | Ptep |
| | IABZ01107681.1 | NCBI | Metazoa | Crustacea | Isopoda | *Ligia exotica* | Lexo |
| | GDIQ01037451.1 | NCBI | Metazoa | Crustacea | Branchipoda | *Daphnia magna* | Dmag |
| | GCHA01009383.1 | NCBI | Metazoa | Crustacea | Maxillopoda | *Tigriopus japonicus* | Tjap |
| | GFPE01023016.1 | NCBI | Metazoa | Hexapoda | Collembola | *Holacanthella duospinosa* | Hduo |
| | GASN02045144.1 | NCBI | Metazoa | Hexapoda | Zygentoma | *Thermobia domestica* | Tdom |
| | JAMg_model_17691 | i5k | Metazoa | Hexapoda | Orthoptera | *Locusta migratoria* | Lmig |
| | GAWZ02013861.1 | NCBI | Metazoa | Hexapoda | Orthoptera | *Gryllotalpa sp.* | Gryl |
| | GAXB02007033.1 | NCBI | Metazoa | Hexapoda | Mantophasmatodea | *Tanzaniophasma sp.* | Tanz |
| | GFVY01084750.1 | NCBI | Metazoa | Hexapoda | Phasmatodea | *Clitarchus hookeri* | Choo |
| | GAWP02043902.1 | NCBI | Metazoa | Hexapoda | Grylloblattodea | *Grylloblatta bifratrilecta* | Gbif |
| | BGER005407-RA | i5k | Metazoa | Hexapoda | Blattodea | *Blattela germanica* | Bger |
| | ADJ18242.1 | NCBI | Metazoa | Hexapoda | Hemiptera | *Nilaparvata lugens* | Nlug |
| | GBIJ01015122.1 | NCBI | Metazoa | Hexapoda | Hemiptera | *Bemisia tabaci* | Btab |
| | OFAS009231-RA | i5k | Metazoa | Hexapoda | Hemiptera | *Oncopeltus fasciatus* | Ofas |
| | XP_012263913.1 | NCBI | Metazoa | Hexapoda | Hymenoptera | *Athalia rosae* | Aros |
| V-ATPase E | XP_003424562.1 | NCBI | Metazoa | Hexapoda | Hymenoptera | *Nasonia vitripennis* | Nvit |
| | XP_011158447.1 | NCBI | Metazoa | Hexapoda | Hymenoptera | *Solenopsis invicta* | Sinv |
| | "ERR1309559" | NCBI | Metazoa | Hexapoda | Coleoptera | *Harmonia axyridis* | Haxy |
| | - | this study | Metazoa | Hexapoda | Coleoptera | *Henosepilachna vigintioctopunctata* | Hvig |
| | XP_018569198.1 | NCBI | Metazoa | Hexapoda | Coleoptera | *Anoplophora glabripennis* | Agla |
| | XP_023028494.1 | NCBI | Metazoa | Hexapoda | Coleoptera | *Leptinotarsa decemlineata* | Ldec |
| | XP_970621.1 | NCBI | Metazoa | Hexapoda | Coleoptera | *Tribolium castanum* | Tcas |
| | XP_018320654.1 | NCBI | Metazoa | Hexapoda | Coleoptera | *Agrilus planipennis* | Apla |
| | XP_017783515.1 | NCBI | Metazoa | Hexapoda | Coleoptera | *Nicrophorus vespilloides* | Nves |
| | XP_022912201.1 | NCBI | Metazoa | Hexapoda | Coleoptera | *Onthophagus taurus* | Otau |
| | KRT79209.1 | NCBI | Metazoa | Hexapoda | Coleoptera | *Oryctes borbonicus* | Obor |
| | XP_019762642.1 | NCBI | Metazoa | Hexapoda | Coleoptera | *Dendroctonus ponderosae* | Dpon |
| | GGMU01144561.1 | NCBI | Metazoa | Hexapoda | Coleoptera | *Serangium japonicum* | Sjap |
| | GFXC01017540.1 | NCBI | Metazoa | Hexapoda | Coleoptera | *Anoplophora chinensis* | Achi |
| | GDNM01008195.1 | NCBI | Metazoa | Hexapoda | Coleoptera | *Xylobiops basilaris* | Xbas |
| | IADJ01034222.1 | NCBI | Metazoa | Hexapoda | Coleoptera | *Trypoxylus dichotomus* | Tdic |
| | GBDM01008134.1 | NCBI | Metazoa | Hexapoda | Coleoptera | *Helicoverpa armigera* | Harm |
| | ABF51412.1 | NCBI | Metazoa | Hexapoda | Lepidoptera | *Bombyx mori* | Bmor |
| | GAHQ01004419.1 | NCBI | Metazoa | Hexapoda | Lepidoptera | *Ostrinia scapulalis* | Osca |
| | NP_524237.1 | NCBI | Metazoa | Hexapoda | Diptera | *Drosophila melanogaster* | Dmel |
| | XP_002098037.1 | NCBI | Metazoa | Hexapoda | Diptera | *Drosophila yakuba* | Dyak |
| | XP_001978953.1 | NCBI | Metazoa | Hexapoda | Diptera | *Drosophila erecta* | Dere |
| | XP_002053651.1 | NCBI | Metazoa | Hexapoda | Diptera | *Drosophila viris* | Dvir |
| | XP_005178098.1 | NCBI | Metazoa | Hexapoda | Diptera | *Musuca domestica* | Mdom |
| | XP_023303251.1 | NCBI | Metazoa | Hexapoda | Diptera | *Lucilia cuprina* | Lcup |
| | XP_011209552.1 | NCBI | Metazoa | Hexapoda | Diptera | *Bactrocera dorsalis* | Bdor |
| | XP_004518208.1 | NCBI | Metazoa | Hexapoda | Diptera | *Ceratitis capitata* | Ccap |
| | XP_312551.1 | NCBI | Metazoa | Hexapoda | Diptera | *Aedes gambiae* | Agam |

**Table S8.** The primers used in this study. Tm indicates the annealing temperature.

| primer name | primer sequence (5'-3') | Tm (°C) | Application |
|---|---|---|---|
| diap1_F1 | CGIGAIGCIGGITTYTWYTA | 45 | degenerate PCR for cloning *diap1* from insects |
| diap1_F2 | GAYKIICCITGGGARSARCAYG | 45 | |
| diap1_R | CAIGYIRYIAIRTGICCRCAIGG | 45 | |
| Hvig_diap1_1 | CTTCGACCCAATCTTTCAGACCGCC | 68 | RACE of diap1 in *H. viginctiopunctata* |
| Hvig_diap1_2 | AAAGCGCGTGCTGTTCCCACGGATC | 68 | |
| Hvig_diap1_3 | GATCCGTGGGAACAGCACGCGCTTT | 68 | |
| Hvig_diap1_4 | GAAACAACGCAGAGGAAAGCTCGAC | 68 | |
| Hvig_diap1_RT-PCR_F | ATTGAATTTGGCAAGCATGAGGACA | 68 | RT-PCR and qRT-PCR |
| Hvig_diap1_RT-PCR_R | CAGTAGTACGTCGAAGTGTCAGTGA | | |
| Hvig_v-ATPase_A_RT-PCR_F | TCTATAAGACTGTGGGCATGTTG | 65 | |
| Hvig_v-ATPase_A_RT-PCR_R | TATTCATGGCCTCCCTGATTACA | | |
| Hvig_v-ATPase_E_RT-PCR_F | AATGTCTTGGAGGAAGCACGTAAAA | 66 | |
| Hvig_v-ATPase_E_RT-PCR_R | TACGAGAACATGGGATTCAAACAAC | | |
| Hvig_rp49_RT-PCR_F | ATAGGGTTAGAAGACGTTTCAAGGG | 64 | |
| Hvig_rp49_RT-PCR_R | CTTCCAATTCTCTGACATTATGCAC | | |
| Hvig_diap1_F | AAGTGCGGTATCAAGGGACA | 66 | 1st amplification of template of dsRNA |
| Hvig_diap1_R | ATCGCCAAAACTGCCACTTT | | |
| Hvig_v-ATPase_A_F | TCTGTTGAGTTGGGTCCTGG | 67 | |
| Hvig_v-ATPase_A_R | CCACGGGCATGTTAGATGTG | | |
| Hvig_v-ATPase_E_F | AGCGATGCGGATGTTCAAAA | 66 | |
| Hvig_v-ATPase_E_R | TGGGCTATCAACTCCAACCT | | |
| Hvig_T7-diap1_F | taatacgactcactatagggCGCAGGCTTTTACTACCTCG | 67 | 2nd amplification of template of dsRNA |
| Hvig_T7-diap1_R | taatacgactcactatagggCACAAGCTACCATATGACCGC | | |
| Hvig_T7-v-ATPase_A_F | taatacgactcactatagggAATGTGCCTGCCCTGTCTAG | 68 | |
| Hvig_T7-v-ATPase_A_R | taatacgactcactatagggCCACAACCAAAAGCTCCAGG | | |
| Hvig_T7-v-ATPase_E_F | taatacgactcactatagggAGCGATGCGGATGTTCAAAA | 66 | |
| Hvig_T7-v-ATPase_E_R | taatacgactcactatagggCCCCAGTCTTTTACGTGCTT | | |
| T7-KS | taatacgactcactatagggAGACCACTCGAGGTCGACGGTATC | 55 | amplification of template of dsRNA from pBluescript vector |
| T7-SK | taatacgactcactatagggAGACCACCGCTCTAGAACTAGTGGATC | 55 | |

**Supplementary Sequences**

The sequences of *diap1*, *v-ATPase A* and *v-ATPase E* in *Henosepilachna vigintioctopunctata* (Hvig) and *H. axyridis* (Haxy). The coding sequence regions are showed by underlines. The target sites of f-RNAi and amplificated sequences by qRT-PCR in *H. vigintioctopunctata* sequences are shaded in grey and yellow, respectively.

```
>Hvig_diap1
GGTAAATGTCATGGTTATGCAAGGAAGTAACCATCCCGTGTTTATGTTTATATTACTAG
TTCGTTTCAATTACTCTAAAATTTGTAGTGGATATAATCAAGATAATTGTTCTGGCTTT
ACGATTTCTGCAACTCATAAGTTTTAAACCTCGCATTCGAAAATGGTTCCACCAGTAGA
AGTATTGTCTTATCCAAGCTCGACTAGGAAGTTTGGTAACGGACTTGATTTGAAGAATA
TGCCCGCAGCAGTAAGTGTGAAGAGAAATATAAATCATACCATAGGAAAAGACACTGGC
GATAATGGTTGTTCCTTTTTGAATCTAACTCCACCCGCAAATTTATTGGCTACGATCGA
GGGACGCCTGAAGACGTACAAAAATTGGCCCAACAAGAATATAGATCCCCAGAAGTTAG
CGGCCGCCGGCTTTTTCTATTCTGGAAAAACTGACATCGTCGAGTGTTTCAAGTGCGGT
ATCAAGGGACACAACTGGTTGTTGAACGACGATCCAATGGAAGATCACAAAAAATGGAA
TAGGAATTGTTCTTTTGTAAGAGAAAACGCACCCGAAGAAAATAACGTCCCACAAACTG
GCACCGGTAGTGATTATTGTGGCAATTTAGACGTCGTAACCTCACAATATACAGTCAGC
GAAGAACCTGGAGATTATTATCGTAACTTGGGTGTGGACATCTCTCCGTTCTTGCAAAC
TGGCGCTAAAACGAATCAGTCCGAGGGTCATAACCTGGAGGGGTTGTTACTAAGAACGA
GGAAGGGCCCAAGTCACCCAGATCAGATCATTTACGAGCGTAGAGTGGCGACATTCGCG
AATTGGCCCAAGTCCTTGAAACAGAAACCCACGGACTTGGCGGCCGCAGGCTTTTACTA
CCTCGGAATCGGCGACCAGACGTTGTGCTTTTACTGCGGCGGCGGTCTGAAAGATTGGG
TCGAAGAAGACGATCCGTGGGAACAGCACGCGCTTTGGTTCCCCCAGTGTAATTATCTA
TTATTGAAGAAAACACCCGCTTTCGTCAAAGACGTCCAAGAAAAACATAAAGGCGATTT
GTCGTCATCCAAGCAAAACGAGACCGAAGTGGTAGCAAGTAGTAGCAGTAGTCACAACT
CCAAAGAATCTCCAAGTGCGGTGGTAGAAGAGCGAGAAAGAAACAACGCAGAGGAAAGC
TCGACATTATGCAAAATATGTTATAAAAATGAATTGGCTGTTGTATTTCTACCTTGCGG
TCATATGGTAGCTTGTGTAGATTGTGCATCAGGATTAAAAGAATGTGCTATTTGCCGTA
AAGAGATCCAAGCGAATGTTCGAGCCTTTTTGTCATAGTTCGCGAACAGTTAACAGTAA
CAGTTACTGCTTGACCACACTCATTTTCAAAGAAGTATGGTCGAATTATGTCTCCAGCT
CAAAATCACCACCAAACAGTGACACGTTGTGGATGCATTTGTGTTTCGACAATTACAGG
TGGGTTCTCAGAACCCTAAAAGTGGCAGTTTTGGCGATTAACAAACCCATCTAGGTGAA
AATGTGCTGCGTCGCTCACGAAGATTTTGCTCGAAATACAGCATTCCCTTTTGATGCCG
TATGTTGCAAACTTTCTTCACTGTGCATGGGCAAAAGGCTTCAGTTGTTGTGGTTATTG
AATTTGGCAAGCATGAGGACACAGATCTTTACTGATTATACGCTGTAGAGAGCTTCTTG
AAATTTGAAATTCTTGTCCACGACGTCGAATTGAGATTCCTGGATTGTCACTGACACTT
CGACGTACTACTGTGATTTTGACATTTGGACGACCATTGTATATGACGTATACAACGGA
TCCTGTCTCCCTGAATTCTTCATCTCTTCACAGTTGACGAAGTTAAAACACTATTCCGA
CAACATTTCGAATGAAATTTTAGAACTGCTCTGCCAAGCTTTCATTATTTTGAAATAT
TGTTCAATAATGAAGTAACGTTGTTCTATCGTGTAATGTTCCATTTTTAATAACCTTAA
ACTGTC
```

>**Hvig_*v-ATPase*_A**

CCGCGGTCAGCTGACTGCCTATTCGTGTGAGTAGAGTTTCCACATCTGTAAAGGTGCTT
TGGCTTTGAAATCTTTTTAGTCTTGTGAGAAATTAGGGTGGTACGAAGACCCGAAGTAT
ATTTTGATTATTTGTTTATATCGGTAAAATCGACAAAGATGTCCAACTTGCCTAAGATA
AGTGACGAGGAGCGAGAAGCAAATTATGGATATGTACATGCAGTATCAGGACCTGTTGT
TACTGCTGAGAAAATGTGTGGATCAGCTATGTACGAGTTGGTCAGAGTTGGATATTTTG
AACTTGTGGGAGAAATTATTCGTCTGGAAGGAGATATGGCCACTATTCAGGTTTACGAA
GAAACATCTGGTGTCACTGTAGGTGATCCTGTACTTCGTACTGGAAAACCATTGTCTGT
TGAGTTGGGTCCTGGTATTATGGGATCAATTTTTGATGGTATTCAGCGTCCATTGAAAG
ATATTAATGACCTTACGCAAAGTATTTATATCCCTAAGGGTGTCAATGTGCCTGCCCTG
TCTAGGACTTCCAAATGGGATTTCAACCCATACAATATTAAATTGGGTTCACATTTGAC
TGGTGGTGACATTTATGGTTTGGTGCATGAAAACACTTTGGTGAAGCAGAAACTAATGC
TACCTCCAAAATCCAAGGGAACAGTTACATATATCGCTGAACCAGGAAGTTATACTGTT
GATGATATTGTTTTGGAAACTGAATTTGATGGTGAACGTTCAAAATATACCATGTTACA
AGTATGGCCCGTACGTCAACCTCGACCTGTCAGTGAGAAATTGCCAGCTAATCATCCTC
TGCTTACAGGACAGAGAGTTTTGGATTCTCTTTTCCCTTGTGTACAAGGTGGTACAACT
GCTATTCCTGGAGCTTTTGGTTGTGGTAAAACTGTAATCTCCCAGTCTTTATCCAAATA
TTCAAACTCTGATGTCATTGTTTACGTAGGATGTGGAGAAAGAGGAAACGAGATGTCTG
AAGTACTTCGTGATTTCCCTGAATTAACTGTCGAAATTGAAGGTCAAACTGAATCGATC
ATGAAACGTACCACGCTAGTCGCCAACACATCTAACATGCCCGTGGCAGCTCGTGAAGC
CTCTATCTATACTGGTATCACACTCTCTGAATACTTCAGGGATATGGGATACAATGTAT
CTATGATGGCTGATTCAACTTCACGTTGGGCAGAAGCCTTGAGAGAAATTTCTGGACGT
TTAGCTGAAATGCCTGCTGATTCAGGTTACCCAGCCTACTTGGGTGCTCGACTTGCTTC
CTTCTATGAACGTGCAGGTAGAGTCAAGTGCTTAGGTAATCCCGATCGTGAAGGTTCAG
TATCTATTGTGGGTGCAGTATCTCCACCTGGTGGTGATTTCTCCGACCCTGTTACATCT
GCTACCCTTGGTATTGTACAAGTGTTTTGGGGTTTAGATAAAAAATTGGCACAAAGAAA
GCATTTCCCTTCCATCAATTGGTTAATTTCTTACTCAAAGTACACAAGAGCATTGGATG
ATTTCTATGATAAAAACTTCCCAGAGTTTGTGCCATTGAGAACCAAAGTGAAGGAAATT
TTACAGGAAGAAGAGGACTTGTCTGAAATTGTGCAATTAGTTGGTAAAGCATCTTTAGC
AGAAACCGATAAGATTACTCTTGAAGTAGCCAAGCTCTTAAAGGAGGACTTCTTACAGC
AGAACTCATATTCTGCCTATGACCGATTCTGTCCATTCTATAAGACTGTGGGCATGTTG
AAAAACATGATTGGTCTTTATGATATGTCAAGACATGCGGTAGAAACCACAGCTCAATC
TGAAAATAAAATTACATGGAATGTAATCAGGGAGGCCATGAATAATATATTATATCAGC
TGAGCAGTATGAAATTCAAAGACCCTGTTAAAGATGGTGAAGCCAAAATCAAGGCTGAA
TTTGATCAACTTTATGAAGATATACAGCAAGCTTTCAGAAACTTAGAAGATTAAAAAGA
TATTTGAATAGTTGACATAATTCCTAAAAGCCAGTTTGGTAAATGGATTTTATCAAAAT
GGCCATACTAAATAAAAATTTAAAATATTCATACAATGAAATTATCGAAACTTGTCGAT
GAGTGTACAATATTATGTTCATATTAAAATAAGAATAATAATTAGAGTTTAAGCAAACG
TTTCTGGTGAAGGTATTTCCTGTTATTCCTTTTCACCTTGATTTTTCCTTCATTCCAAT
CTAATTCTTAGGTTAACCACAAATCTTTGAGATATTCTCAGTATTATTTCTCCCTTTTA
TTATTTTGAGTGGTCCAATTTTAGAAGATTTGCAAGATTTAGTAAGTGATTACGATATC
AACTTAATTTTGAAGTTTTAATTAATATTAATTTGTAAATGTACAGTTATGAAATACAT
AGAAGTGTGTAGTCTATTATATCTTAAATATAGTTCAATGTTCTGGACAATATGATGCA
ATATATTGTTGTTTAAAAAAAAAAA

>**Hvig_*v-ATPase_E***
TGTGTTTGATGGCTTGTGAACTTGAAATCACTTATTCTATATTATGTCACCTATTTTAG
ATAAGAGAATCACTAGCAAATGATCTATGACTTACTAAATAAAAGAAGAAGATATAATT
ACTCTGGTGGGGCAGTTTGATCATATGATCAATAAAACTTTTCACTTTTGACTTGGTGT
TAAAGTATTAGGGTGTACGGACAAAGTAAATTTATTAACTTCTGAAGATTATATATACT
TCCTAATTAATTTTTTAAACAAAAACTCATTTGGAAATAACCATGGCTCTCAGCGATGC
GGATGTTCAAAAACAGATTAAACATATGATGGCTTTATTGAGCAAGAGGCAAATGAAA
AAGCTGAGGAGATTGATGCCAAAGCTGAGGAAGAATTCAACATTGAAAAAGGACGTTTA
GTTCAACAGCAAAGGTTGAAAATTATGGAATATTATGAAAAGAAAGAAAAGCAAGTCGA
ATTGCAGAAAAAGATTCAATCTTCAAATATGTTAAATCAAGCTCGACTTAAAGTTTTGA
AAGTAAGAGAAGATCATGTTAGAAATGTCTTGGAGGAAGCACGTAAAAGACTGGGGGAT
GTGACTCGAGACCGTAAAAAATATGCTGAAATTTTGGAGTCCCTGATTCTTCAAGGTCT
TTATCAGTTGTTTGAATCCCATGTTCTCGTACGCGTTCGACCTCAAGATCGTGATATCG
TTCATTCCATTATTGCCAATACAGAGAAAAAGTACAAAGATGGATGTGGTAAAGAAATA
AGCTTGAAGATCGACGATCAAACTCATCTTGCCCAAGATACGACGGGAGGCATAGAGCT
GTACGCCCAGAAAGGAAGAATCAAGATCAACAACACCCTTGAAGCTAGGTTGGAGTTGA
TAGCCCAACAGTTGGTTCCAGAAATTCGTACTGCTCTATTTGGACGCAACACTAATAGA
AAATTCACTGATTGAATGAATCATATCATTCGTCAGTTGCAGATGTATTATATATCCAT
TTCTAAGCTATCAGTGAAAATTATTAAAATTATATGTCTCATAATTTACACACTTTGGT
GCAATTTATACCTGTTCATTGTATAAAACTACTTCCATCTTTTTAATAGTAATGAATAT
CCTATTGGGATTTTAGTTTTCAGATATATGGAAATGTATTTGTTAAATTATATAAGGCT
AGAATTGTGTTGTTATAAATCAAATACTATATCCATAATAAAATATAACCTGTTGTACT
TTGTTGAAATATTTCCACCTTCATTATGATTTATATAGTGCAC

>**Haxy_*diap1***
AATCCGTTGAATGTACAGCTACATTCCGCGATTTTCTACTATAGTGGTTTCAAAATATA
AAGTTTTATATTATTATCGTGTTTTTGGAAATATGTGTTCAACAGATTATCTTATACG
ACCAGTCAGAGGTCTACATTACCAAGGTAAGAATCAAGATCTGAAAAACATGAAGTCGT
TACTGCAGCAGTACACTCATTCCTTTGGCAGAGACGTAATAGACAACGGCATAGCACTC
CTCGACGCAAAGTTACCAGTGCACACAACAATAGACCGACTGAAGACCTTCAAAAACTG
GCCAAACAAACGAATCGAACCGAAAAAACTAGCGGACGCAGGCTTTTTCTACTGCGGCC
GAGCTGACATAGTCGAGTGTTTCAAGTGTGGAATCAAGGGTCACAATTGGGTAGAAAAC
GACAGACCCATGGAAGATCACATCAAATGGAACAGTAACTGCCCTTTCGTAAGAGAAAA
CGCCACAGAACAAGACCTATTACGGGTAAGACAGAGCCAGGACATATGCGGCAACATCG
GTATGGAAATTCTACCCAATTCAAGACCCGAGGATGACACCGAAATGCTAATAAACCAG
AACCAAATCATAAACTTGAGGTCAGAGTCCAGTCTGGCAGAATTGGGCATAGAAAAGAC
CAAAGGACCCAGACATCTGGATCAGATAATCTTGGAGACCAGACTGGCAACTTTTGAAA
ACTGGCCTAAAGCAATGAAACAGAAACCCATAGACCTGGCTGCTGCAGGTTTTTACTAT
TTAGGCGTCGGAGACCAGGTCATGTGCTTCTATTGTGGGGGAGGTTTAAAAGACTGGGT
TGAACAAGACGATCCCTGGGAACAACATGCATTGTGGTACCCCGAATGCAATTACCTTT
TATTGAAGAAAACCCCAGCCTTTGTCGAAGATATTCAGAAAAAACGAATAGCTAATAAA
GTGGAAAAAGAAGAAGAATCACATAATAAAGAAGGAGAATCATGTAAAAAAGAAGTAGA
ATCTTGTATAAAGGAAAATGAGATTGAAGCGTGTTGTAGCTCTAATAGTGACACCAAAG
AAACTCCTAGCAATCCTATCACAATTGTAGAGGAGAGAAAATCTGAAGAACGTATGCCA
GTGTGCAAAATTTGTTATACAAACAATGCGGCAATTTTGTTTTTGCCATGTGGACATTT
GGTTTCTTGTGCAGATTGTGCGTCAGCCTTGAAAAAGTGTGCAGTATGCCGTCAGGACA
TTAAGGCAGTGGCGCGGGCATTCCTTTCGTAGTGTATAAATAAAATGTGTATAATATTG
AGAGTCTTATTTTTCACCTTTTTATATCTTCAGTAAAATGTATAAAATCATCAATCCAT

AGAATCATTAAAGAAAAAACCATTGAAATTAACAGTTGATATTGGTTTATAGAAAAGTT
CTAGAGGTAGATATTCTGAAGATTTTAGGGTACAGAAAGTAATCATTACAAAAGTAATG
GAAATATCTTGATAGTGATTAAAATTTATAATTATCAAATCCTAAAGTCAAGAATTTGT
GGTGATTACATATCTTGTAAAGTTTTATATATATGAGGATTACTCTATAAATACAAACA
TTTTTTGTGGAACAGAATGACTACTACTTTGTTTATTGAAATACTGATGTTAATTTTAA
TAAATAATGCCACATA

>**Haxy_*v-ATPase_A***

GATTCAAGCTTTAATCAGTAAAGTCAGCTGACTTTTCGGAACCTTGTGACAATTTCAGT
ACTCAGGGGTTCACGTCTGTAGTTCATATTTTCAGTCTTTCTGGGAAATTGGGCGAGGT
TTTGGTGATCCTGTGGTGTTTTTCATTTTACTAATCTTTCGGTGAAAAACAATACAAAA
TGTCGAACTTACGATTAATAAGGGACGAGGAAAAGGAATCTGAATATGGTTATGTTCAT
GCAGTTTCTGGACCAGTCGTAACTGCTGAAAAAATGTGTGGTTCGGCTATGTACGAACT
TGTAAGAGTCGGTTACTTTGAATTGGTTGGTGAAATTATCCGTCTTGAAGGTGACATGG
CCACAATTCAGGTATATGAAGAAACTTCTGGTGTTACTGTTGGAGATCCCGTTCTACGT
ACTGGTAAACCATTGTCTGTTGAGTTGGGTCCTGGTATTATGGGTTCAATTTTTGACGG
TATCCAGCGTCCTCTGAAAGATATCAATGTTTTAACAGAGAGTATCTACATTCCTAAAG
GTATCAATGTACCTTGTTTGTCCAGGACTGCTAAGTGGGACTTCAATCCTATCAATATC
AAATTGGGATCCCATTTAACCGGTGGAGATATTTATGGTATAGTCCACGAAAACACCTT
GGTGAAACAAAAATTGATGTTGCCTCCAAAATCTAAGGGTACAGTCACATACGTCGCAG
AACCTGGAAGTTATACTGTTGATGATATTGTCTTGGAAACCGAATTCGATGGTGAACGC
TCAAAATATACTATGTTGCAAGTTTGGCCAGTACGTCAACCACGTCCAGTCAGTGAAAA
ATTGCCAGCTAATCATCCTCTACTTACTGGACAGAGAGTTTTGGATTCTCTTTTCCCAT
GTGTCCAGGGAGGAACTACTGCTATACCTGGTGCTTTCGGTTGTGGAAAAACTGTCATC
TCACAATCTCTGTCCAAATATTCCAACTCAGATGTCATCATCTATGTCGGTTGCGGAGA
AAGAGGTAACGAAATGTCTGAGGTACTCAGGGACTTCCCCGAACTGACTGTCGAAATCG
AAGGCCAGACCGAATCCATCATGAAGCGTACCGCCTTGGTGGCCAACACCTCCAACATG
CCTGTCGCCGCCCGTGAGGCTTCCATCTACACCGGTATCACTCTATCCGAGTACTTCAG
GGACATGGGTTACAACGTGTCCATGATGGCCGATTCCACCTCTCGTTGGGCTGAAGCCT
TGAGAGAAATCTCCGGTCGTCTGGCCGAGATGCCTGCCGATTCAGGTTACCCCGCCTAC
CTGGGGGCCCGTCTGGCCTCCTTCTACGAGCGTGCCGGTCGCGTCAAATGTCTGGGTAA
CCCCGATCGAGAGGGCTCCGTGTCCATCGTAGGAGCCGTATCGCCACCTGGTGGTGACT
TCTCAGATCCCGTCACTTCCGCCACTCTGGGTATCGTGCAGGTGTTCTGGGGTTTGGAC
AAGAAGCTGGCCCAACGTAAACATTTCCCCTCCATCAACTGGCTGATTTCCTACTCGAA
GTACACCAGGGCTCTGGACGATTTCTACGACAAGAACTTCGCAGAGTTTGTGCCGTTGA
GGACCAAGGTTAAGGAAATCTTGCAGGAAGAAGAGGACTTGTCCGAAATTGTACAACTG
GTAGGAAAAGCATCACTCGCAGAAACTGACAAGATCACTCTGGAGGTAGCCAAGCTACT
TAAAGAAGATTTCTTGCAACAGAACTCCTATTCATCGTACGACAGATTCTGCCCCTTCT
ACAAGACTGTCGGCATGTTGAAAAACATGATCGGTCTTTACGACATGGCCAGACATGCA
GTAGAAACTACCGCCCAATCCGAAAACAAAATTACATGGAACGTAATCAGGGAAGCGAT
GAACAACATCTTGTACCAACTCAGCAGTATGAAGTTCAAGGATCCCGTGAAAGACGGAG
AGGCTAAAATCAAAGCCGATTTCGATCAGCTTTACGAGGACATACAGCAGGCTTTCAGA
AACTTAGAAGATTAGGTAGACTGACATAAGGTTTAGAACTCATTAAGGCCTCTGACATT
GTCTGGACTCATTTTCATATCCAAGTCAACTTTGTATACGGCCATTATAACACATACAT
ATGTTGATATTTCATTGTAAGTGTACAATACTATGTATATATATGAGGTGGATCAAGAA
TGCATCTGAAACTTACATCAAACGGTCGGTTTCACGCAATAAGAATTTTTTGAACGGAA
TGATTGTTGATTAGCAACAATTGGCAAGACATGGATACATTTGAAGCTGATATTACTTA
GAAATTTTCTGAGATGTAATTTTGGATCACCTTGTCTATTGAATTAGAGTAATTTAGAT

```
TTTAATCAAGATTTAGTGGTGAAGGTTTCAGCTAATTCTGTTCACCTTTTTCCTTCATT
CCAATTATACTTTTAGGGAAATTACAGTTTTTTTTTTGAGATGATTTTCTCAGTATTACC
CGTTACGATGTTTTTTTGTCATATAAATTTTACTTTTTTTTTTTTGAAGTGCTGCATATA
ATTCTTCATATCTATAGTTTCCTATTGCTTTACATTTTAATCCAAGTTTTATTGTACAG
TTATCAATTTTCATATATAATAGTTTAAATTTTGAATGTAGTTGAAATGTTCTATTGTG
AACAATAATTATGCAATATATTGTTGTTTTGAAAAAAAA
```

**>Haxy_*v-ATPase_E***

```
CAACTACTAAATTCCAGCTAAAATAAAAAAGCAGTAATTTATCGTGTTCTGCCTAAGAT
GCAATTTTTATTTAACTAATGGCAACAGTACTTGTCAAGGATACGTCAATTATTGTCAA
AGTGGAATTGAGATCATATGATCAATGAAGACCCTGTTTAGTTAAGTACTTGCTTACAA
GGTTCCTCCCTGTTTTATAAAATCTAATTAGAAAAACTAGAAATTACCAAATTTCTTAA
TTTTTCAAATTAAATACAAATTCTAACACAACCATGGCTCTAAGCGATGCAGATGTTCA
AAAACAGATCAAGCATATGATGGCTTTCATTGAGCAAGAAGCCAATGAAAAAGCTGAAG
AAATTGATGCAAAAGCTGAGGAGGAGTTCAATATTGAAAAGGGCCGTTTGGTTCAACAG
CAGCGATTGAAAATTATGGAATATTATGAGAAAAAGGAGAAACAGGTTGAGCTCCAAAA
GAAAATTCAATCCTCTAATATGCTGAATCAAGCACGACTTAAAGTATTGAAAGTTCGTG
AAGATCATGTGAGGAATGTTTTGGAAGACGCTCGTAAGAGATTGGGAGATGTTACTAGA
GACCAGGGAAGGTACAGAGAGATTCTGGAATCTCTTATTCTTCAAGGTTTATACCAGCT
TTTTGAATCTAATGTTGTTATTCGCGTTCGTCCCCAAGACCGAGATCTTGTCAGGTCAA
TAACTTCAAATGTTGAAGAAAAGTATAAGGATGGATGTGGTAAGGAGGTCAATTTGAAA
ATTGATGATGAGTCCCACTTGAACCAAGATTCGACTGGTGGTGTGGAACTTTTAGCTCA
AAGAGGAAGAATCAAGATCAACAACACCCTGGAAGCTAGATTAGAGCTAATTGCTCAAC
AATTGGTTCCAGAAATCCGTACTGCTCTGTTTGGACGTAACGTAAACAGAAAATTCACC
GATTAAATCCATTCCCAAACCTCAAACACTGTATTATTAATTGAATTATTTTCAATGCA
CAAGAAATAATATAATATTAAATTTATTCTCCATTGTTTACATATAGTTGTGCAATTGA
TCCATGATTGATGTATAAAGCAATTCCATTTTAATAGTATAAAAGTTATACCCCTGTTA
GTCTAATTTTTGGTTTTAGATTACAAAAAATATTTGTGAAGTTCCGTAGATCTAGAATT
TGATTTGTTGTAACATTATTGAAATCGAAGTATAGCCTGTTATATATTATGCTTATTTC
GTTTCCCAATCATACTAAAATAGTAAGTGAAGGAAATGGATCCTATAAAAATACTGAAG
AATACTCTGTAG
```
